# From telomere to telomere: the transcriptional and epigenetic state of human repeat elements

**DOI:** 10.1101/2021.07.12.451456

**Authors:** Savannah J. Hoyt, Jessica M. Storer, Gabrielle A. Hartley, Patrick G. S. Grady, Ariel Gershman, Leonardo G. de Lima, Charles Limouse, Reza Halabian, Luke Wojenski, Matias Rodriguez, Nicolas Altemose, Leighton J. Core, Jennifer L. Gerton, Wojciech Makalowski, Daniel Olson, Jeb Rosen, Arian F. A. Smit, Aaron F. Straight, Mitchell R. Vollger, Travis J. Wheeler, Michael C. Schatz, Evan E. Eichler, Adam M. Phillippy, Winston Timp, Karen H. Miga, Rachel J. O’Neill

**Author notes:** Corresponding author: Rachel J. O’Neill. Authors contributed equally (Jessica M. Storer, Gabrielle A. Hartley, Patrick G. S. Grady).

## Abstract

Mobile elements and highly repetitive genomic regions are potent sources of lineage-specific genomic innovation and fingerprint individual genomes. Comprehensive analyses of large, composite or arrayed repeat elements and those found in more complex regions of the genome require a complete, linear genome assembly. Here we present the first de novo repeat discovery and annotation of a complete human reference genome, T2T-CHM13v1.0. We identified novel satellite arrays, expanded the catalog of variants and families for known repeats and mobile elements, characterized new classes of complex, composite repeats, and provided comprehensive annotations of retroelement transduction events. Utilizing PRO-seq to detect nascent transcription and nanopore sequencing to delineate CpG methylation profiles, we defined the structure of transcriptionally active retroelements in humans, including for the first time those found in centromeres. Together, these data provide expanded insight into the diversity, distribution and evolution of repetitive regions that have shaped the human genome.

## Introduction

In the decades since Barbara McClintock’s seminal discovery that genetic elements can transpose from one genomic location to another (*1*), studies of mobile elements and repeat arrays have shown that eukaryotic genomes are in constant flux. Transposable element (TE) insertions and repeat-mediated structural rearrangements, while most often neutral, can impact gene regulation, create new coding structure, and profoundly affect chromosome stability. Transposition, expansion, and contraction of repeats have supported novel, species-specific genomic innovations (e.g. (*2, 3*)), major evolutionary transitions (*4*), and human and primate-specific adaptations (e.g. (*5*)). Correlatively, TEs and other forms of repetitive DNA, constituting half of the human genome, are the largest contributor to human genetic variation and impact human health (*6*) through deleterious copy number variants (CNVs), structural variants (SVs), insertions, deletions, and alterations to gene transcription and splicing.

One of the major challenges in tracking and understanding repeat structure, function and variation is that recent insertions by transposable elements (TEs), large, complex repeats, and sequences found in tandem arrays have been largely impenetrable to previously available sequencing and assembly technologies. Despite this challenge, the expansion of a species-agnostic repeat database (Dfam)(*7*), extensive manual curation (e.g. (*8*), and the development of improved algorithms for repeat discovery (e.g. (*9, 10*)) have all laid indispensable groundwork underlying our efforts to create and finish a complete map and catalog of the repertoire of human repeats. However, previous assemblies of a reference human genome (e.g. GRCh18, GRCh38) that have been developed over the last two decades still contained large gaps and collapsed repeats (*11*), rendering a comprehensive repeat annotation for the entire human genome impossible.

Capitalizing on recent advances in ultralong sequencing and novel assembly methods, the T2T Consortium has released the first complete human reference genome based on the pseudo-haploid genome of an androgenetic hydatidiform mole (CHM13hTERT cell line, hereafter CHM13) (*12*). This assembly, CHM13v1.0, resulted in the addition of over 200 megabase pairs (Mbp) of DNA and resolution of collapsed and unassembled regions in previous reference genomes, providing an unprecedented resource for studying human genomes. The gap-filled and decompressed regions, representing 8% of the human genome, are dominated by tandemly arrayed repeats (such as in the alpha satellite arrays that are found in higher order repeat arrays (HORs) within centromeres(*13*)) and complex repeats in pericentromeres, subtelomeres, and chromosome arms, particularly the acrocentric chromosomes. CHM13v1.0 supported new annotations for human repetitive sequences residing in the unassembled regions of the previous human genome assembly gaps and provided enhanced resolution of repeat calls genome wide. This effort culminated in our first truly genome-scale assessment of the impact of repeats on the landscape of a human genome and a resource to support future work on the impact of repeats to genome structure, function, diversity and evolution. Given the extent of repetitive content in the human genome (53.9% identified in CHM13 in our study), herein we highlight some key advances to the field that this new resource provides, while simultaneously illustrating the power of combining multiple approaches and tools to enable new discoveries.

Eukaryotic repeats are classified into two main types based on their genomic organization: tandem repeats and interspersed repeats (Fig. 1A)(*7, 14*–*17*). Tandem repeats are further subdivided into satellites and simple repeats; satellites are often further defined by their regional chromosomal distribution (centromeric, for example). With the exception of pseudogenes retroposed from structural RNAs (tRNAs, rRNAs, etc.), interspersed repeats largely refer to TEs, which are divided based on their mechanism of propagation. Class I elements are those that are mobilized through transposase, helicase, or recombinase, and include TEs such as Tc1-Mariner and hAT. Class II elements are spread within genomes via retrotransposition and are further divided into classes based on their autonomy. For example, LINEs and LTR elements encode their own catalyzing enzymes while SINEs are non-autonomous, relying on LINE-encoded proteins for retrotransposition. These varied repeat types constitute a major portion, and in some cases the majority (for example, 85% of wheat genomes (*18*)), of eukaryotic genome sequences. Their varied modes of propagation, from simple insertion events to promoting non-allelic recombination, facilitate genomic diversity, often in bursts of activity followed by periods of neutral evolution. Furthermore, the defense mechanisms that have evolved to counter the deleterious effects of mobilization, such as DNA methylation, can influence the sequence evolution of targeted elements. Repeats represent the nexus of evolutionary forces, the selfishness of mobile elements and the cellular mechanisms marshalled to silence them. The genomic turbulence engendered by repeats makes them the most challenging genomic regions to study, but hard-fought insights from these regions have revealed regulatory and coding domains critical to organismal life histories and human health. A full accounting of repeat domains permitted by a gapless telomere-to-telomere DNA sequence is, therefore, essential to a full understanding of the origins and function of the human genome.

**Figure 1.**
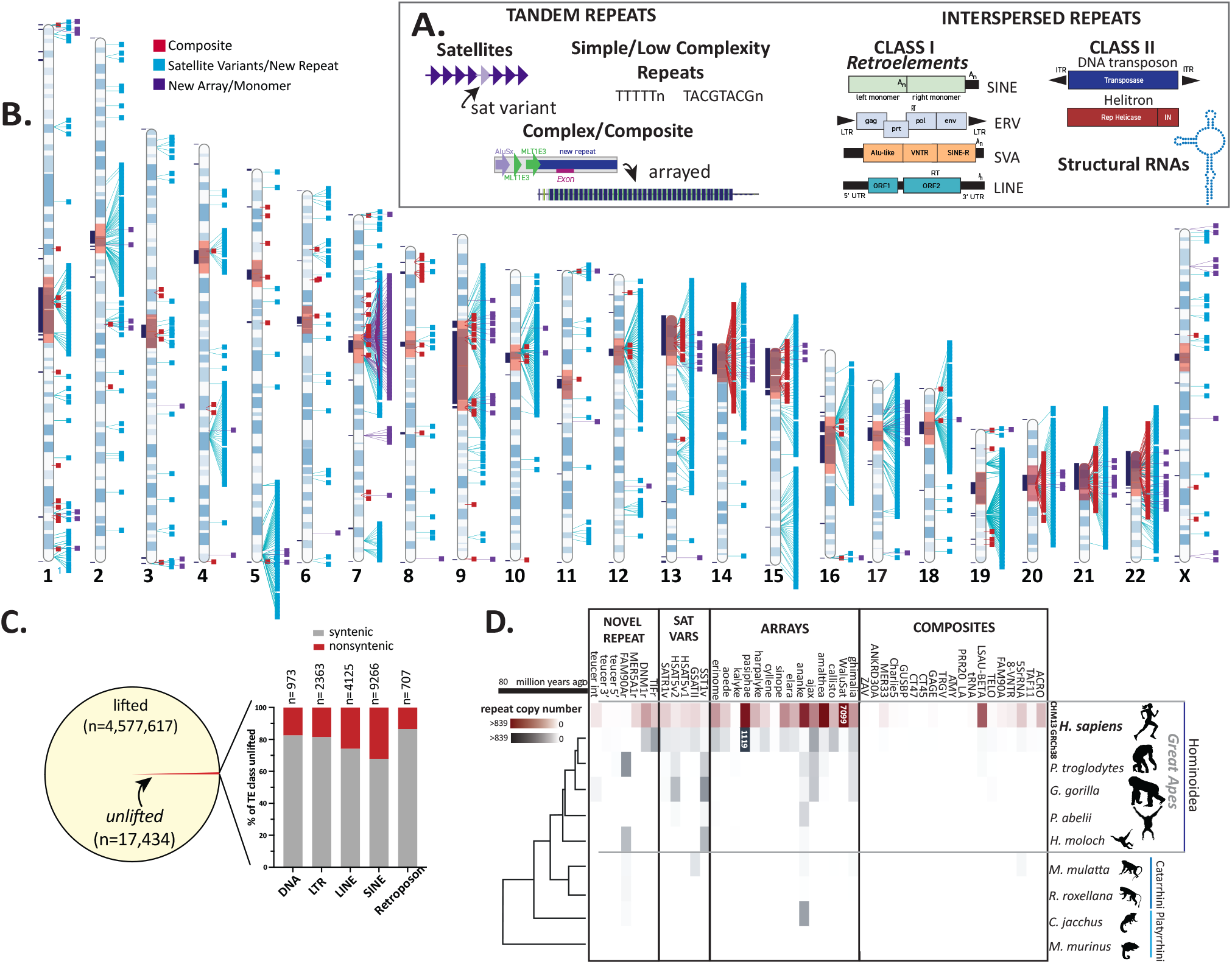
T2T-CHM13 assembly supports new repeat annotation pipeline and the identification of new repeat families. **(A)** Schematic illustrating examples of the two major types of repeats in the human genome: tandem repeats, including satellites, simple and low complexity repeats and composites, and interspersed repeats, including Class I, Class II TEs and structural RNAs. **(B)** Ideogram of CHM13 indicating the locations of newly annotated composite elements (red), satellite variants and novel repeats (aqua), and new arrays or monomers of sequences found within those arrays (purple). Gaps in GRCh38 with no synteny to CHM13 (12) are shown in black boxes to the left of each chromosome, centromere blocks (including centromere transition regions (13)) are indicated in orange. **(C)** The number of TEs lifted and unlifted from CHM13 to GRCh38. (Right) Bar plot showing percentage of TEs by class (DNA, LINE, LTR, Retroposon and SINE) that were unlifted from CHM13 gap filled regions (non-syntenic, red) and syntenic regions (grey); n= number of elements within each class affected. **(D)** Copy numbers of new repeat annotations grouped by novel repeats, variants of known satellites, tandemly arrayed sequences, and composite element (inclusive of subunits) for CHM13 (maroon), GRCh38, and genomes for other primates from the Hominoidea, Catarhini and Platyrrhini lineages (grey). Heatmap scale denotes number of repeats within the array (0-839). Array sizes >839 are indicated within colored blocks. Phylogenetic relationship and millions of years since divergence are indicated on the bottom. Not shown: variants of known centromeric satellites (but see (13)) and the new repeat annotation for an AluJb (25) fragment, which could not reliably be delineated in copy number from other closely related full length AluJb elements.

Analyzing CHM13v1.0 with a novel repeat annotation workflow followed by manual curation, we developed an unprecedented catalog of human repetitive sequences, refined previous annotations, and discovered novel repeats, TE families, repeat variants, and complex, composite repetitive structures. Manual curation added a compendium of new repeats, arrays of non-centromeric satellites, and annotation of complex, composite elements, complement the centromere satellite (*13*) and segmental duplication (*19*) annotations accompanying the CHM13 assembly. We identified high-confidence transduction events derived from L1s, *Alu*, and SVA elements, 295 of which contain coding sequences. As an exemplar, using phylogenetics and statistical modeling we have tracked the fate of a single transduced sequence from a primate ancestor that has led to a repeat expansion encompassing 450 kilobase pairs (Kbp) across four chromosomes, illustrating the impact of different repeat types on one another and on chromosome structure.

Building upon this resource, we assessed RNA polymerase occupancy using precision run-on sequencing (PRO-seq) (*20, 21*) and CpG methylated sites using Oxford Nanopore long read data (*22*), both at single-nucleotide resolution, to derive the first transcriptional and epigenetic annotation for major TE classes across a human genome. This comprehensive approach supported phylogenetic and statistical modelling that exposed differential evolutionary patterns for specific repeat types (retroelements, satellites, macrosatellites, composite elements) that are found genome-wide from those found only in specific chromosomal regions, such as centromeres and telomeres. Our analyses revealed both shared and family-specific epigenetic profiles and RNA polymerase signals, many overlapping repeat annotations of clinical significance.

Using marker assisted mapping and unique k-mers, we assessed the transcriptional landscape of previously inaccessible centromeres. We find that engaged RNA polymerase signal is low, but detectable, across alpha satellites arranged within live centromeric HOR arrays; rather, active transcription is detected in retroelements demarcating boundaries of these arrays. Together, these data reveal the dynamic relationship between transcriptionally active retroelement subclasses and DNA methylation, as well as potential mechanisms for the derivation and evolution of novel repeat families and composite elements. Moreover, the resources developed through this effort provide a foundation for studying RNA polymerase occupancy across the human genome.

## Results

### Comprehensive repeat annotations for a complete human genome

Newly filled gaps and corrections resolved in the T2T-CHM13v1.0 genome assembly added over 200Mbp of DNA, an estimated 75-90% of which are repetitive, necessitating an update of the GRCh38 repeat models and the development of new approaches to produce these models and thus achieve comprehensive repeat annotations. To accomplish this task, we developed a computational pipeline to ascertain new repeat annotations and tandem arrays while reducing false positives from pseudogenes, segmental duplications, and Dfam overlaps (Note S1, S2, Fig. S1). At each step, computational analysis was supplemented by extensive manual curation and polishing.

In total, 49 newly annotated repeat types from *RepeatModeler* were curated, including 27 novel repeats (Fig. 1B, S2) and 22 potentially older TE repeats whose alignment scores precluded classification and were thus set aside for future analyses (Table S1). Among the 27 novel repeats were one novel centromeric satellite (86.6% of bp found within centromere regions defined by (*12, 13*)) and ten repeats further classified into five variants of known satellites (three centromere transition satellites (GSAT, SHAT2, HSAT5), two interstitial satellites (SATR1, SST1)), and five novel repeats. Manual curation identified an additional 13 previously unannotated interstitial satellite arrays (and monomers of the satellites), three new repeats and 19 composite elements consisting of TEs combined with newly annotated repeat “subunits” (16 curated composite subunits) (Fig. S2). In total, 62 new repeat entries were classified and submitted to Dfam as human repeats, with 19 elements added to a new “composite” track for the CHM13 genome browser (Table 1, Table S2).

**Table 1:** Complete genome assembly supported new human repeat annotations. Repeats identified through RepeatModeler and manual curation (RMv2) shown in counts and bp, by new category, for CHM13v1.0 and GRCh38 (excluding the Y chromosome) (Fig. 1B, Table S2).

|  | **CHM13v1.0 RMv2** |  | **GRCh38 RMv2** |  |
| --- | --- | --- | --- | --- |
|  | **count** | **bp** | **count** | **bp** |
| COMPOSITE SUBUNITS | 4,442 | 2,804,927 | 1,162 | 536,979 |
| NEW REPEATS | 1,086 | 807,408 | 783 | 570,985 |
| SATELLITE VARIANTS | 900 | 787,520 | 671 | 591,294 |
| MONOMERS OF ARRAYED SATELLITES | 11,918 | 1,129,174 | 2,651 | 415,508 |
| **ALL NEW REPEAT ENTRIES** | **18,346** | **5,529,029** | **5,267** | **2,114,766** |

Development of an updated repeat library over a complete reference genome yielded new annotations of repeats within CHM13 gap-filled regions, and also provided copy number support required to identify additional, previously unnoticed repeat elements genome-wide (Fig. 1B, Fig. S2). Using this new, CHM13-based repeat library, the CHM13v1.0 assembly was fully annotated for all repeat classes, resulting in a total of 1.64 gigabase pairs (Gbp) of repeat annotations (53.93% of the genome), 207.6Mbp within the 230Mbp of newly assembled genomic sequence (90%), and 5.5Mbp of new repeats identified herein (Table 2, Table S3, Fig. S3). Re-annotation of GRCh38 (without the Y chromosome) with the CHM13 repeat database resulted in annotation of 2,059,378bp of previously uncatalogued repeats (Table 2, S2), while re-annotation of the GRCh38 Y chromosome produced new annotations including 6 composite elements, 7 satellite arrays, and 162 satellite variants/new repeats, totaling 158,604bp in new repeat annotations (Fig. S4, Table S4).

**Table 2.**
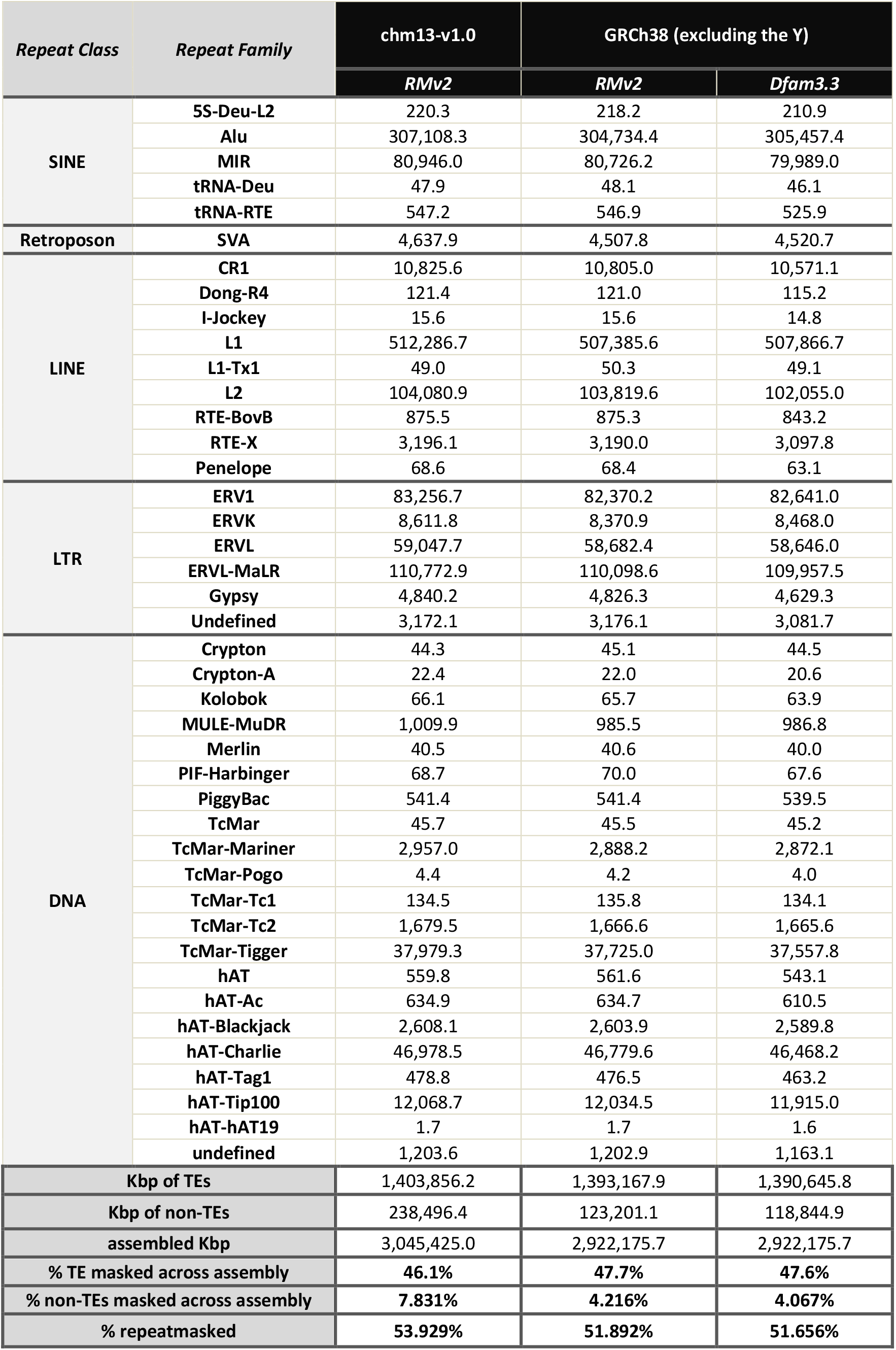
Repeat annotations are more refined for CHM13v1.0. Kbp of repeat annotations, by repeat class and family, for different human genome assemblies based on CHM13v1.0 RMv2, GRCh38p13 RMv2, and GRCh38p13 Dfam3.3 only. Note: the newly added AluJb is included in the Alu repeat family category.

The re-annotated GRCh38 and newly annotated CHM13v1.0 were compared using reverse liftOver coordinates (CHM13 to GRC38) to identify TE insertions specific to CHM13 (Note S2). TEs found in CHM13 and not found in GRCh38 were further grouped into those that are in gap-filled regions (non-syntenic overlap) or that are potentially polymorphic between these two genomes or were collapsed in the GRCh38 assembly (syntenic overlap but missing in GRCh38) (Fig. 1C). Across 4,577,617 TEs with lifted coordinates (i.e., shared between CHM13 and GRCh38), 148,882 lifted TE pair annotations were discordant between the two genomes (Fig. S5A); 97.2% of these (144,670 discordant liftOver pairs) were subfamily reclassifications. These discordant pairs are typically short loci with low scores and are the result of different matrices automatically chosen by *RepeatMasker* between assemblies leading to slightly differing TE annotations and not reflecting true assembly sequence differences (Fig. S5B, Table S5). Among the 17,434 unlifted TEs unique to CHM13, all TE classes are represented (Fig. 1C) with 27.2% of TE sites specific to gap filled regions in CHM13 (4,750 total TEs) (Table S5, S6). Unlifted TE sites are found genome-wide, with a higher density identified on the acrocentric chromosomes 14, 15, 21, 22 (Fig. S5C, Table S7).

While many of the new repeat classifications coincide with gaps filled in the new assembly, the additional data provided by this assembly supported the annotation of previously undiscovered repeats and TEs genome-wide (Fig. 1B). To determine if these novel repeat classifications were unique to human, we searched for orthologous sequences in the previous human reference (GRCh38) and available genome assemblies for primates representing the great apes (*Pan troglodytes, Gorilla gorilla, Pongo abelii*), Hominoidea (*Hylobates moloch*), Catarrhini (*Macaca mulatta, Rhinopithecus roxellana)* and Platyrrhini (*Callithrix jacchus, Microcebus murinus*) (Note S2). Relative copy numbers of each repeat classification vary between CHM13 and GRCh38, a reflection of the incompleteness and collapsed regions within the latter assembly and copy number differences between human haplotypes. When comparing copy numbers of new repeat annotations between CHM13 and long-read, high quality assemblies available for other great apes (chimpanzee, gorilla and orangutan) (*23*), we still find a dramatic increase in copy number across most of the novel repeats in the human genome (Table S8). Many repeats appear only as monomers in other primate genomes or are absent in Platyrrhini, Catarrhini, and lesser apes; these reduced counts are largely influenced by the quality of these assemblies and potentially high rates of divergence among repeats, and highlight the need for telomere-to-telomere assembly approaches for comparative analyses (*24*). Finally, eight of the novel repeats are human-specific, with an additional eleven found only as monomers in other species (Fig. 1D, Table S8).

### Composite elements shape human-specific genomic variation

The complete CHM13 assembly provided a unique opportunity to identify complex repetitive “composites” as well as their evolutionary history in the primate order. We annotated 19 composite repeat elements (Table S2), defined as a repeating unit consisting of three or more repeated sequences, including TEs, simple repeats, novel repeat annotations and satellites (Note S2). This effort included updating previously known composites with new subunit repeat annotations and new TE classifications, chromosomal locations and overall structure. Composite repeats in this context are distinguished from composite retroelements (such as SVA (*26*) and LAVA (*27*)) that are capable of retrotransposition; rather, complex composite core units are arranged in tandem arrays as “megasatellites” or “macrosatellites”(*28*), likely derived through unequal crossing over that contributed to their copy numbers. Composites represent a class of structural elements that contribute to human diversity and disease through structural variation (SV) and copy number variation (CNV), particularly when exonic regions are “captured” in a core unit (e.g. (*29*)).

Composites are subdivided into categories that are defined by the different evolutionary forces that shaped their distribution in the human genome. Most composites are found in a tandem array only on a single chromosome (Fig. S6A-F, S7B-G), and in eight cases each core unit contains protein-coding annotations (Fig. S7), indicating that unequal crossing over events and concerted evolution among composite units can contribute to the expansion or contraction of gene families within humans (Fig. 1D). Several of these composites were previously annotated as staggered segmental duplications (*19*) encompassing only the tandem array (e.g. Fig. S8), illustrating the challenges in annotation of composite arrays and individual composite subunits. Interestingly, the LMtRNA (Fig. S8A) composite array is bounded by two full length HERV elements (HERV-L and HERV-9C) in inverted orientations, implicating them in the formation of the array in an ancestral primate genome.

One composite, 5SRNA_Comp, consists of a portion of the 5SRNA, an *Alu*Y and two subunit repeats and is found as an array of 128 repeating units with high sequence similarity (100%) on Chromosome 1 (Fig. S9A,B). Monomers of 5SRNA_Comp are found at 49 locations across 13 chromosomes, enriched for centromere transition regions therein (Fig. S9C). All monomeric forms lack the *Alu*Y; rather they carry an LTR2 at the same location (Fig. S9D). Thus, the distribution of monomers is likely the result of segmental duplication events through non-allelic homologous recombination (NAHR), while the *Alu*Y insertion (resulting in deletion of the LTR2) in one copy preceded the expansion of this composite into a high copy number array. Using methylation profiles developed for CHM13 and long-read based methylation clusters (*22*), we find that the methylation pattern of the 5SRNA_Comp is not consistent across the array, rather we find a methylation dip region (MDR) internal to the array, strikingly similar to the centromere dip region (CDR) identified in higher order arrays of alpha satellites in CHM13 (*22*) (Fig. 2A). The location of the MDR is not linked to DNA sequence as neither the GC content nor sequence identity is variable across repeat units in this array (Fig. 2A, S9B), implicating other epigenetic factors in defining the drop in methylation.

**Figure 2.**
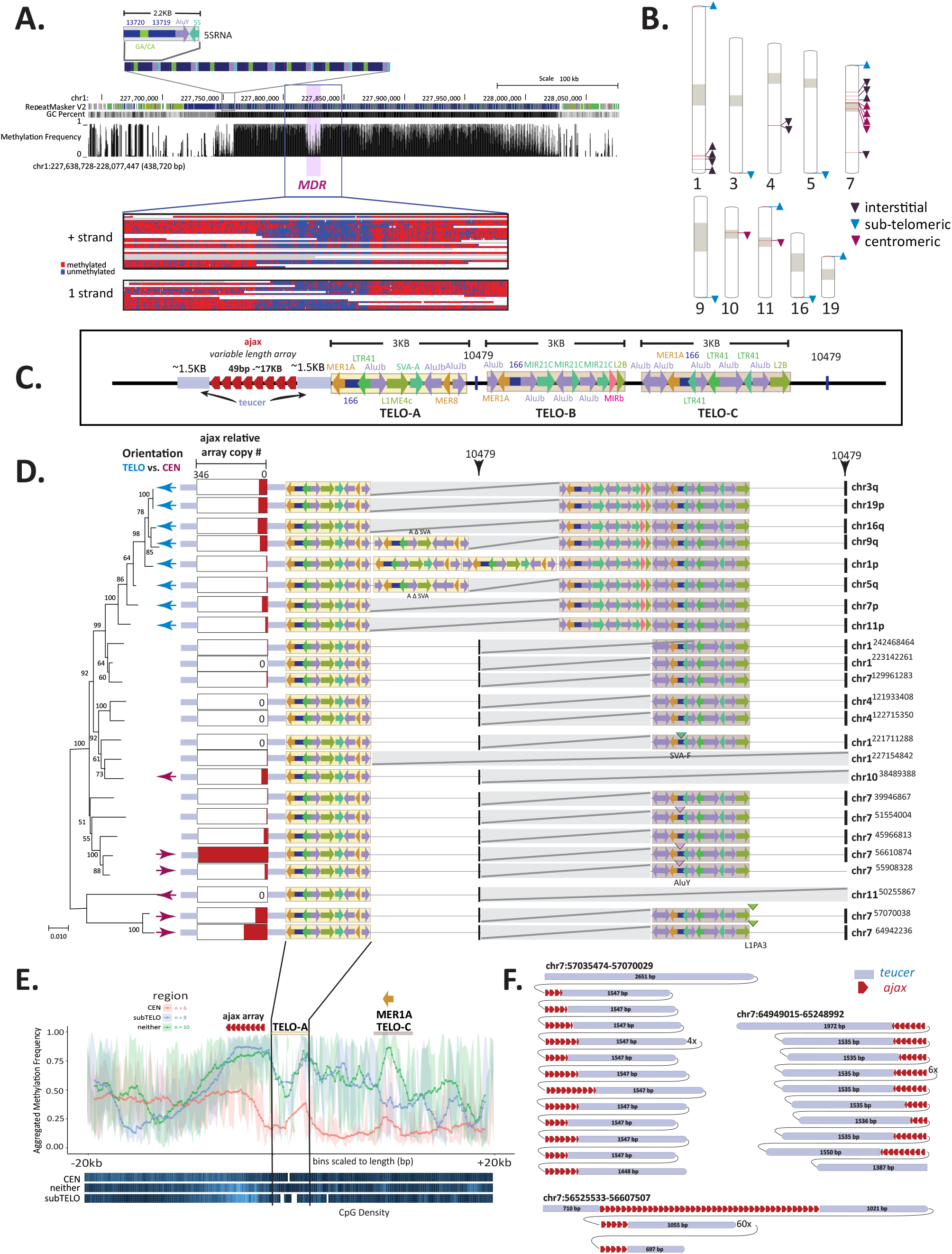
Novel composite elements identified in CHM13 reveal shared repeat structures and epigenetic patterns across chromosomes. **(A) (Top)** CHM13v1.0 Genome Browser showing the 5SRNA_Comp subunit structure and array (top, middle). RepeatMaskerV2 track, CG percentage, and methylation frequency tracks are shown. The methylation dip region (MDR) is indicated. **(Bottom)** Zoom of individual nanopore reads showing consistent hypomethylation in the MDR (chr1:227,818,289-227,830,789) and hypermethylation in the flanking regions (chr1:227,804,021-227,845,689). Both positive (top) and negative (bottom) strand aligning reads show the same methylation pattern. **(B)** The location of TELO-composite elements across CHM13 is indicated by red bars on chromosomes. Tan blocks demarcate centromeres and centromere transition regions. Chromosome regions containing TELO-composites across the karyotype (22) are color coded [interstitial – purple, sub-telomeric, within 200Kbp of chromosome end– aqua, centromeric – red] with orientation indicated by arrow direction. **(C)** Each CHM13 TELO-composite element consists of a duplication of a teucer repeat (blue) separated by a variable 49bp (ajax) repeat array (red arrowheads) and three different composite subunits (TELO-A, -B, -C). Repeat and TE annotations are shown. Some copies of TELO-composite contain the novel repeat “10479” between the TELO-A and TELO-C subunits, and/or following the TELO-C subunit. **(D)** Schematic alignment of all complete CHM13 TELO-composite elements (location indicated to the right, centromeric and interstitial locations indicated with starting bp). Locations of “10479” (black box) repeat are arrowed (top). Relative number of each TELO subunit as pictured in (**C**) with deletions represented by grey bars. Additional TE insertions are indicated for TELO-C. Relative array size of ajax repeats shown to scale among all TELO-composite elements. Orientation of the element indicated with respect to the centromere (purple arrow) telomere (blue arrow) and interstitial (no arrow) regions. Left, RaxML phylogeny of TELO-composite elements with bootstrap values at each node and distance indicated by length of branch. **(E)** Metaplot of aggregated methylation frequency (average methylation of each bin across the region, 100 bins total) centered on the TELO-A subunit, ±20Kbp, grouped by chromosomal location (orange – centromeric, blue – subtelomeric, green – interstitial). CpG density for each group is indicated at the bottom (white -no CpG, dark blue -low CpG, bright blue -high CpG). The location of the ajax repeat array and the MER1A element within the TELO-C subunit are indicated (top). **(F)** Organization of the Chromosome 7 locations of the ajax repeat (each red arrowhead denotes a single monomer of the repeat) and teucer (blue). “#X” indicates a subunit is arrayed at # copy number.

Two composites are found arrayed across several chromosomes. The ACRO_Comp (*30, 31*) is a unit found across 12 chromosomes (Fig. S10A), including as tandemly arrayed sequences across the five acrocentrics (Chromosomes 13, 14, 15, 21, 22) with high sequence identity across composite units (Fig. S10B). The LSAU-BETA composite is found across 16 chromosomes and in both tandem arrays and as single monomers (Fig. S11A, B). The LSAU-BETA composite has a variant form (LSAU-5403) in CHM13 (Fig. S11B) and includes subunit repeats consisting of D4Z4 (*32*) and LSAU ((*33*), overlaps with the DUX4 genes and microRNA genes (MIR8078) and has been implicated in facioscapulohumeral muscular dystrophy (FSHD)(*32, 34*). Complete reference sequence spanning these complex arrays afforded the opportunity to assess intra-array variability. We find that LSAU composites found in centromere transition regions share lower identity within an array (80-95%) than LSAU composites found within interstitial arrays (near 100% identity), illustrating the utility of the CHM13v1.0 reference for future studies of the evolutionary trajectories of repeat arrays contextualized to chromosome location.

In addition, we annotated a highly complex composite, TELO_Comp, that consists of multiple arrays and other composites (Fig. 2B,C). TELO-Composites are found on 10 chromosomes (Fig. 2B) at interstitial, pericentromeric and subtelomeric loci. The canonical TELO_Comp consists of three 3Kbp composites (TELO-A, -B, -C subunits), each containing multiple TEs, downstream of a variable length array of a 49bp satellite repeat unit, ajax, bounded by a duplicated sequence, teucer (Fig. 2C). Across 24 loci, all TELO_Comp elements contain a TELO-A subunit downstream of the ajax satellite array (Fig. 2D). Among the subtelomeric elements (Fig 2D, blue arrows), all contain TELO-B and TELO-C subunits upstream of a shared subunit repeat found across all TELO-Comp elements (10479). In depth analysis of the overall structure of the subunits across all loci, and phylogenetic analyses of the TELO-A subunit (Note S3, Table S9), indicates that subtelomeric units are a monophyletic group of recent origin, while interstitial and pericentromeric units are polyphyletic. Elements within this latter group lack a TELO-B subunit; between the TELO-A and TELO-C subunits, all of these TELO_Comp elements contain a second 10479 subunit repeat, with the exception of three elements (Chromosomes 1, 10, 11) that also lack a TELO-C subunit. While high bootstrap support for the clustering of subtelomeric elements indicates recent derivation, likely by segmental duplication events (Fig. S12A, Table S10), location-specific repeat diversification in subunit content and structure as well as ajax repeat copy numbers, which retain high sequence identity (Fig. S12B) is observed. Moreover, each subtelomeric unit contains the ajax array proximal to the telomere, indicating that inverted orientations are favored at subtelomeric loci.

Meta-analysis of aggregated methylation frequency across the TELO-Comp units (+/-20Kbp) (Fig. 2E) shows that the ajax satellite array is hypermethylated across all elements, with a discernable drop in methylation across TELO-A subunits and peak of methylation in the MER1A unit in elements containing TELO-C. Subtelomeric and interstitial TELO-Comp elements share similar methylation profiles, with higher methylation levels across the entire element, while pericentromeric TELO-Comp units have lower overall methylation levels. This implicates local epigenetic states impact overall methylation levels but do not change relative levels within the ajax array and TELO-subunits. Comparison of aggregated methylation frequency across TELO-Comp units at the same loci in the diploid assembly for HG002 (Fig. S13)(*22*), show that overall methylation levels are higher across all TELO_Comp elements, including those found in centromeres, as expected from global differences in methylation level between CHM13 and HG002. However, the overall methylation pattern for the TELO-Comp elements (Fig. 2E, S13) is retained, indicating it is an epigenetic signature of this repeat.

On Chromosome 7, three TELO_Comp loci were found that contain additional tandem arrays of the TELO-Comp subunit consisting of ajax and teucer sequences, with variable ajax repeat numbers and variable tandem arrays of the teucer-ajax subunit (Fig. 2F). Further phylogenetic analyses for both ajax and teucer sequences reveal subtelomeric arrays evolve under neutral evolution while pericentromeric arrays evolve under concerted evolution (Fig. S14-S15, Table S11-S12). Moreover, phylogenetic analyses of ajax-teucer monomers from the Chromosome 7:56525533 locus indicate that both elements form a composite that evolves as a single unit, suggesting a higher-order repeat, or super-repeat, structure across that locus (Fig. S14-S15). Collectively, the inclusion of composite elements in the annotation tracks for CHM13v1.0, afforded by the polished and contiguous reference assembly, will provide the research community with a set of guideposts around which to pinpoint potentially pathogenic variants.

### TE-driven Genomic DNA Transductions and Their Evolutionary Consequences

TE-mediated transduction is a process by which retroelements co-mobilize DNA flanking either their 3’ or 5’ end to new genomic loci (*35*–*38*). Several studies have shown that transductions mediated by non-LTR retrotransposons, i.e., *Alu*, L1, and SVA are common in the human genome (*35*–*37, 39*–*41*). 3’ transduction is a consequence of a weak 3’ polyadenylation signal present in the source element, which is skipped by a polymerase during transcription. In this scenario, transcription continues until another, downstream polyadenylation signal is reached (*36, 37*). In contrast, 5’ transduction most likely occurs when a fraction of a retrotransposon is transcribed from an external promoter (*38*). Transductions can affect the genome in a number of ways. For instance, an exon can be transduced along with a retrotransposon and integrated into another gene resulting in exon shuffling (*35, 39, 42*). Moreover, it has been demonstrated that the transduction process is associated with the development of some diseases and is a possible source of somatic mutations during tumorigenesis (*43, 44*). Here, we applied a set of computational approaches to discover all transduction events caused by active human retrotransposons, i.e., L1s, *Alu*s and SVAs, in CHM13.

To catalog and characterize high-confidence transduction events mediated by non-LTR retrotransposons in CHM13, we utilized a modified TSDfinder tool (*35*) and transduction annotations were assigned to confidence bins 0-4 (Note S4, Fig. S16). In total, we analyzed 1,182,410 *Alu* elements, 971,811 L1s and 7,069 SVAs annotated in CHM13 (Fig. S3A) with lengths that ranged from 10bp to 18,712bp (median of 293bp). Of these, 183, 76, 55 and 386 events were annotated for L1 3’, SVA 5’, SVA 3’, and *Alu* 3’, respectively, as transductions with the highest confidence (Table 3). Although the number of transductions induced by *Alu* elements is the highest among all elements, when normalized to the overall count of elements within the genome, SVA elements appear to be the most productive TEs. Generating mostly interchromosomal transduction events (Fig. 3A), SVAs generate 1.85% transductions per element, compared to 0.033% and 0.019% for *Alu* and L1, respectively. When comparing transduction events between CHM13 and GRCh38 (Table 3A), we find more events in CHM13, due to the gap-filled regions and high confidence annotations. For example, 6% of transductions annotated as level 4 were found within newly assembled centromere regions. Moreover, we find that the number of 5’ transduced segments mediated by SVAs is comparable with SVA 3’ transductions, suggesting that this type of transduction is a common phenomenon in the human genome. Interestingly, the median length in *Alu* 3’ transductions was considerably greater than other elements (Table 3B).

**Table 3.**
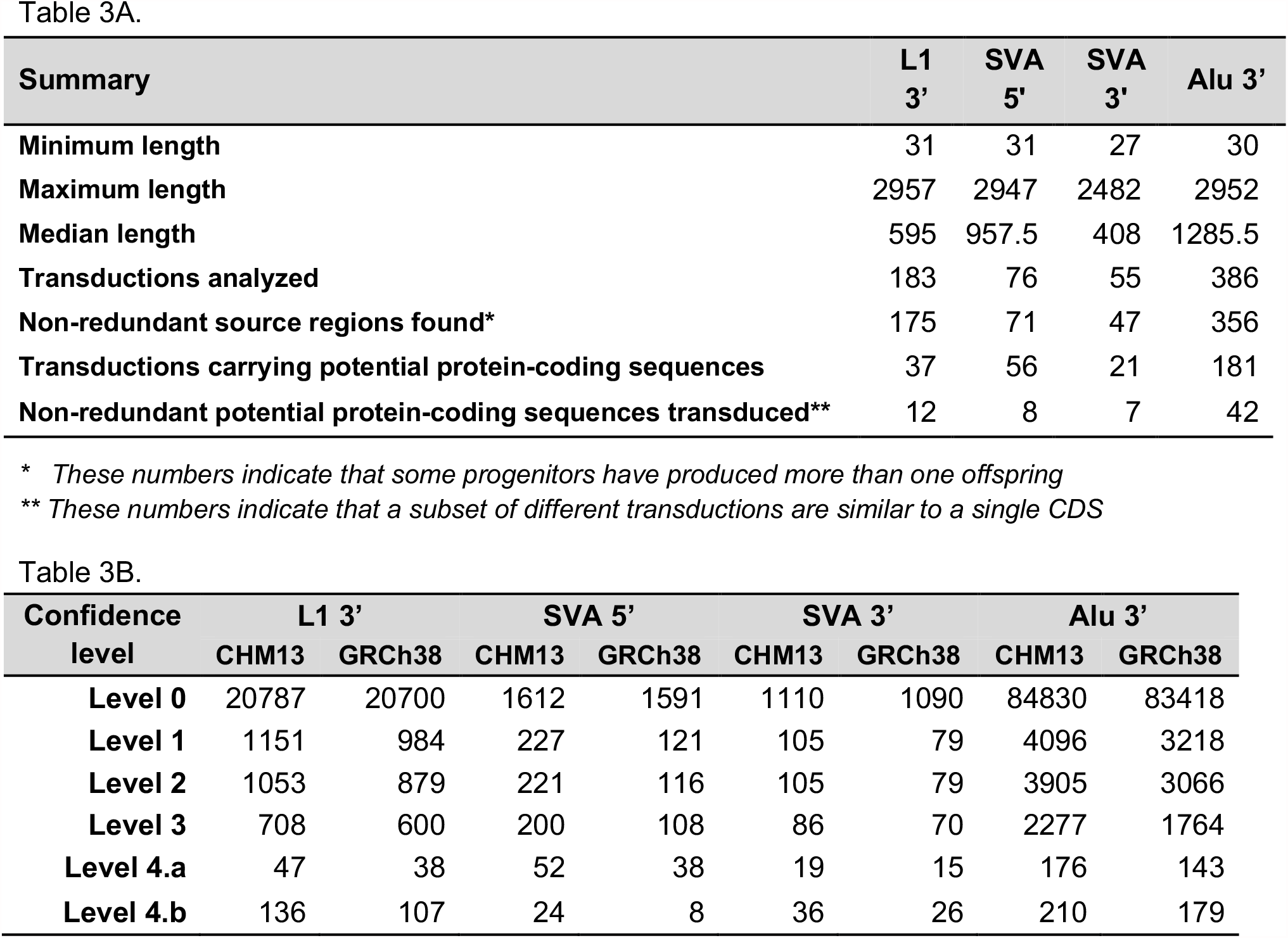
Transduction events by the most active family of TEs shape the human genome. **(A)** Statistics of transduced DNA fragments (highest confidence cases -level 4.a./4.b). **(B)** Statistics of transduced DNA fragments (highest confidence cases -level 4.a./4.b).

**Figure 3.**
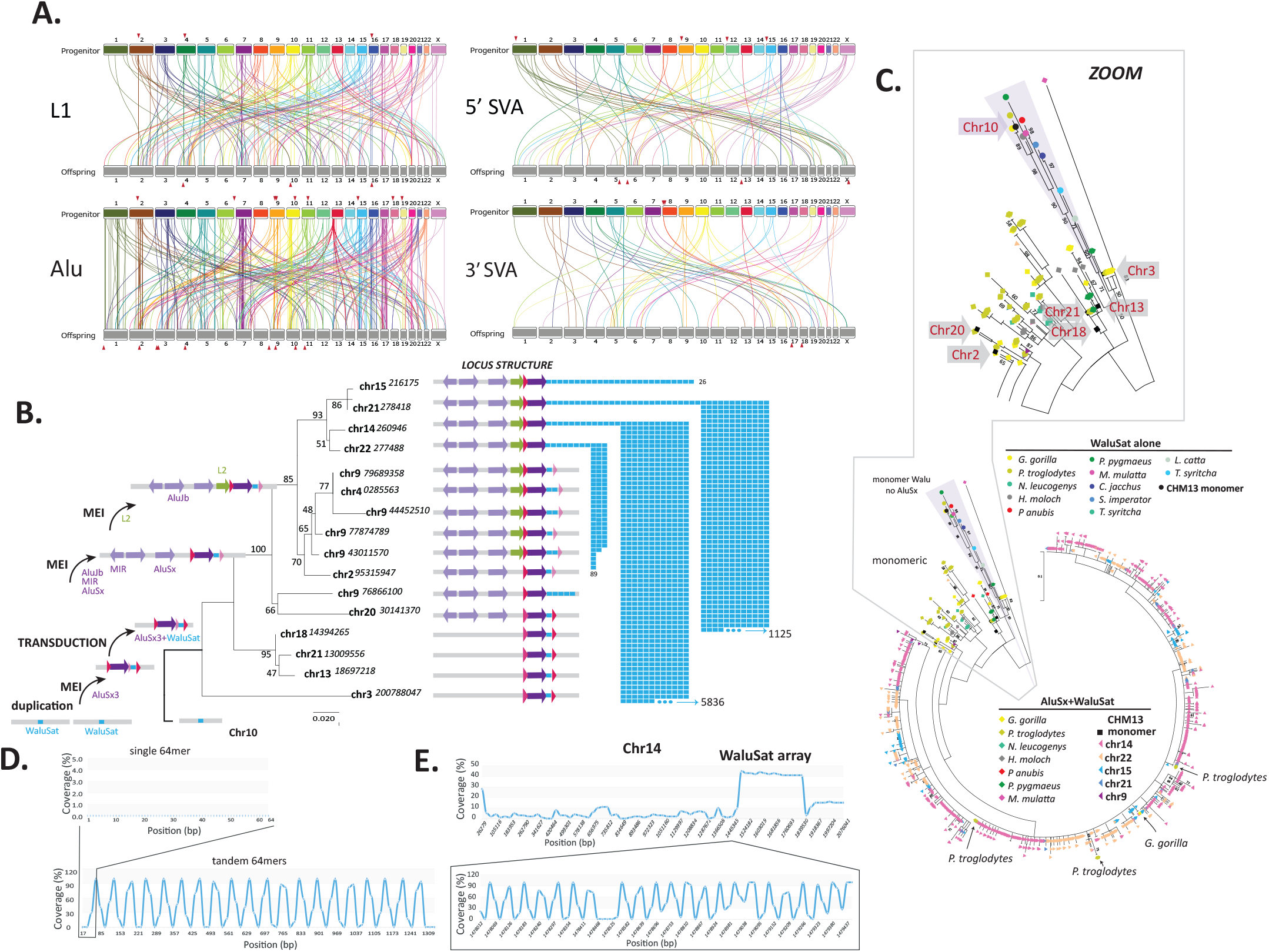
Transduction events in the human genome diversify repeat and gene content. **(A)** Linkage graphs showing transduction progenitors and offspring across CHM13 for each repeat class (LINE, SINE, 5’ and 3’ SVA transduction events). Connections are color coded by the progenitor chromosome. Connections corresponding to transduced sequences with >90% identity to protein-coding sequences are indicated with red arrowheads (Table S13, Fig S19). **(B)** Maximum likelihood (ML) phylogenetic analyses of the AluSx3-WaluSat locus across CHM13. Chromosome location is indicated (starting nt position shown) at each branch. Bootstrap values shown at each node, distance indicated by length of branch. Left shows the sequential order of events, initiating with a duplication of the Chromosome 10 WaluSat locus followed by mobile element insertion (MEI) of an AluSx3 into one copy. This MEI event led to the transduction of WaluSat and insertion of AluSx3-WaluSat across four chromosomes (3, 13, 21, 18). MEI events upstream of the AluSx3-WaluSat are concordant with phylogenetic relationships among loci and indicate that the derivation of AluSx3-WaluSat loci across other chromosomes were the result of segmental duplication events. Once the AluSx3-WaluSat was duplicated to the acrocentric chromosomes 14, 15, 21, 22, a massive expansion of the WaluSat sequence (blue boxes) occurred. The number of WaluSat monomers within each array is indicated on the right. **(C)** Phylogenetic analysis of the WaluSat monomer across primate lineages. ML analyses show the WaluSat sequences transduced by AluSx cluster together (main circle), as do the WaluSat monomers (Chromosomes 2, 3, 10, 13, 18, 21) monomers (boxed), which are found in Catarrhini and Hominoidea primates. Only the WaluSat monomer on CHM13 Chromosome 10 clusters with Hominoidea, Catarrhini, Platyrrhini, and Prosimians (zoom), indicating this locus is the progenitor of the satellite repeat. **(D)** G-quadruplex (G4) analysis of a single 64-mer monomer of the WaluSat sequence showed no predicted G4 structures (top), while an in silico construct of a tandem array of the WaluSat shows high G4 coverage at the junction between individual WaluSat monomers across the array. **(E)** G4 analysis of the p-arm of Chromosome 14 shows a peak in G4 predictions coincident with the WaluSat array. Bottom is a zoom inset of a subset of the array showing that the junctions between most monomers carry predicted G4 structures.

Our results indicate a high number of transductions in the human genome with an average of 1.24 events per 1Mbp. Chromosomes 7, 2, 17, 4, and 11 are the most enriched targets with 67, 56, 43, 40, and 39 transduced loci, respectively, whereas Chromosomes 19, 21, 12, 8, and 18 have the lowest number of events, with 19, 18, 17, 15, and 7, respectively (Fig. 3A and Fig. S17, S18). We find that truncated elements are capable of transducing sequences and can produce as many offspring as full-length elements. Among all high confidence, verified transductions (700 events in total), 231 transduced loci did not match the same TE-subfamily assignment as their identified parents. While it is expected that both progenitors and offspring would belong to the same TE-subfamily, annotation discrepancies may be due to the fact that the resulting copies are subjected to independent mutation and locus-conversion processes (*45*).

To assess the potential impact of TE transduction events on protein-coding gene evolution via exon shuffling, we compared each transduced sequence with the human proteome. A considerable fraction of transduced DNA events (Table 3B) exhibit similarity to a human protein coding sequence with an identity between 24.5% and 100%, suggesting either exon shuffling or gene duplication mediated by transductions(*42*) (Fig. 3A, Fig. S19). One notable example is the formation of a paralogous protein called SLC35G4 (NP_001269229.1), solute carrier family 35 member G4, mediated by an SVA_C 3’ transduction in which an intronless protein-coding gene called SLC35G5 (chr8:8406845-8408078) has been transduced and generated two offspring (chr17:36141023-36142703 and chr18:11771758-11772991)(*39*). In sum, our results indicate TE facilitated transduction is a dynamic and common phenomenon that has affected 0.026% of the CHM13 genome. It is worth noting that our transduction annotation is likely an underestimation of the total number of events given the high stringency thresholds employed. However, the CHM13 assembly has afforded a multi-tier analysis that can be further applied to identify bona fide transduction events in lower confidence categories (Table S13).

Among the gaps assembled in CHM13 (*12*), we discovered previously unannotated repeat arrays of a 64nt sequence (Fig. 1B) present in high copy numbers on the short (p) arms of acrocentric Chromosomes 14,15, 21, and 22 (*12*). The 64nt repeat unit is found in single or low copy number (<5) on eight other chromosomes (Fig. 3B). Comparative and phylogenetic analyses of the structural features across all loci revealed that a solo monomer resides on Chromosome 10, while all other occurrences are adjacent to an *Alu*Sx3 element (hence <u>with *Alu* sat</u>ellite, or WaluSat). Moreover, the retention of target site duplications on a subset of these *Alu*SX-WaluSat loci indicates WaluSat was transduced by an insertion of an *Alu*Sx element and mobilized to several chromosomes (Chromosomes 3, 13, 21, 18). Subsequent TE insertion events (Chromosomes 9, 20) upstream of the *Alu*Sx-WaluSat mobile element insertions allowed for the delineation of segmental duplication events that spread the *Alu*Sx-WaluSat to locations on Chromosomes 4, 9, 14, 15, 21, and 22. Once segmental duplication events placed the *Alu*Sx-WaluSat on the p arms of chromosomes 14, 15, 21 and 22, WaluSat amplified from 26 copies (Chromosome 15) to 5,836 copies (Chromosome 14).

Phylogenetic analyses of monomers and tandem arrays of the WaluSat repeat revealed that the amplified WaluSat arrays on the acrocentric p arms are highly similar to each other, and less similar to solo monomers, which instead bear more similarity to the solo WaluSat monomers in other Catarrhini species (Fig. 3C). The presumably ancestral 64nt WaluSat on human Chromosome 10 forms a monophyletic branch with WaluSat in primates inclusive of Prosimians, supporting the hypothesis that WaluSat on Chromosome 10 was duplicated prior to transduction by an *Alu* in the last shared common ancestor with Catarrhini (Fig. S20) and indicating that WaluSat monomers evolve independently of *Alu*Sx-WaluSat (Fig. S21). We hypothesize that the high degree of similarity and copy number variation among p arm WaluSat arrays is due to frequent non-allelic or ectopic recombination events on acrocentric chromosomes (*12, 19*), which may be exacerbated by replication challenges associated with the predicted periodic G-quadruplex structures (*46*) identified at junctions of WaluSat sequences within arrays (Note S4, Fig. 3D,E).

### Transcriptional, epigenetic, and structural differences define TEs across the human genome

PRO-seq detects nascent transcription from all RNA polymerases with nucleotide resolution at genome-scale. The resulting read density profiles quantitatively reflect the occupancy of active polymerases across the genome. Sites of accumulating RNA polymerase activity (*20, 47*), such as promoter-proximal pause sites, 3’ cleavage/polyA regions, splice junctions and enhancers, indicate points of transcription regulation (*20, 48*). In addition, because PRO-seq captures RNA synthesis before mechanisms that affect RNA stability take place, unprocessed and unstable RNAs can be readily detected with high sensitivity. The latter is a critical advantage over RNA-seq which primarily detects accumulated processed and stable RNAs. As part of the CHM13 gene (*12*) and repeat annotations, we assessed the sites of nascent transcription and the density of active RNA polymerases at single nucleotide resolution genome-wide with PRO-seq (*20*)(Note S5). Coupled with epigenetic information (*13, 22*), these data can be used to assess how repetitive elements influence genome structure and function.

We therefore used PRO-seq to provide the first profiles of RNA polymerase activity that distinguish different families and activity classes of retroelements in the human genome. As an example, we focused on the TE derived macrosatellite SST1 (also called MER22 (*49*) and NBL2 (*50, 51*)) that has demonstrated meiotic instability (*52*) and whose methylation status is of clinical relevance to multiple cancer types, including neuroblastoma, ovarian (*53*), breast (*53*), gastric (*54, 55*), colon (*53*), and hepatocellular (*56*) carcinomas.

Arrays of SST1 monomers had been identified on the arms of Chromosomes 4 and 19 (*52, 57*) and were clustered in centromeric regions (*49*). In CHM13, we report 613 SST1 loci (708,553 bases) (Fig. 4A, B, Table S14), and, while SST1 arrays are reported to be variable in the population (*52*), our annotations represent a ∼2-fold increase over the 342 loci (315,515 bases) identified in GRCh38 (excluding the Y chromosome, which carries an additional 587 loci, Fig. S4). To determine the evolutionary relationships of SST1 monomers and arrays across human chromosomes, we first performed a RAxML phylogenetic analysis with representative loci subsampled from the 16 autosomes on which SST1 resides (Note S3, Fig. 4A, Table S15). The array situated on the long (q) arm of Chromosome 19 represents the ancestral site of SST1 in the human genome succeeded by its diversification into the centromeric arrays and the interstitial array on Chromosome 4. In lesser apes (gibbons), SST1 arrays are found in centromeres formed from the orthologous region of the array on human Chromosome 19 (*58*), and SST1 is found as a centromeric satellite in Old World monkeys (*49*), indicating that the Chromosome 19 locus carries a high propensity for centromere seeding and array size expansions/contractions across primate lineages.

**Figure 4.**
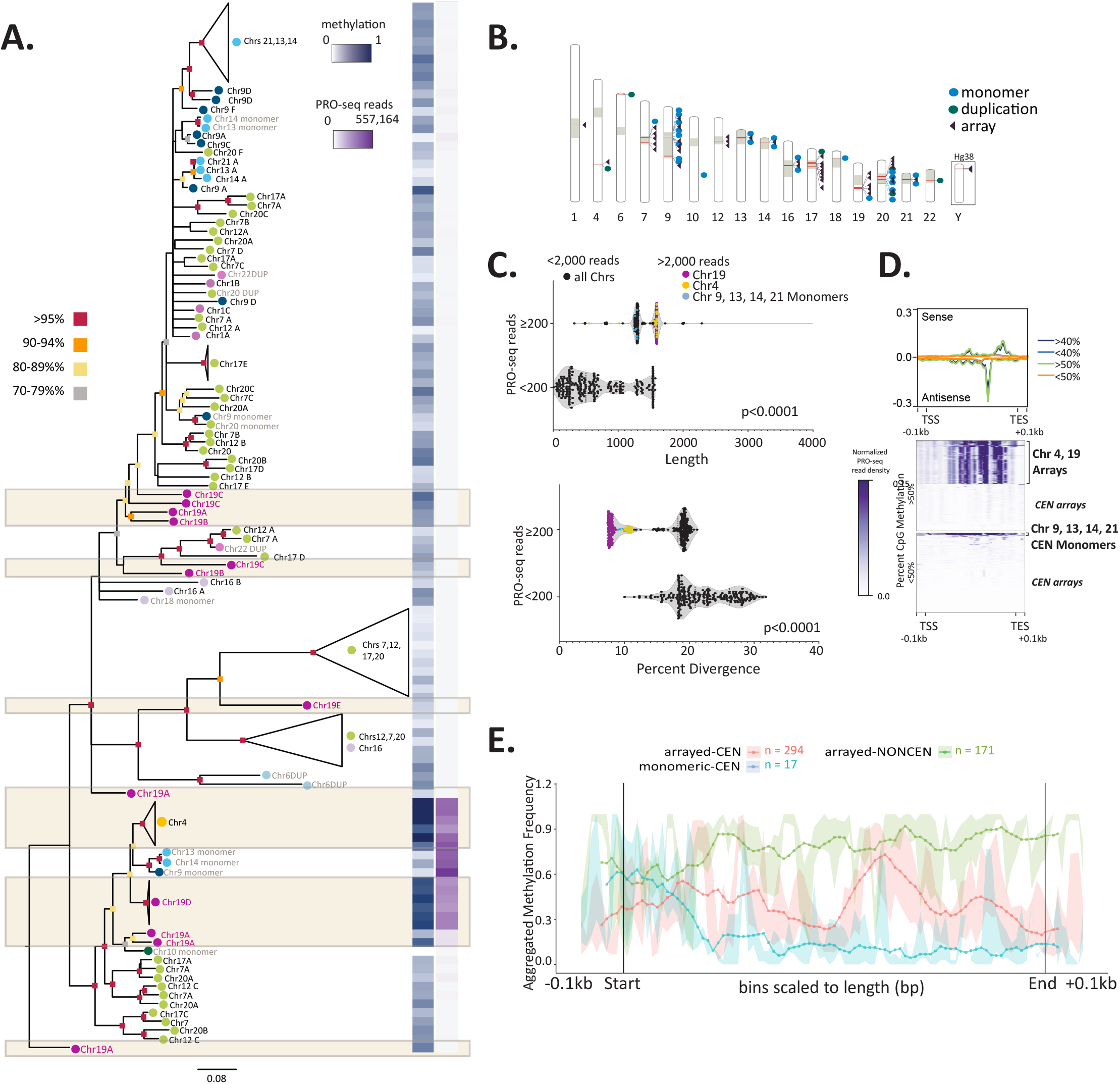
Transcriptional, epigenetic and structural differences define SST1 elements across the human genome. **(A)** RAxML phylogenetic analysis of SST1 elements (subsampled to represent each chromosomal location, and aligned using MAFFT (66), Tables S14-S17). Bootstrap values are indicated by color (as per key to the left) at the base of each node. Branch lengths indicate distances and unresolved nodes were collapsed. “Chr#” followed by A-F indicates the array designation by CHM13 chromosome unless SST1 is present as a monomer or as duplicons (DUP) (indicated in grey text). Colored circles by chromosome labels indicate phylogenetic clusters (e.g. green – Chromosomes 7, 12, 17, 20; aqua – Chromosomes 13, 14, 21). Right: For each SST1 sequence or group of collapsed sequences on the tree, average methylation frequency (0 -hypomethylated; 1 -hypermethylated) is indicated in blue and PRO-seq read coverage is indicated in purple as per key inset. **(B)** The location of SST1 elements across CHM13 is indicated by red bars within the chromosome schematic (Table S14). Tan blocks indicate centromeres and centromere transition regions as per (13). SST1 arrangement as a single monomer (blue dot), duplication (green dot) or array (purple triangle) is indicated. Locations of SST1 arrays on the Y chromosome are shown for GRCh38 (CHM13 is 46,XX). **(C)** Violin plot of SST1 elements shows statistically significant differences between expression levels (repeat overlap of PRO-seq reads) and length of the element (t test, p<0.0001) as well as percent divergence (t test, p<0.0001). Dot colors indicate interstitial arrays on Chromosome 19 (purple) and Chromosome 4 (yellow) and centromeric monomers of SST1 found on Chromosomes 9, 13, 14, 21 (blue) with a read overlap higher than 2,000. All locations with a read overlap lower than 2,000 are indicated in black. 200 and 2,000 read overlap cutoffs determined by analyzing the range of read overlap among all SST1s (Fig. S22). **(D)** CHM13 PROseq profiles (upper panel) of SST1 grouped by average methylation levels (< and > 50%). Each element is scaled to a fixed size, TSS (transcription start site), TES (transcript end site), and ±0.1Kbp are shown (bottom). Heatmaps (lower panels) of PRO-seq density (purple scale, normalized reads per million aggregate for sense and antisense) grouped by average methylation levels (>50% top, <50% bottom). Clusters of specific SST1 loci are indicated to the right. **(E)** Metaplot of aggregated methylation frequency (100 bins total) of SST1 elements (500bp-2Kbp), ±0.1Kbp, grouped by chromosomal location and arrayed vs monomeric/duplicated (orange – centromeric (CEN) array, blue – centromeric monomer, green – noncentromeric array). Truncated noncentromeric/CEN monomers and duplications not shown; length filtering resulted in n=1.

The number of overlapping PRO-seq reads, average methylation, and percent divergence for each SST1 element in CHM13 was compared to delineate correlations between transcriptional, epigenetic, and structural features of SST1 across genomic loci. PRO-seq revealed that the SST1 arrays on Chromosomes 4 and 19 and centromeric monomers on Chromosomes 9, 13, 14, and 21 are highly transcribed in comparison to other SST1 loci and are found within a single phylogenetic cluster (Fig. 4A, C, D, Note S5-S6, Figure S22, Table S16), indicating that centromeric SST1 repeat arrays are transcriptionally inactive. Statistical analyses of SST1 repeats showed that the highly transcribed repeats are both longer and less diverged from the consensus sequence (*t* test, p<0.0001) (Fig. 4C, Fig. S23, Table S17) despite their basal location in the phylogenetic tree (Fig 4A). CpG methylation levels are high (>50%) for SST1 within Chromosome 4 and 19 arrays, low (<50%) for centromeric monomers and variable (low and high) for centromeric arrays (Fig. 4A, D, Fig. S23, S24, Table S16). Metaplots of aggregated methylation frequency across SST1 repeat units support this observation and indicate that while interstitial arrays and monomeric SST1s carry the same methylation frequency at their 5’ end, monomeric SST1s lose most methylation across the body of the element (Fig. 4E, Fig. S24, S25). Irrespective of this methylation pattern, heat maps of PRO-seq density show all highly transcribed SST1s have two internal peaks of high RNA polymerase occupancy that are closely spaced and in opposite orientations (Fig. 4D, Fig. S24B), characteristic of RNA pol II promoters and enhancers.

Combined, these data suggest selective pressure to retain the genomic integrity of older, less diverged SST1 arrays and monomers that are actively transcribed, while silenced repeats found in centromeric arrays are more susceptible to sequence variation. Contrary to expectations that CpG methylation renders repeats transcriptionally silent (*59, 60*), we find that high levels of average methylation across interstitial, arrayed SST1s define these highly transcribed repeats (Fig. 4A, D, E) and bears a resemblance to methylation patterns observed over gene bodies (*61, 62*). While numerous studies have reported chromosomal instability and cancerous phenotypes associated with demethylation of SST1 repeats (e.g. (*63, 64*)), refining the annotations for SST1 genome-wide will support work to assess which SST1 repeats and genomic locations are variable and differentially transcribed/methylated in specific disease states as well as delineate defining epigenetic marks and proteins (*53, 65*).

Capitalizing on the single-base resolution of nascent transcription and methylation profiles, we set out to define first profiles of RNA polymerase activity that distinguish different families and activity classes of retroelements in the human genome. First, we collected all TE subfamilies with elements that remain active in humans (*Alu*Y, HERV-K, SVA-E/F, and L1Hs) and divided them into elements that still carried the potential for mobilization (full length) (Fig. S26, Table S18), and those that are truncated and thus lack mobility (immobile) (Note S7). We then assessed PRO-seq signal and CpG methylation density across each element within each subfamily and category. For each element type, density profiles were correlated with known features of specific repeats, such as ORFs, promoters, and long terminal repeats (LTRs). For each element within a category, we further assessed correlations among methylation frequency, CpG site density, and sequence divergence from the consensus.

Across all full length retroelements in CHM13, PRO-seq density profiles show clear signals of RNA polymerase accumulation (Fig. 5A-E, Fig. S27). *Alu*Y elements show two signal peaks; the first corresponds to the known RNA pol III promoter site within the first monomer, while the second, broader peak indicates the site of a second, ancient 7SLRNA promoter (*67*), whose presence might promote polymerase pausing (Fig. 5A). The peak distribution closely mimics the relative size of the left and right *Alu* monomers. While active transcription continues in immobile *Alu*Y elements, there is no longer visible signal of promoter exclusivity and RNA polymerase signal spreads across the element. Full length *Alu*Y elements retain a strikingly similar methylation profile and show low divergence levels; truncated and older elements (*Alu*J, *Alu*S, Table S19, Fig. S27-S31) show broad methylation profiles with low CpG content and higher divergence (Fig. 5A). Unlike the other active retroelement families whose full-length elements show high PRO-seq signal (purple lines in parallel plots, Fig. 5B-E), full length *Alu*Y elements show the full diversity of transcriptional activity, likely influenced by local chromatin/epigenetic features of surrounding insertion sites, which appear heavily methylated.

**Figure 5.**
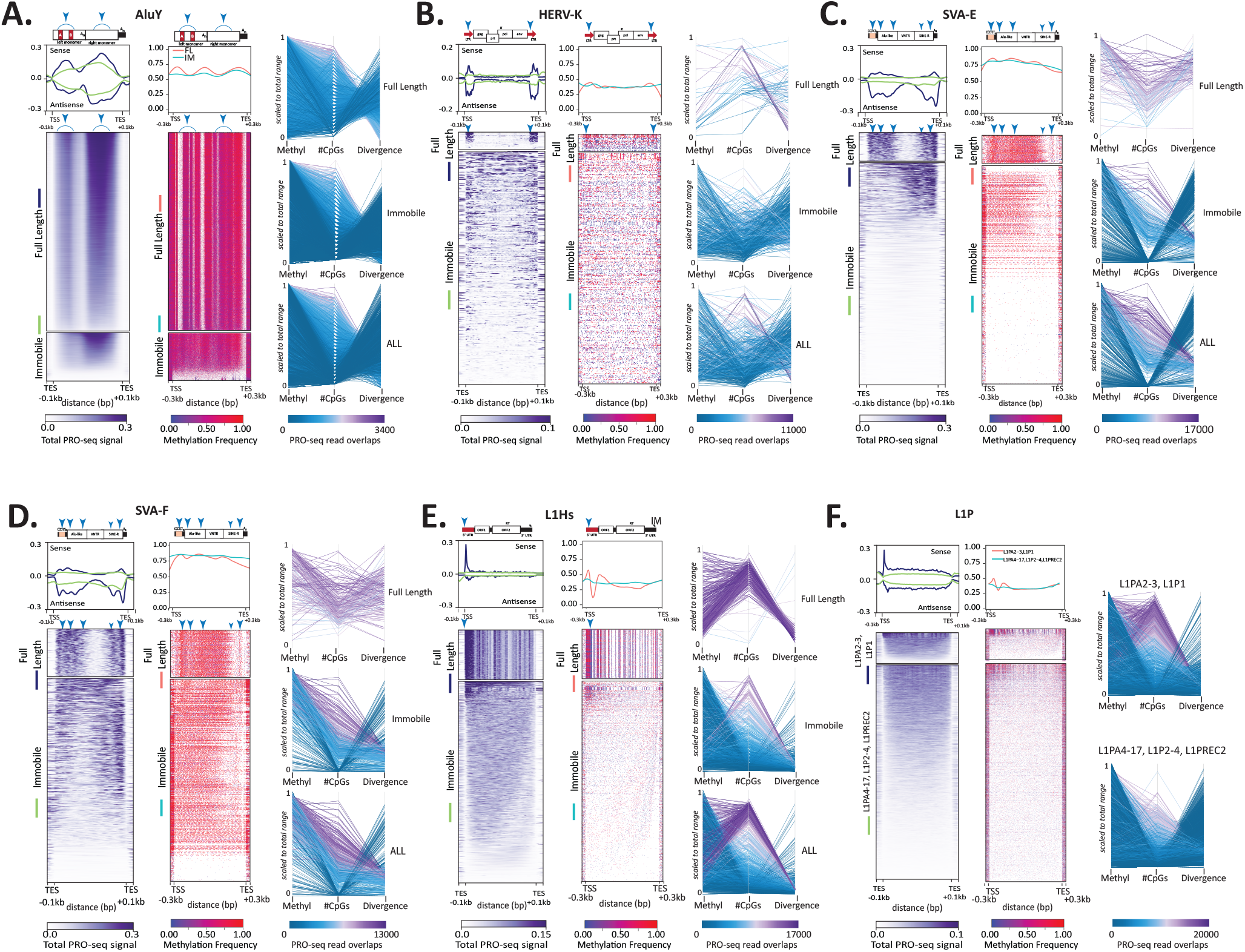
Transcriptional profiles of TEs are highly correlated with sequence divergence and epigenetic features. **(A-F)** RNA polymerase occupancy, methylation levels, CpGs and divergence for **(A)** AluY, **(B)** HERV-K, **(C)** SVA-E, **(D)** SVA-F, **(E)** L1Hs, and **(F)** L1P elements from CHM13. Heatmaps of (left panel) CHM13 PRO-seq density (red scale, normalized reads per million) and average profiles showing sense and antisense strands (upper panels) and (right panel) methylated CpGs (red-purple scale, aggregated frequency per site) for TEs grouped by their potential for mobility in A-E (full length (FL) and immobile (IM) or L1PA subfamily (F, all immobile)). Each repeat element is scaled to a fixed size; TSS (transcription start site), TES (transcription end site), and ±0.1Kbp or ±0.3Kbp are shown (bottom). Representative schematic of elements and respective subcomponents are shown above the composite profile, scaled to the TSS and TSS; red blocks indicate previously known promoter regions. Blue arrows indicate sites of RNA polymerase pile-ups. (Right of each set) Parallel plots for each TE are shown, highlighting each group of TEs (FL/IM, or L1P subfamily). Vertical axes represent scaled values for average methylation, # of CpG sites, and divergence from RepeatMasker consensus sequences for each instance of the element. Coloration by the number of overlapping ProSeq reads where purple represents the highest read overlap and blue the lowest, on the scale matching each plot.

HERV-K elements, the remaining active LTR-family of retroelements in the human genome (*68, 69*), show clear peaks of RNA polymerase signal over the LTRs, which contain strong, bidirectional promoters (*70*). While PRO-seq signal is still detected in truncated HERV-Ks, there is no peak discernible in the LTRs despite their retention, indicating they are targeted for silencing (*71*). Full length HERV-Ks that are highly expressed have generally low methylation levels despite high CpG content (Fig. 5B, Fig. S27-S29). SVA_E and SVA_F elements are the only SVA elements in the human genome that retain mobility (*72, 73*) and both show similar PRO-seq peaks (Fig. 5C, D) that distinguishes them from their inactive counterparts SVA_A-D (Fig. S27-30). While no known promoters have been previously reported for SVA elements, we find evidence for RNA polymerase promoter proximal pausing at the 5’ end of the element at predicted TSS (*74*). Interestingly, we find PRO-seq peak signal at the 3’ end within the HERV-K/LTR5a derived portion of the element, overlapping with the Kruppel-associated box (KRAB)-containing zinc finger proteins (KZFPs) controlled enhancer activity (TEEnhancer) identified in this region (blue arrows Fig. 5C, D) that contributes to human-specific early embryonic transcription (*75*). While some SVA_F truncated elements retain the 5’ promoter signal, most SVA elements retain the 3’ signal (Fig. 5C, D, Fig. S27, S29-S30) and thus, while immobile, may also retain the ability to modulate gene expression.

L1Hs elements, considered a major contributor to human structural variation (*76*), show a strong promoter-proximal pause signal at the 5’ end, where the promoter resides (*77*)(Fig. 5E). This location is also the site of high methylation levels, both of which define full length elements (Fig. 5E, Fig. S27-S31). Full length L1Hs elements show a clear trend of lower methylation levels despite high CpG content, including a hypomethylated TSS. As elements become inactivated through 5’ truncation (*78, 79*) and increased divergence, CpG content drops considerably. (Fig. 5F, S27-S31). This is exemplified in L1Ps (Fig. 5F), which show a concomitant shift to higher methylation frequencies and lower expression profiles (Fig. S27-S31), indicating that CpGs are targeted for methylation and subsequent deamination from cytosine to thymine.

### The transcriptional landscape of human centromeres

The availability of high confidence centromere annotations for CHM13 (*13, 80, 81*) provides an unprecedented opportunity to assess transcription and active RNA polymerase activity across the centromere and pericentromere. Recent work implicates centromere transcription as integral to proper centromere function, impacting the pivotal event in centromere assembly: the loading of newly synthesized CENP-A histones (*82*–*88*). While mounting evidence suggests RNA is a critical component of the epigenetic cascade leading to faithful CENP-A assembly, an assessment of nascent transcription across human centromeres has not been possible before the complete T2T-CHM13v1.0 assembly.

Across all *genome-dependent* and *genome-independent* approaches (Note S8, Fig. S32-S33), we observed low levels of satellite transcription (Fig. S34-36, Table S20), indicating that RNA polymerase occupancy at satellites, inclusive of all satellites annotated in CHM13v1.0, is lower than that observed for all other repeat types. The low levels of satellite transcription are not explained by differences in genomic abundance between satellite repeats and other repeats. Indeed, after normalizing the observed PRO-seq levels to the expected levels based on the abundance of repeats in the genome (shuffled reads), satellites are the lowest among all other repeat types (Fig. S36), indicating repression of satellites genome wide.

Since centromere transcription and CENP-A deposition are dynamic processes (*89*), we set out to test whether satellite transcription varied across the cell cycle. Following synchronization and release into mitosis (Fig. 6A, left), both CASK (Fig. 6A) and BT2 mapping (Fig. S37) methods show overall repeat transcription drops in mitosis, with the exception of tRNA transcription, which increases slightly. SINEs, LINEs and LTRs increase transcription rates at the 1hr timepoint, and reach a steady state by 1.5hrs, coincident with the transition to G1 post CENP-A loading. Notably, satellite transcripts are detected, but at low levels across the cell cycle (Fig. 6A, Fig. S37), consistent with the data obtained for CHM13 (Fig. S34-36). Given the developmental stage of CHM13 (*22*), we used publicly available datasets to determine if the low level of satellite transcription was specific to CHM13 or its early developmental stage. Across cell types (CHM13, RPE-1) and developmental stages (ES-embryonic stem cells, DE -differentiated endoderm, duodenum, ilium), retroelements show dynamic PRO-seq profiles yet satellite transcription remains low (Fig 6B, Fig. S38-S39). Across all cell types and timepoints, alpha satellites within the CENP-A containing higher order repeat arrays (HOR) (*13*) show generally higher PRO-seq signal than dHOR (<u>de</u>generate <u>HOR</u> alpha satellite arrays) and monomers or interstitial alpha satellites (MON) (Fig. S40). Thus, while nascent transcription is low, transcription from alpha satellites is detectable within the HOR domain that demarcates the active centromere (Fig 6C). The low level of detectable transcripts within the active HOR domains are in contrast to the transcriptional level of pericentromeric satellite arrays where satellite transcripts promote the recruitment of chromatin modifiers to maintain the heterochromatic status of these domains (*90*).

**Figure 6.**
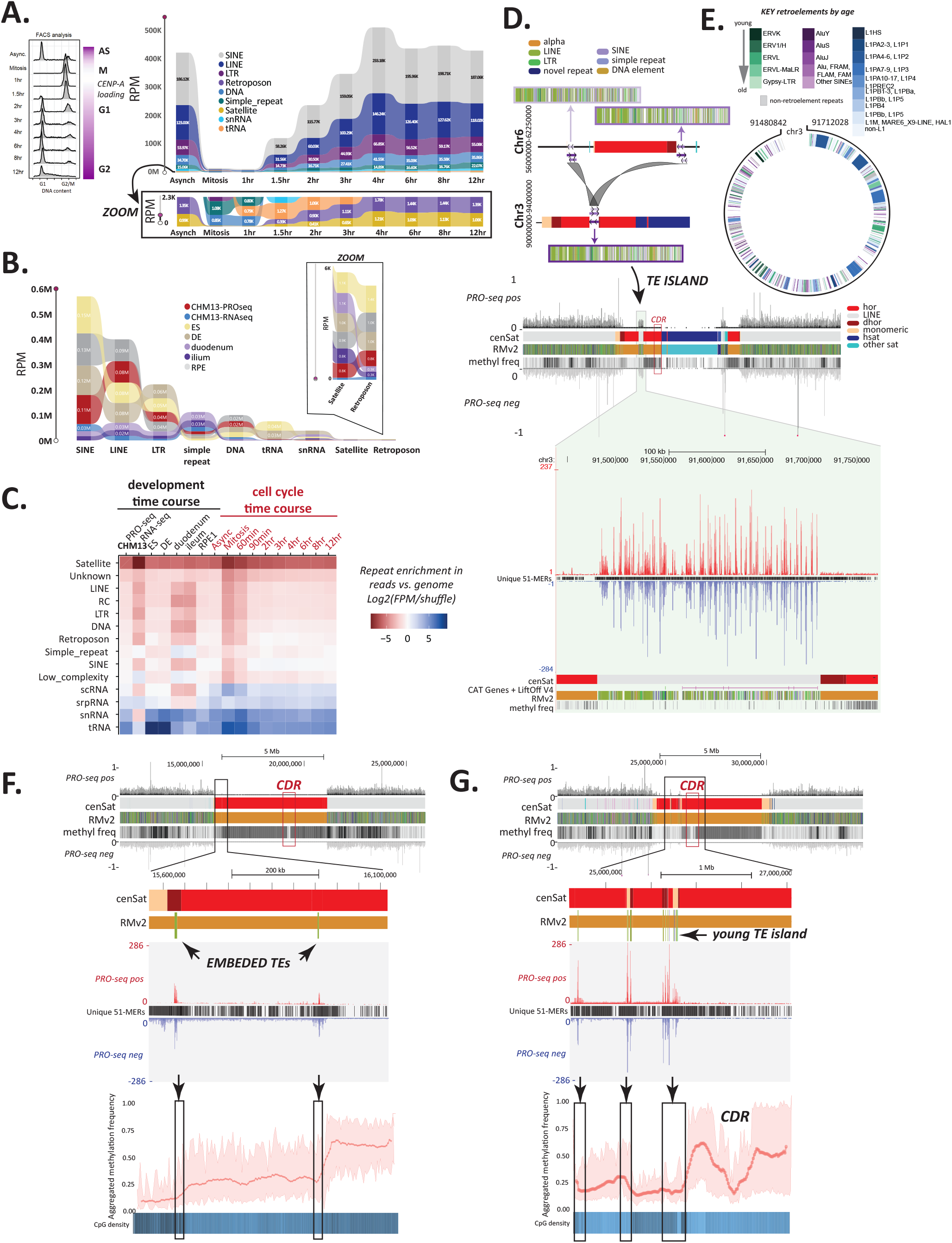
Centromere landscape is characterized by the transcription of TEs rather than satellites. **(A)** (Left) Cell sorting data showing the stages of the cell cycle following synchronization and release. (Right) ribbon plots of repeat abundance in PRO-seq data (shown as Reads per Million RPM) assessed by CASK (Classification of Ambivalent Sequences using k-mers) method in asynchronous and synchronized HeLa cells collected at time points across the cell cycle (key in inset). Zoom shows the reads for the lower range of expressed repeats, including all satellites classified in CHM13 (tan). **(B)** Ribbon plot of repeat abundance in PRO/CHRO-seq data, shown as RPM, assessed by CASK method across different developmental stages and samples. Datasets include CHM13 PRO-seq and native RNA-seq, PRO-seq for RPE-1 (differentiated retinal pigment epithelial cells), and CHRO-seq for H9 ES (embryonic stem cells), DE (differentiated endoderm cells), duodenum tissue, and ileum tissue. Zoom shows the reads for the lowest of categories of repeats across all samples, including the satellites classified in CHM13. **(C)** Repeat enrichment across PRO-seq and RNA-seq datasets (all times points and tissues) ranked from least (red) to most enriched (blue) based on k-mers normalized to genomic frequency in CHM13. **(D)** Island of TE elements found within the HOR of CHM13 Chromosome 3 has undergone segmental duplication to pericentomeres of Chromosome 6. Colored blocks are RM2 tracks for the duplications as indicated by purple (light-Chromosome 6, dark – Chromosome 3). **(D)** Browser track of CHM13 Chromosome 3 centromere including cen transition (gray) and satellite arrays (colored as per key to left). The TE island (black box) resides within the alpha satellite HOR yet does not overlap with the CDR (red box) (22). Top: censat, RM2, methylation frequency and PRO-seq (BT2 normalized to non-mitochondrial alignments), are shown. Zoom inset (right) shows PRO-seq signal across TEs (denoted as per RM2 track) associated with low methylation levels. **(E)** The relative age of retroelements (left) shows the island contains no elements with recent activity, but rather has elements that were active during the divergence of the hominoid lineages. **(F)** Recently active retroelements (green ticks in RM2 track) found embedded within alpha satellite HOR arrays (red) on Chromosome 18 are transcriptionally active (left, middle) and located (black arrows and boxes) at transitions in CpG methylation (metaplot at bottom; 200 bins total) and CpG density (below) within the array. Grey arrowhead indicates an example location of potential read through transcription from the TE into the satellite **(G)** The HOR of Chromosome 19 is interrupted by TE insertions and a “young” TE island, neither of which overlap with the CDR. The TEs mark transitions in CpG density and CpG methylation (bottom, right, color-coded as per Chromosome 18).

Early work in human cell lines showed that there was a higher propensity for recent TE insertions in HORs, with older TEs detected in dHOR and monomeric regions (*91*) (and see (*13*)). TE annotations for CHM13 show that members of the active retroelement families are found within centromeric HOR satellite arrays (Table 4, Table S21) at low frequencies. The HOR of Chromosome 3 contains a TE island adjacent to a dHOR (Fig. 6D, E), reminiscent of TE rich centromeres found in other species (e.g. maize (*92*), cereal (*93*), rice (*94*), wallabies (*95*–*97*), gibbons (*27, 58*)) and the TE islands that define *Drosophila* centromeres (*98*). The Chromosome 3 TE island is 97% similar to two pericentromeric loci on Chromosome 6 (Fig. 6D) and consists of older, non-mobile retroelements (Fig. 6E), suggesting it is the result of segmental duplication rather than serial TE insertions. This TE island, however, is transcriptionally active and demarcates transitions in methylation frequency that demarcate inactive (dHOR) and live HOR arrays (*22*). Single insertions of TEs are found within HORs, dHORs, and monomeric regions (Table S21), retain their PRO-seq signal yet show limited evidence of transcription of adjacent alpha satellites (Fig. 6F,G), indicating that read-through transcription from embedded TEs may impact alpha satellites, but not in the arrays underlying the CDR. While the generally low signal of transcription and unique marker density across HORs precluded an informative deepTools2 (*99*) analysis of PRO-seq profiles (Fig. S41), TE insertions show a marked impact on methylation trends across the HORs. For example, two insertions of L1Hs elements on Chromosome 18 mark shifts in aggregated methylation frequency (Fig. 6F). On Chromosome 19, insertions of TEs that break the HORs into dHORs and monomeric segments define transitions in DNA methylation profiles (Fig. 6G). Notably, the TE insertions within the monomeric blocks adjacent to the CDR-containing HOR form a TE boundary, and in the p-arm side of the array appear to form a young TE island derived from multiple insertion events rather than segmental duplication (Fig. S42). PRO-seq signal is higher in the alpha satellites immediately adjacent to the TE island; however, as in Chromosome 3 the signal from satellites appears limited to the dHOR and monomeric alpha satellites.

**Table 4.** TE density increases in distance from the HOR portion of centromeres. Number of TEs, by class, embedded within satellite regions, defined by location with respect to the centromere and non-centromeric satellites (13). The CENP-A region is found in the HOR.

| cenSAT<br>annotation | SINE | LINE | LTR | Retroposon | DNA |
| --- | --- | --- | --- | --- | --- |
| HOR | 5 | 19 | 1 | 0 | 0 |
| dHOR | 14 | 67 | 1 | 1 | 3 |
| Monomeric (MON) | 176 | 958 | 196 | 23 | 9 |
| HSAT1 | 462 | 0 | 0 | 0 | 66 |
| HSAT2 | 0 | 19 | 0 | 0 | 0 |
| HSAT3 | 2618 | 41 | 5 | 130 | 0 |
| HSAT4 | 16 | 4 | 6 | 1 | 0 |
| HSAT5 | 2 | 0 | 44 | 0 | 0 |
| BSAT | 248 | 32 | 99 | 6 | 4 |

Given the higher proportion of L1Hs insertions in HORs, and previous work showing a strong link between L1 transcription and neocentromere formation (*88, 100*), we compared L1Hs embeds within HORs to those found in dHORs, monomers and chromosome arms to determine whether L1Hs embeds retained their TE signatures or were “overwritten” by their local environment. We find no statistical evidence that L1Hs embedded within HORs and dHORs deviate in length, divergence, or average methylation from those found outside of these regions (Fig. S43-S44, Table S22). However, L1Hs embedded within monomeric segments of alpha satellites are both more diverged and less methylated when compared to L1Hs that are embedded in HORs (p<0.05), dHORs (p<0.01), or not embedded at all (p=<0.001). In addition, both L1Hs and *Alu*Ys within monomeric regions show less transcription than their counterparts elsewhere in the genome, including those in the HOR and dHOR (Fig. S41, S44; it should be noted that the number of embeds within the HOR for *Alu*Y elements is too low to delineate signal from stochastic noise with confidence (Table 4)).

While we find no clear link between alpha satellite transcription and the CENP-A loading domain that coincides with the CDR (*13, 22*), transcription is detectable from embedded TEs and marks shifts in methylation frequencies across satellite domains. Whether and how TEs facilitate these shifts is unknown. In previous work, the activity and copy number of TEs has been linked to alterations in methylation levels within centromeres in interspecific hybrids, resulting in chromosome instability (*97*), indicating a balance of methylation is required for centromere stabilization. With the technological advances presented in the assembly and annotation of the CHM13v1 human reference, comparative studies across other species will aide in revealing how the structure of the satellite dense centromeres of human differs from that of TE-enriched centromere in other species (*101*) and how these differences impact centromere function and chromosome evolution.

### CHM13 serves as a reference for comparative TE analyses across human genomes

Studies of the link between TE activity and chromatin states can extend beyond local influences, as exemplified by LINE and SINE transcriptional activity and the chromosome-wide silencing of the X chromosome during X inactivation (*102*–*104*). Two noncoding RNAs on the X chromosome are central to the inactivation of one X in females, *Xist* and *Tsix (105)*. These two loci overlap one another in a sense/antisense orientation but are in distinct topologically associating domains (TADs); *Tsix* is the antisense repressor of *Xist*, whose upregulation leads to X inactivation (*106*). The bipartite structure of the locus in two TADs facilitates partitioning of the X inactivation center (XIC) and supports appropriate timing of X inactivation through *Xist* transcription in early development (*107*). Moreover, an early step in the formation of heterochromatin across the inactive X is the silencing of LINEs and SINEs within the *Xist* RNA compartment (*102*).

The scarcity of SNPs (*22*) in CHM13, coupled with the short-reads of PRO-seq data, made it impossible to discern transcripts originating from one X allele versus the other within CHM13. However, the methylation-based long-read clustering method developed for CHM13v1.0 (Note S9) afforded the ability to phase reads into their individual alleles, supporting the assessment of methylation differences of TEs between the two X chromosomes in the XIC. Targeting the *Xist/Tsix* locus, nanopore reads were clustered into two distinct alleles with differential methylation profiles (Fig. 7A), supporting the observation that X inactivation occurs in CHM13 (*22*). PRO-seq signal was found across the *Xist* locus while no signal was detected from the *Tsix* locus, indicating X-inactivation has proceeded, resulting in differential methylation profiles across alleles. Low methylation (blue in Cluster 2) marks the initiation of *Xist* transcription, followed by high methylation levels across the *Xist/Tsix* locus on this allele, inclusive of the interspersed repeats found across the locus (Fig. 7A, Table S23). A distinct pause (indicated by a pile-up of PRO-seq signal) after the termination signal of the *Xist* transcript unit was found that coincides with the TAD junction and delineates the *Xist* and *Tsix* domains. These data are inconsistent with a recent report that androgenetic hydatidiform moles lack X-inactivation based on qRT-PCR (*108*), demonstrating the utility of coupling nanopore-based methylation calls with the detection of nascent transcripts through PRO-seq for high resolution functional assessment of loci in the human genome.

**Figure 7.**
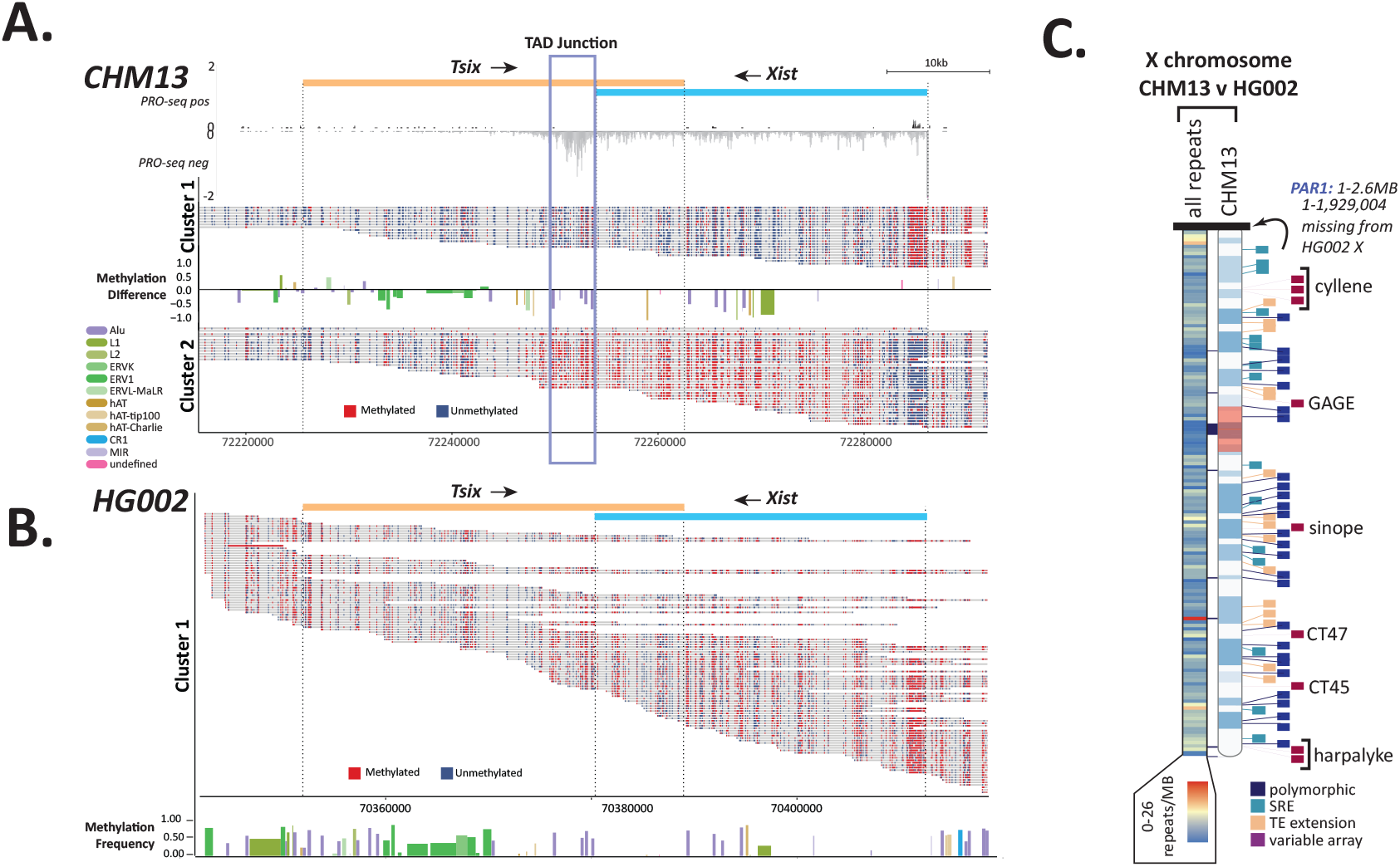
Repetitive elements define differences between human genomes. Single read methylation profiles were extracted, and reads were clustered based on the methylation state of the Xist promoter from CHM13 **(A)** and HG002 **(B)**. Differences in repeat methylation were calculated by taking the average methylation per repeat and subtracting cluster 2 repeats from cluster 1 repeats. Directionality of transcript units are indicated (top). Normalized PRO-seq reads show a marked pile-up of RNA pol II at the predicted TAD boundary at the 3’ end of the Xist transcript. **(C)** Heatmap of Chromosome X showing the location of all repeat differences between the X’s of HG002 and CHM13 (left) and the location of the top four categories of repeat differences: polymorphic (insertion/deletion), SRE (short repeat extension), TE extension, and variable array length (right ideogram). Gaps between CHM13 and GRCh38 are indicated with black blocks between the heatmap and ideogram.

As an example of the prospective that the CHM13 reference, methylation and repeat annotations provide for future work expanding to emerging long-read based human genome assemblies, we compared both the XIC and chromosome-wide repeat content of the Chromosome X from CHM13 and HG002 (XY). As expected, the XIC in HG002 shows high methylation across the locus and only a single allelic cluster (Fig. 7B, Table S23), with no detectable transcripts across the *Tsix/Xist* domain (data not shown). Sequence comparison of the 269,020 repeats assessed between the haploid X of HG002 and CHM13 (Fig. 7C, Table S24-25) uncovered 778 repeat differences, of which 70% were simple repeats and 21% were TEs (64 of which were length outliers (Note S9, Fig. S45)). Collectively, these data revealed a combination of simple repeat and TE expansions, nested TEs (polymorphic insertions in the population), and variability in the size of arrays of novel repeats identified in CHM13 (sinope, cyllene and harpalyke) and composite elements, three of which carry genes in their core unit (GAGE12, CT45 and CT47) (Fig. 7C). Of the five new arrays identified on Chromosome X, three are present in HG002; the remaining two are located in the PAR1 region which has not been fully assembled at this time (Fig. S46). Collectively, these data demonstrate that the depth of repeat annotations based on the CHM13v1.0 assembly can serve as a reference for studying human variation inclusive of repeats that impact local and regional chromatin, gene expression, and gene copy numbers.

### Conclusion

The assembly of the first truly complete, telomere to telomere (T2T), human genome reference facilitated our development of an unprecedented atlas of repeats that comprise over 53% of the human genome. Through this collaborative effort, we have developed a resource of human repeat annotations and methods to guide future efforts in exploring the complexities of repeat biology in human and other genomes. Our efforts focused on two main areas: repeat *sequence* and *functional* annotation. Through repeat sequence annotations, we have updated repeat models and implemented newly developed repeat modelling tools that supported the identification of previously unknown satellite arrays, expanded the catalog of variants for known repeats and TEs, and identified new classes of complex, composite repeat elements. Deeper exploration of such repeats, highlighted by examples herein, revealed the complexity of genetic mechanisms that impact repeats during different phases of their lifecycle and thus illustrate the myriad mechanisms by which they are major contributors to defining the structure and content of the human genome.

For example, we found that a single TE transduction event captured a short sequence, WaluSat, in a primate ancestor. Subsequent segmental duplications of the region carrying this new composite TE-sat repeat spread the sequence to several chromosomes, including four of the acrocentric chromosomes. The satellite portion of the repeat expanded to almost 0.5Mbp of sequence on the acrocentrics, resulting in the alteration of the structure of this portion of the chromosome into regions dense with G4s, structures recently suggested to be novel functional elements within the human genome (*109*). This example highlights the need for future functional studies dissecting the impact of repeats on the local chromosome environment, such as replication timing, local transcription, DNA damage and repair processes, and establishing TAD boundaries. Correlatively, these data lay the groundwork for exploring the impact of local environments (such as gene poor regions as found on the acrocentric arms of human chromosomes) on sequence constraint and mutation rates for newly established repeats.

Our functional annotations defined the transcription of repeats, including sites of engaged RNA polymerase, as well as DNA methylation profiles based on long-read sequencing, both at single nucleotide resolution. These data collectively provided the first high-confidence functional annotation of repeats across the human genome. Using these data, phylogenetic and statistical modelling exposed differential evolutionary patterns across TE families and among chromosomal regions. For example, we find that the tandemly arrayed TE-derived satellite SST1 carries unique methylation and transcriptional profiles, including an enhancer embedded within each unit, that is found only in specific arrays on Chromosomes 19 and 4. These arrays are already known to be hypervariable in the human population and alterations in their activity have been linked to several cancers. However, a full understanding of copy number variation, epigenetic instability and transcription of SST1 elements has been hampered by a lack of complete annotations of copies of these elements elsewhere in the genome. In this study, we doubled the annotations of SST1 sequences across 17 chromosomes (including the Y chromosome from the GRCh38 assembly), providing a reference to define clinically relevant loci with respect to chromosome instabilities that often accompany cancer transformation. Our functional annotation revealed transcriptional signatures of both promoters and enhancers within active SST1 elements that would impact local transcription and chromatin structures. Moreover, the detection of an enhancer implicates SST1 in defining cellular partitions, such as paraspeckes and phase separated condensates ((*63*), reviewed in (*110*)), that would have an impact on other genomic loci.

Combined with the work of defining the linear order and content of centromeric sequences as part of the T2T consortium, we find that engaged RNA polymerase signal is low across centromeric satellites arranged in arrays, irrespective of stages of the cell cycle or development. Rather, active transcription is detected in embedded retroelements coinciding with shifts in methylation states that demarcate active centromere domains. To date, the centromere biology field has been limited by a lack of a linear assembly across human centromeres, challenging the development of models to describe genetic and epigenetic elements that define centromeric chromatin. Given the recent work focused on satellite transcription and centromere function, our data, in concert with the extensive centromere annotations for CHM13 (*13*), reveal that these high density repeat regions are not static in sequence, epigenetic nor transcriptional activity and that there is a high degree of substructure across the centromeric regions that impact function. This work lays the foundation to support functional studies that aim to tease apart different transcriptional and epigenetic components that differentiate centromere assembly, maintenance and initiation within a centromere and across centromere variants. Expanding this work to compare the landscape of the variable centromere forms that are found across all domains of life, as well as in those that arise in human disease, will reveal the complex life cycle of centromeres (*101*).

Studies of human genetic variation have been relatively blind to repeat variation among individuals, particularly arrayed and complex repeats, as these types of sequences are recalcitrant to short read sequencing technologies, mapping and functional annotation methodologies. As a prospective of the utility of complete reference genomes in studying human genetic variation, we compared two T2T X chromosomes. We find 218Kbp of repeat differences among these two chromosomes (0.18% of the chromosome, excluding the 1.9Kbp PAR), including repeat variation in complex arrays that carry exonic material and thus affect gene dosage. Thus, comparative analyses of T2T-level assemblies reveal the potential for discovering an even wider range of repeat variation across the 46 chromosomes that constitute the human genome.

Finally, our work demonstrates the need to increase efforts towards achieving T2T-level assemblies for non-human primates to fully understand the complexity and impact of repeat-derived genomic innovations that define primate lineages, including humans. For example, we find repeat variants that appear enriched or specific to the human lineage that may impact gene content, such as protein coding sequences found in composites and those derived from TE transduction events. In the absence of T2T-level assemblies from other primate species, we cannot truly attribute these novelties to specific human phenotypes; the efforts of the T2T consortium lay the groundwork for such in-depth comparative analyses. The extent of variation described herein for a single human chromosome across two individuals (HG002 and CHM13) highlights the need to expand the effort to create human and non-human primate pan-genome references to support exploration of repeats that define the true extent of human variation. Notably, these discoveries and tools can be brought to bear on the challenge of establishing breeding programs for critically endangered species across all forms of life (*111*), for which fragmented genomes hamper an understanding of the true population-level genetic variability in species with extremely low population sizes.

## Acknowledgments

We would like to thank the UConn Computational Biology Core for computational support. We thank the NIH Intramural Sequencing Center and the UConn Center for Genome Innovation for sequencing. This work utilized the computational resources of the NIH HPC Biowulf cluster (https://hpc.nih.gov),the UConn HPC Xanadu Cluster (https://bioinformatics.uconn.edu; https://health.uconn.edu/high-performance-computing), the Stanford Research Computing Center, the Stanford Sherlock HPC cluster, and the PALMA-II HPC cluster of the University of Münster (https://confluence.uni-muenster.de/display/HPC). Special thanks to Mark Diekhans for assistance with UCSC browser tracks and liftOver chain development.

## Funding

This work was supported, in part, by the Intramural Research Program of the National Human Genome Research Institute, National Institutes of Health (AMP). Grants from the U.S. National Institutes of Health: R01GM123312-02 and R21CA240199 to SH, GAH, PGSG, RJO; R01HG002939 and U24HG010136 to JS, JR, AFAS; R01HG009190 to AG, WT; R01HG00990905 to CL, AFS; R35GM128857 to LW, LJC; R01GM132600, P20GM103546, and U24HG010136 to DO, TJW; U24HG010263, U24HG006620, U01CA253481, R24DK106766-01A1 to MCS; R01HG002385, R01HG010169 and U01HG010971 to MV, EEE; R01 1R01HG011274-01, R21HG010548-01, and U01HG010971 to KHM; National Science Foundation: NSF 1613806 and NSF 1643825 to RJO; NSF DBI-1627442, NSF IOS-1732253, and NSF IOS-1758800 to MCS; Connecticut Innovations: 20190200 to RJO; Mark Foundation for Cancer Research: 19-033-ASP to MCS; Stowers Institute for Medical Research: LGdL, JLG; Howard Hughes Medical Institute Hanna H. Gray Fellowship: NA; EEE is an investigator of the Howard Hughes Medical Institute.

## Author contributions

Repeat annotation pipeline implementation: SJH, JS, JR, TJW, AFAS, and RJO Repeat manual curation: SJH, JS, GAH, PGSG, JR, AFAS, DO, TJW, and RJO Dfam Database Update: JS, JR, and AFAS

ULTRA data analyses: DO and TJW Transduction analyses: RH, MR, and WM

PRO-seq analyses: SJH, CL, LW, AFS, LJC, and RJO

PRO-seq data production: CHM13, RPE: SJH; HeLa: LW CASK pipeline: CL and AFS

Methylation analyses: AG and WT

Centromere Sat Team data integration: NA and KHM Segmental duplication and dot plot analyses: MV and EEE Assembly Team Lead: AMP

Centromere Annotation Team Lead: KHM Variant Team Lead: MCS

Methylation Analysis Team Lead: WT Repeat Analysis Team Lead: RJO Segmental Duplication Team Lead: EEE

Data Visualization: RJO, GAH, SJH, PGSG, and AG Figures: RJO

Project administration: SJH and RJO

Supervision: LJC, JLG, NA, WM, AFAS, AFS, TJW, MCS, EEE, AMP, WT, KHM, and RJO

Manuscript draft: RJO and SJH.

Supplement Draft: SJH, GAH, PGSG, JS, and RJO.

Editing: SJH, RJO, JS, TJW, AFS, WM, JLG, GAH, PGSG with the assistance of all authors.

## Competing interests

KHM has received travel funds to speak at symposia organized by Oxford Nanopore. WT has two patents (8,748,091 and 8,394,584) licensed to Oxford Nanopore Technologies. All other authors declare that they have no competing interests.

## Data and materials availability

*Sequencing data and assemblies (NCBI BioProject PRJNA559484):*

https://www.ncbi.nlm.nih.gov/bioproject/559484

*Sequencing data, assemblies, and other supporting data on AWS:*

https://github.com/marbl/CHM13

*Assembly issues and known heterozygous sites:*

https://github.com/marbl/CHM13-issues

*UCSC assembly hub browser:*

genome.ucsc.edu/cgi-bin/hgTracks?genome=t2t-chm13-v1.0&hubUrl= http://t2t.gi.ucsc.edu/chm13/hub/hub.txt

*Dot plot visualization and browser:*

https://resgen.io/paper-data/T2T-Nurk-et-al-2021/views/t2t-identity

*T2T Consortium homepage:*

https://sites.google.com/ucsc.edu/t2tworkinggroup

*Repeat Library for New Repeat Entries:*

https://gitlab.com/SJHoyt/t2t_transposable-elements/Repeat_annotations/Repeatmasker_and_polishing/RepeatLibrary_NewRepeatEntries.embl

All data, code, and materials used in this study are outlined in Supplementary Materials.

## Supplementary Materials

SupplementaryMaterials.pdf Notes S1-S9

Figs. S1 to S46

Tables S1 to S25

References (*1*–*57*)

## Notes

https://github.com/marbl/CHM13

https://www.ncbi.nlm.nih.gov/bioproject/559484

https://github.com/marbl/CHM13-issues

https://resgen.io/paper-data/T2T-Nurk-et-al-2021/views/t2t-identity

https://gitlab.com/SJHoyt/t2t_transposable-elements

