## Supplemental Material for "From telomere to telomere: the transcriptional and epigenetic state of human repeat elements"

**Tables:**

Additional Files: Supplementary Tables.xlsx Tables S1-S3, S5, S7-18, S20-23, S25

**1. Data Availability** 2

**2. Repeat Annotations** 3

Discovery of new repeats models with RepeatModeler & loci identification with RepeatMasker 3

Annotation of tandem repeats and satellites 5

GAP identification 5

Manual curation of new repeat models 5

Composite Elements 6

Compilation and polishing of Repeat Annotationv2 6

Reverse liftOver analysis and repeat fasta comparison 9

Copy Number Comparison across primates 12

Methylation Metaplots 13

Dot Plot Analyses 13

**3. Phylogenetic Analyses** 21

TELO_Comp Phylogenetic Analyses 21

WaluSat and WaluSx Phylogenetic Analyses 22

*SST1 Phylogenetic Analyses* 22

**4. Transduction Analyses** 26

Discovery of 3' and 5' transductions in CHM13 26

Validating and categorizing the putative transductions 26

Functional annotation of transduction events 28

G-quadraplex (G4) analysis 35

**5. Precision Run-on Sequencing (PRO-seq) and RNA Sequencing analyses** 35
Cell permeabilization for CHM13 and RPE-1 35
Illumina Library Preparation for CHM13 and RPE-1 35
Pre-processing and mapping of CHM13 and RPE-1 PRO-seq data 35
Mitotic Synchronization and Release for HeLa time course 40
Cell Cycle Analysis for HeLa time course 40
Cell Permeabilization for HeLa time course 41
Library Preparation for HeLa time course 41
Pre-processing and mapping HeLa time course data 41
H9 ChRO-seq data availability and pre-processing 41
Pre-processing, mapping and post-processing of RNA-seq data 41
**6. Statistical analyses and data visualization** 42
**7. Identification of full length and immobile TEs; TE aging** 42
**8. Centromere Transcription Analyses** 52
Mapping dependent PRO-seq analyses 53
CASK: Classification of Ambivalent Sequences using k-mers 53
RepeatMasking of PROseq and RNAseq reads 54
TE Embeds within cenSAT annotations 54
Detecting Repeat Transcription - Method Comparisons 54
**9. Repeat comparisons between CHM13v.10 and HG002** 69
Methylation clustering 69
chrX liftOver analysis and repeat fasta comparison 69
**References** 73

### 1. Data Availability

| | |
|---|---|
| Repeat Library for New Repeat Entries | https://gitlab.com/SJHoyt/t2t_transposable-elements/Repeat_annotations/Repeatmasker_and_polishing/RepeatLibrary_NewRepeatEntries.embl |
| Sequencing data and assemblies (NCBI BioProject PRJNA559484): | https://www.ncbi.nlm.nih.gov/bioproject/559484 |
| Sequencing data, assemblies, and other supporting data on AWS | https://github.com/marbl/CHM13 |
| UCSC assembly hub browser | http://genome.ucsc.edu/cgi-bin/hgTracks?genome=t2t-chm13- |

| | |
|---|---|
| | v1.0&hubUrl=http://t2t.gi.ucsc.edu/chm13/hub/hub.txt |
| RepeatMasterv2 Track CHM13v1.0 | http://t2t.gi.ucsc.edu/chm13/dev/t2t-chm13-v1.0/downloads/t2t-chm13-v1.0.rmskV2.bigBed |
| RepeatMaskerv2 Track GRCh38 + chrY | http://t2t.gi.ucsc.edu/chm13/dev/GRCh38/downloads/GRCh38-rmskV2.bigBed |
| RepeatMaskerv2 Track HG002 chrX | http://t2t.gi.ucsc.edu/chm13/dev/HG002-X-v1.0/downloads/HG002-X-v1.0.rmskV2.bigBed |
| RepeatMasterv2 Track Composites CHM13v1.0 | http://t2t.gi.ucsc.edu/chm13/dev/t2t-chm13-v1.0/downloads/t2t-chm13-v1.0.composite-repeats-singletons-arrays.bed.gz |
| RepeatMasterv2 Track new satellites and arrays CHM13v1.0 | http://t2t.gi.ucsc.edu/chm13/dev/t2t-chm13-v1.0/downloads/t2t-chm13-v1.0.new-satellites-monomers-arrays.bed.gz |
| PRO-seq CHM13/RPE-1 | PRJNA559484 |
| PRO-seq HeLa | GSE179576 |
| RNA-seq CHM13 | PRJNA559484 |
| CHM13 Meryl 21-mers and 51-mers | https://s3-us-west-2.amazonaws.com/human-pangenomics/index.html?prefix=T2T/CHM13/assemblies/alignments/marker/ |
| Scripts and code used herein | https://gitlab.com/SJHoyt/t2t_transposable-elements |

### 2. Repeat Annotations

#### Discovery of new repeats models with RepeatModeler & loci identification with RepeatMasker

To assess previously unannotated repetitive regions of the genome, a RepeatMasker4.1.2-p1 run was completed on the CHM13v1.0 assembly using the Dfam 3.3 library (*1*) with the following settings: sensitive setting (*-s*), using the species tag of human (*-species human*) and the NCBI BLAST-derived search engine RMBlast (*-e ncbi*): *$ RepeatMasker -s -species human -e ncbi.* These regions (RA1a) were then hard-masked, producing “hard masked genome 1”, HM1, Fig. S1. A RepeatModeler2.01 analysis was performed on the remaining (unmasked) regions. The output file, Novel_Modeler_Repeats.fa, was run through LTR Harvest (*2*) (accessed from genometools/1.5.10) and Transposon PSI (v08222010) (*3*) to further refine novel repeat calls. As the RepeatModeler2.01 algorithm implements a random sampling of the genome, the consensi generated from RepeatModeler2.01 (All_Novel_repeats.fa) were used as a library for a secondary RepeatMasker run to collect all associated instances for each model generated on the CHM13 genome assembly (RA1b).

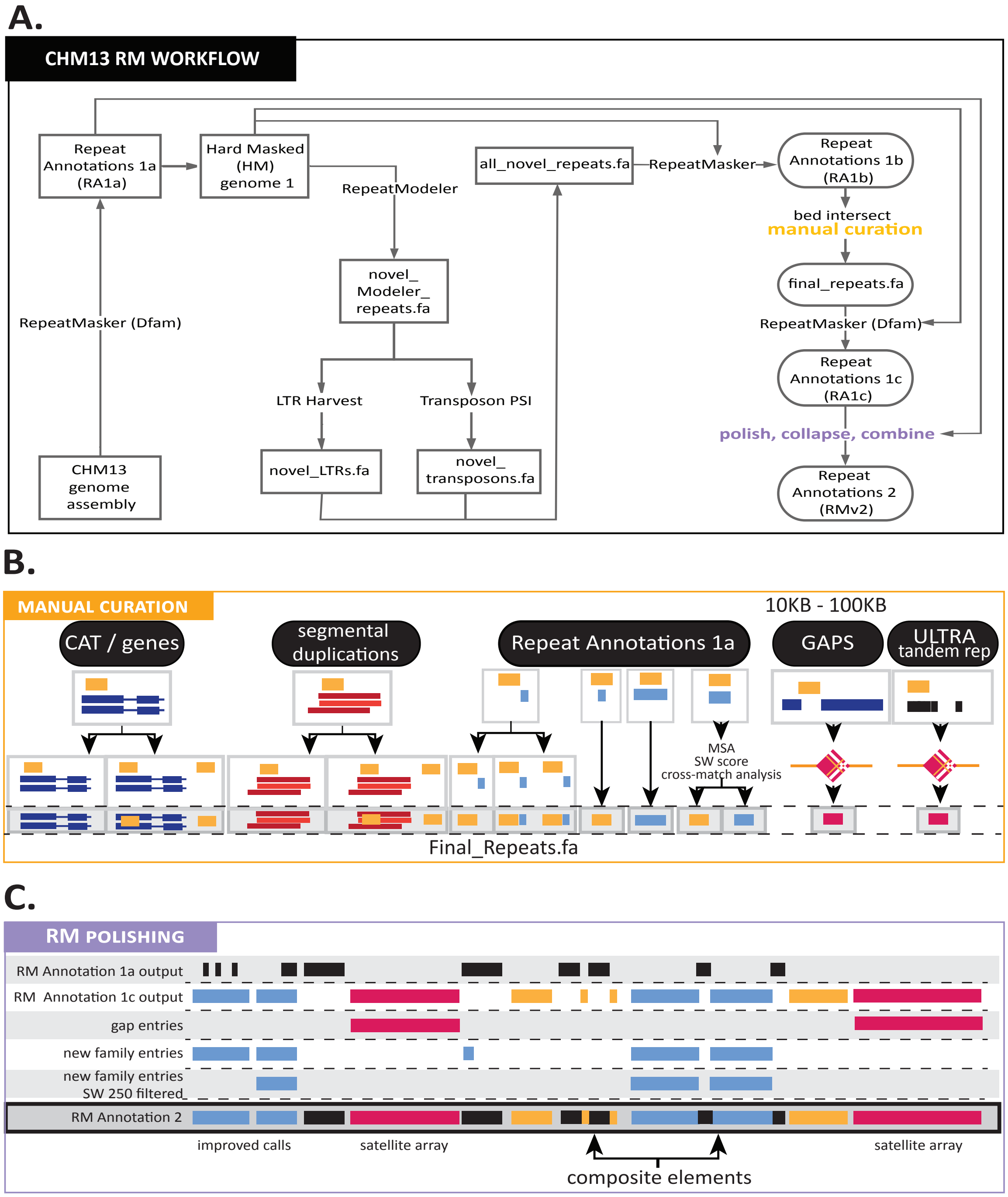


***Fig. S1. A novel workflow afforded comprehensive annotations of a complete human genome.*** *Workflow implemented to obtain updated repeat models and the derivation of RepeatMasker Annotations 2 (RMv2), consisting of compiled and polished RM annotations submitted to Dfam* ***(A)*** *and applied to CHM13 and GRCh38 as RepeatMaskerv2 tracks. Workflow consisted of multiple iterations of RepeatMasker and RepeatModeler (A). The components intersected during manual curation* ***(B)*** *include CAT/gene annotations (4), segmental duplications (5), repeats masked using Dfam (ver3.3) repeat models (1), tandem repeat arrays identified as gaps in annotations >10Kbp and overlap with ULTRA tandem repeat models (6).* ***(C)*** *Repeat model polishing was derived from a compilation of repeat masker*

*output (previous repeat models; HM1), repeat masker 2 output (updated models; RM Annotation 2), and gap entries. New family entries identified from RMv2 were further filtered following multiple sequence alignment (MSA) among members of the predicted category.*

### Annotation of tandem repeats and satellites

Tandem repeats and satellites were initially annotated in the above RepeatMasker run, based on a combination of alignments to satellite sequences in the RepeatMasker library and de novo repeat identification with Tandem Repeats Finder (TRF) v4.09 GUI version at https://tandem.bu.edu/trf/trf.html using standard parameters (*7*). This workflow left large sections of the genome unannotated, either because repeats within the sequence were too decayed to be recognized by TRF, or because the satellites were not contained in RepeatMasker's database. We expanded annotation coverage of missing repetitive regions using ULTRA (*6*), an open-source tool that can annotate and provide statistically consistent scoring for very large repeat units (up to a repeat period of 4000), arbitrarily-long repetitive regions, and ancient repeats that have highly decayed repetitive signals. ULTRA v1.0 was run with the settings: *$ ultra -mi 2 -md 2 -p 4001 -mu 2 -ws 90000 -os 10000*, respectively impacting maximum number of insertions (-mi), deletions (-md), and repeat periodicity (-p), minimum number of repeat units (-mu), and window size and overlap (-ws and -os).

### GAP identification

Gaps in the CHM13 repeat annotation were identified via bedtools v 2.29.0 (*8*) by subtracting RA1a and RA1b (Fig. S1) from the whole CHM13 genome sequence. The resulting regions were then filtered for size (only gaps larger than 5Kbp were considered). These gaps were manually curated in a UCSC Genome Browser session to check for any feature annotation overlap. Tandemly repeated sequences for each gap were identified with a combination of TRF v4.09 (*7*) and ULTRA (*6*). Additionally, self-alignment plots generated in YASS (*9*) confirmed sequence repeat monomer length. Monomers and full tandem arrays of these newly identified regions were compared via alignment in MAFFT v7.471 (*10*) and Geneious v2019.1.3 to check for any possible overlap with either the new regions or previously annotated tandem repeats; only previously unknown repeats were kept and classified as "new array/monomer".

### Manual curation of new repeat models

Following the production of RA1b, curation steps were implemented to refine new repeat models and produce RA1c (Fig. S1A, S1B). Overlaps with CAT/gene annotations (*4*), segmental duplications (*5*), and tandem repeats found within GAPs and through ULTRA overlaps were manually curated. Multiple sequence alignment (MSA) plots of the transposable element (TE) instances from RA1a aligned to the putative new repeat consensi from RA1b were used to determine the divergence, and therefore the overall age of the TE family. Highly diverged, older sequences were set aside for later assessment, as these sequences may correspond to old fragments of known TEs. This hypothesis was reinforced following a cross_match analysis of the older consensi to the Dfam database consensi. In addition, repeats that failed to match a known repeat, even distantly, were assessed by evaluating 100 nt on the 5' and 3' flanking region of the instances contributing to the initial RepeatModeler consensi. RepeatMasker was used to assess the flanking regions to determine if the neighboring sequence matched consistently to known repeats and were saved for later evaluation as possible ancestral repeats (Table S1). To confirm the set of new families (Table 1, Table S2) had not been previously defined in the Dfam database, we performed a cross_match analysis of the consensi corresponding to the new families to the curated consensi in the Dfam database. In addition, a cross_match self-comparison of the new consensi was performed to assess possible array sequence structures or intra-library duplicates. Intra-library duplicates and matches to the Dfam database were removed. The identification of composite subunits was accomplished through assessment of genome-wide instances with Circa and pattern recognition in both the UCSC browser and RM output. Bedtools closest v2.29.0 (-k 2 -iu -D ref) and (-k 2 -id -D ref) was also used to assess the neighboring repeats and their frequency increasing the likelihood that they were part of a larger repeat, or composite. This curation led to the generation of a repeat library (RMv2) (final_repeats.fa) of newly identified repeats, satellites monomers, variants of previously known/classified satellites, subunits of composites and composite elements.

**Composite Elements**

We defined a composite element as a repeating unit consisting of three or more repeated sequences, including TEs, simple repeats, novel repeat annotations and/or satellites, that is found in at least one location in the genome as a tandem array. A composite subunit is a novel repeat annotation that is most often found within a composite in a consistent pattern. Note that a composite subunit repeat may be found outside of the composite, but it is not common. Segmental duplications (SDs) were called for CHM13v1.0 using a 1Kbp cutoff(*5*); while the location of some composite elements within a family are present as a single copy and thus are likely SDs derived by non-allelic homologous recombination (NAHR)(*11*), a composite family is distinguished by the presence of composite elements in an array in at least one location, thus falling into a “megasatellite” classification (*12*).

**Compilation and polishing of Repeat Annotationv2**

The new models discovered as a result of the RepeatModeler2 analysis contained pieces of simple repeats and small pieces of previously defined TEs. As such, a RepeatMasker analysis performed by simply adding the new entries alongside previously annotated TE/repeat models in a library resulted in a large number of false positives. Therefore, a pipeline was developed (Fig. S1C) to combine the new entry annotations and the previously generated TE models in the Dfam database to produce a high confidence repeat masker annotation track for CHM13. A third RepeatMasker run was performed on HM1 using a library which included the Dfam database plus all new entries resulting in Repeat Annotations 1c (RA1c). RA1c was then combined with RA1a (Dfam library only) and the resulting combined outputs were intersected with gap entries (“new array/monomer”) and new family entries. New family entries were filtered for elements with high confidence based on MSA plots and a SW score of 250. These combined efforts resulted in the production of a final CHM13v1.0 RepeatMaskerv2 track (RMv2) for the UCSC genome browser.

Following development of RMv2, unit-length and composite unit genomic instances were determined by performing a self-comparison via cross_match to determine the maximum SW score for a particular TE model. A score conservatively lower than the determined maximum was then used in the alignAndCallConsensus.pl program (-sc #) to align the TE instances to the consensus. The resulting MSA were used for the classification of composite repeats and subunits therein (*1*, *13*, *14*) (https://github.com/Dfam-consortium/RepeatModeler).

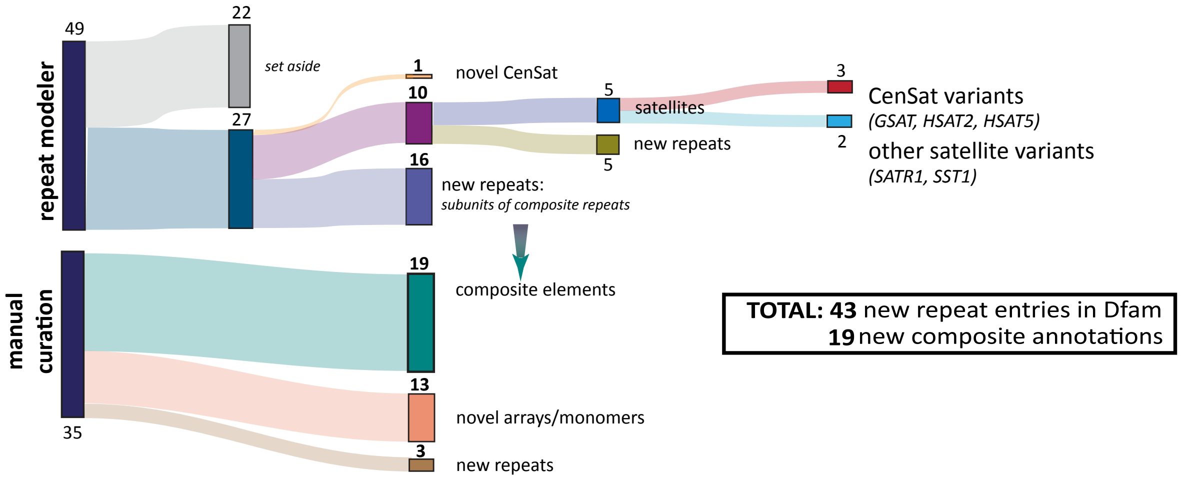


***Fig. S2*. *Summary of new repeat annotations for CHM13.*** *Compiled annotations resulted in a final RM track for CHM13 that included the annotation of new repeats and satellite arrays outside of centromeric regions (15), the identification of extensions or variants to current repeat models, and the identification of composite elements consisting of multiple repeats. Plot of 49 novel repeats identified through RepeatModeler and 35 through manual curation.*

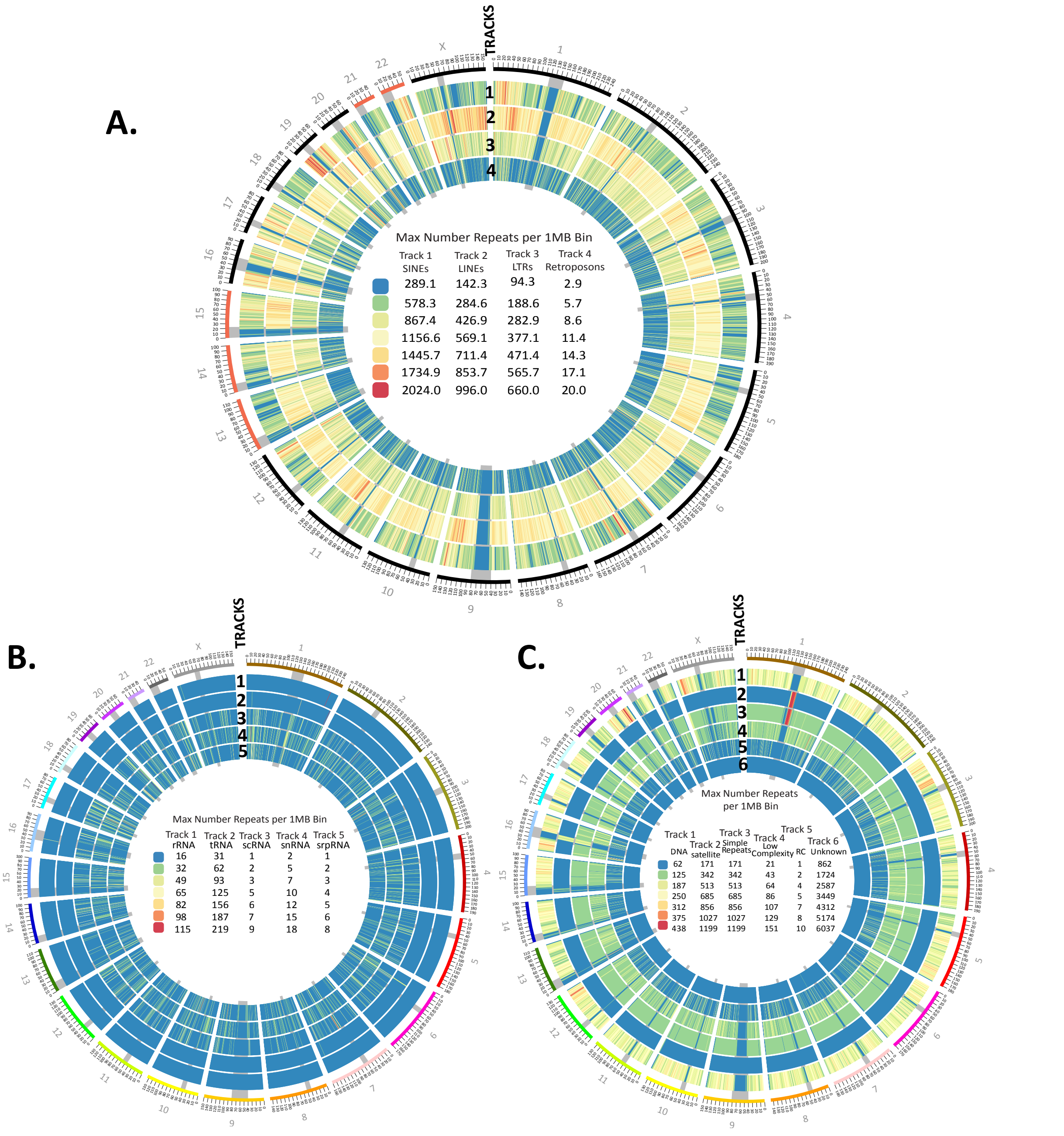


***Fig. S3. Repeat density across CHM13 by family classification.***
*Counts of all repeats identified by our repeat annotation pipeline were binned into 1Mbp windows across all chromosomes (color coded and numbered, outer ring) in CHM13 and are shown as Circos heatmaps corresponding to **(A)** retrotransposon classes, **(B)** RNAs, and **(C)** all other repeat classes. Centromere blocks (including centromere transition regions) are denoted by grey bars that span all tracks. Tracks are numbered (1, 2, …) starting from the outer ring as indicated. Each repeat class track is scaled independently with the scales located in the middle of each respective Circos. Note: the rDNAs are not included on the acrocentrics (Chromosomes 13, 14, 15, 21, 22).*

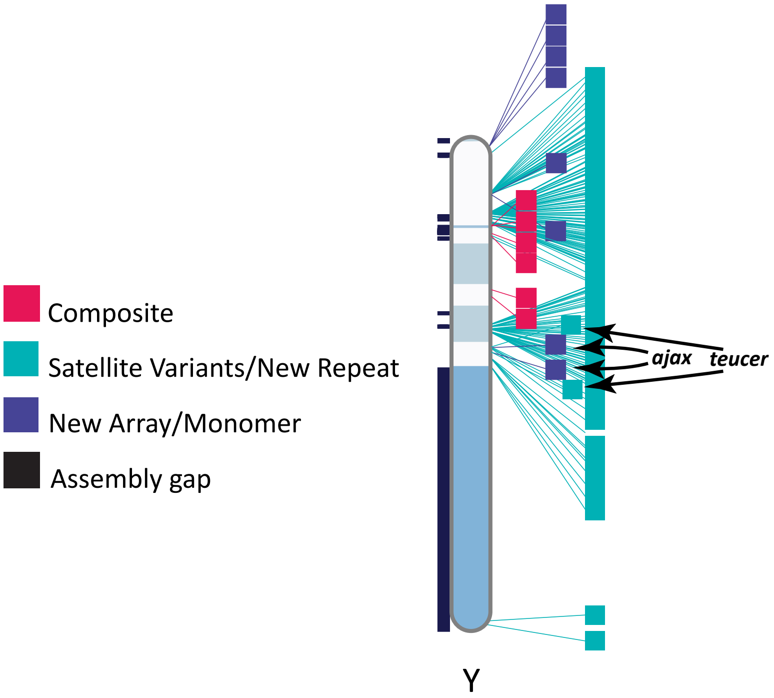


***Fig. S4. CHM13-based repeat annotations reveal new repeat classifications on the GRCh38 Y chromosome assembly.***
*Ideogram of GRCh38 Chromosome Y indicating the locations of newly annotated composite elements (red), satellite variants and novel repeats (aqua), and new arrays or monomers of sequences found within those arrays (purple). Gaps in the Chromosome Y assembly are shown in black boxes to the left of the chromosome. Notably, ajax and teucer are found together at two loci without TELO_comp as part of an inversion in the Azoospermia Factor c region of the Y chromosome, a region with recurrent de novo microdeletions linked to male infertility (16).*

### Reverse liftOver analysis and repeat fasta comparison

A bed file was generated from the CHM13 RM2 output and a reverse liftOver (*17*) performed to the GRCh38 genome assembly. The IntersectBed tool (*8*) was used with both strict (-f 0.9 -r) and permissive (-f 0.5 -r) parameters to compare the lifted GRCh38 coordinates to the GRCh38 RM-comp output. The results of the intersection were parsed based on the TE annotation match. The possible categories of the intersection analysis included: full match, class match, no match, artifacts, and of interest. A full match required, under either the strict or the loosened parameters detailed above, that both the family (e.g., *Alu*Sx) and class (e.g., SINE/*Alu*) of the CHM13 and GRCh38 RM-comp loci were identical. In the event the family differed, the intersected locus was labeled as a class match. If neither the family nor class matched, it was subsequently labeled as a no match. Loci lacking a match following intersection analysis were set aside for a further detailed direct fasta comparison as described below. Special attention was paid to intersection loci in which the locus was identified as an SVA element, as these elements contain, and are frequently mislabeled, *Alu* elements. Exceptions were made in the level of match if an SVA in the CHM13 output matched to an *Alu* in the GRCh38 RM output and vice versa.

A parallel and complementary analysis comparing the unlifted CHM13 loci fasta sequence was completed. A similarity score was assigned to each repeat based on crossmatch output as a percentage of the maximum score. Sequences with a score of greater than 90% and/or shorter than 50 bp were the threshold for concordant similarity or insufficient information for comparison, respectively. All other sequences were considered as potential polymorphic loci. The term polymorphic is used here to describe the genetic variation occurring between individuals in a population, such that each individual may contain a unique repertoire of TE insertions.

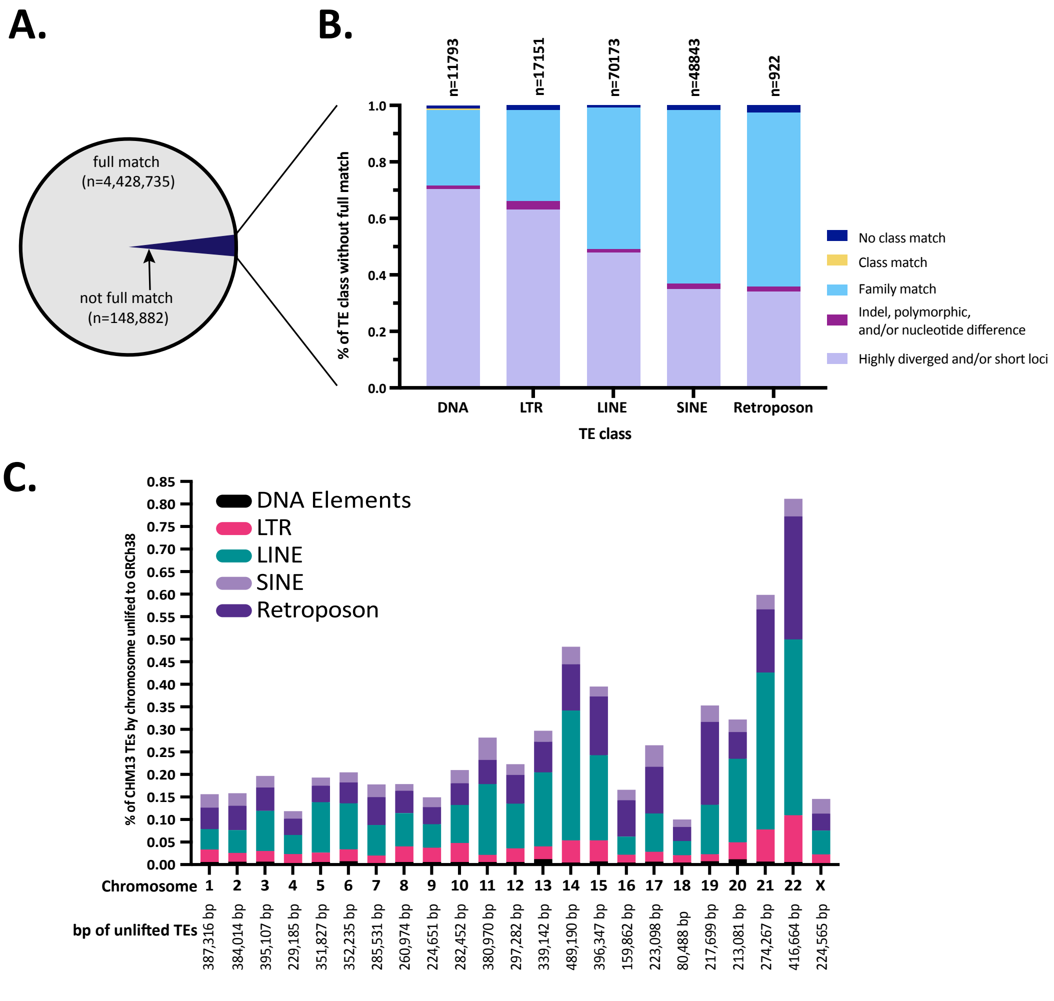


***Fig. S5. Lifted TE pair annotations are discordant between CHM13 and GRCh38. (A)*** *TEs in CHM13 with a full match with GRCh38 represent TEs with no change in annotation. The remaining 148,882 without a full match were further classified by discordance category.* ***(B)*** *"Not full match" classifications were further broken down into discordance categories as follows: 1) TEs lacking a class match (dark blue), 2) TEs with a class match but changed family (yellow), 3) TEs with a family match with a subfamily change (light blue), 4) TEs with nucleotide differences (purple) and highly diverged sequences/short loci (light purple), both of which are low confidence changes that may be the result of batch effects with RepeatMasker.* ***(C)*** *Of TE annotations in CHM13, percent unlifted to GRCh38, shown by chromosome (X axis) and repeat class normalized to bp of each chromosome in CHM13. TE family indicated by color in key inset; n= number of bp on each chromosome in CHM13 represented by unlifted TEs.*

| TE class | no discordance | Subfamily reclassification | | notable discordance | | | Total per TE class |
|---|---|---|---|---|---|---|---|
| | Full match | Highly diverged and/or short loci | Family match | Class match | No class match | Indel, polymorphic, and/or nucleotide differences | |
| **DNA** | 511833 | 8310 | 3153 | 57 | 141 | 132 | 523626 |
| **LTR** | 722493 | 10824 | 5540 | 0 | 277 | 510 | 739644 |
| **LINE** | 1482351 | 33631 | 35209 | 0 | 529 | 804 | 1552524 |
| **SINE** | 1706522 | 17126 | 29995 | 0 | 787 | 935 | 1755365 |
| **Retroposon** | 5536 | 314 | 568 | 0 | 23 | 17 | 6458 |
| **All classes** | 4428735 | 70205 | 74465 | 57 | 1757 | 2398 | 4577617 |
| **All discordances** | | 148882 | | | | | |
| **Combined discordance categories** | | 144670 | | 4212 | | | |

**Table S5.** *LiftOver counts for each lifted TE class broken down into classification level matches (Fig. S5). Among discordant loci, the vast majority (97.2%) are subfamily reclassifications. The remainder, notable discordant loci represent those with no family match, no class, match, or are polymorphic between the two genomes.*

| TEs only | Counts | % of total |
|---|---|---|
| **lifted from CHM13-v1 to GRCh38** | 4577617 | 99.62% |
| **unlifted from CHM13-v1 to GRCh38** | 17434 | 0.38% |
| **Total** | **4595051** | **100.00%** |

| TE Class | Syntenic | Non-syntenic | Unlifted total |
|---|---|---|---|
| **DNA element** | 803 | 170 | 973 |
| **LTR** | 1924 | 439 | 2363 |
| **LINE** | 3057 | 1068 | 4125 |
| **SINE** | 6288 | 2978 | 9266 |
| **Retroposon** | 612 | 95 | 707 |
| **Total** | **12684** | **4750** | **17434** |

***Table S6.*** *LiftOver statistics for TEs shown in Fig. 1C by total counts of TEs (top) and by unlifted TE class (bottom).*

**Copy Number Comparison across primates**

Copy number comparisons across primate genomes were generated with the most recent, available primate genomes for each species: *Pan trogolodytes* (accession: GCA_002880755.3) (*18*), *Gorilla gorilla* (accession: GCA_900006655.3)(*19*), *Pongo abelii* (accession: GCA_002880775.3) (*18*), *Hylobates moloch* (accession: GCA_009828535.2), *Macaca mulatta* (accession: GCA_008058575.1)(*20*), *Rhinopithecus roxellana* (accession: GCF_007565055.1)(*21*), *Callithrix jacchus* (accession: GCF_009663435)(*22*), and *Microcebus murinus* (accession: GCF_000165445.2) (*23*). Custom BLAST databases were generated from each genome and searched for individual instances of the corresponding repeat or composite element. Due to the varying quality and completeness of these genomes, and in order to avoid returning individual composite subunits, the search was done by requiring at least an 85% length match to the query repeat / composite monomer. Standard BLASTN parameters were used for query match divergence, except in the case of highly similar gap tandem array sequences, which were rerun with a 100% match requirement across the 85% length to assure correct counts. All results were quantified manually, and coordinates were checked within each set of results to ensure that only individual instances were counted.

#### Methylation Metaplots

Nanopore CpG methylation data for CHM13 and HG002 was processed according to the methods outlined in Gershman et al., 2021. Genomic coordinates were normalized by the repeat start and end for each repeat type and CpG methylation frequency was calculated by fraction of methylated reads to total coverage within bins in CHM13 or HG002 with the BSGenome Bioconductor package (https://bioconductor.org/packages/BSgenome) (*24*). Multiples of three bins were further smoothed with the "rollmean" function from the R package Zoo (https://cran.r-project.org/web/packages/zoo/ index.html).

#### Dot Plot Analyses

To generate pairwise sequence identity dot-plots we used the software package StainedGlass. The input for this program is sequence fragmented into windows (1Kbp) after which all possible pairwise alignments between the fragments are calculated using minimap2 (*25*). The color used in the dot-plot was then determined by the sequence identity of the alignment which was calculated as:

$$ ID = 100 \biggl( \frac{M}{M+X+I+D} \biggr) $$

where $ID$ was the percent sequence identity, $M$ the number of matches, $X$ the number of mismatches, $I$ the number of insertion events, and $D$ the number of deletion events. When there were multiple alignments between the same two sequence fragments all alignments other than the one with the most matches were filtered out regardless of their sequence identity. The resulting matrix of percent identity scores was then visualized using ggplot and geom_tile. All code and documentation are available at: https://mrvollger.github.io/StainedGlass/

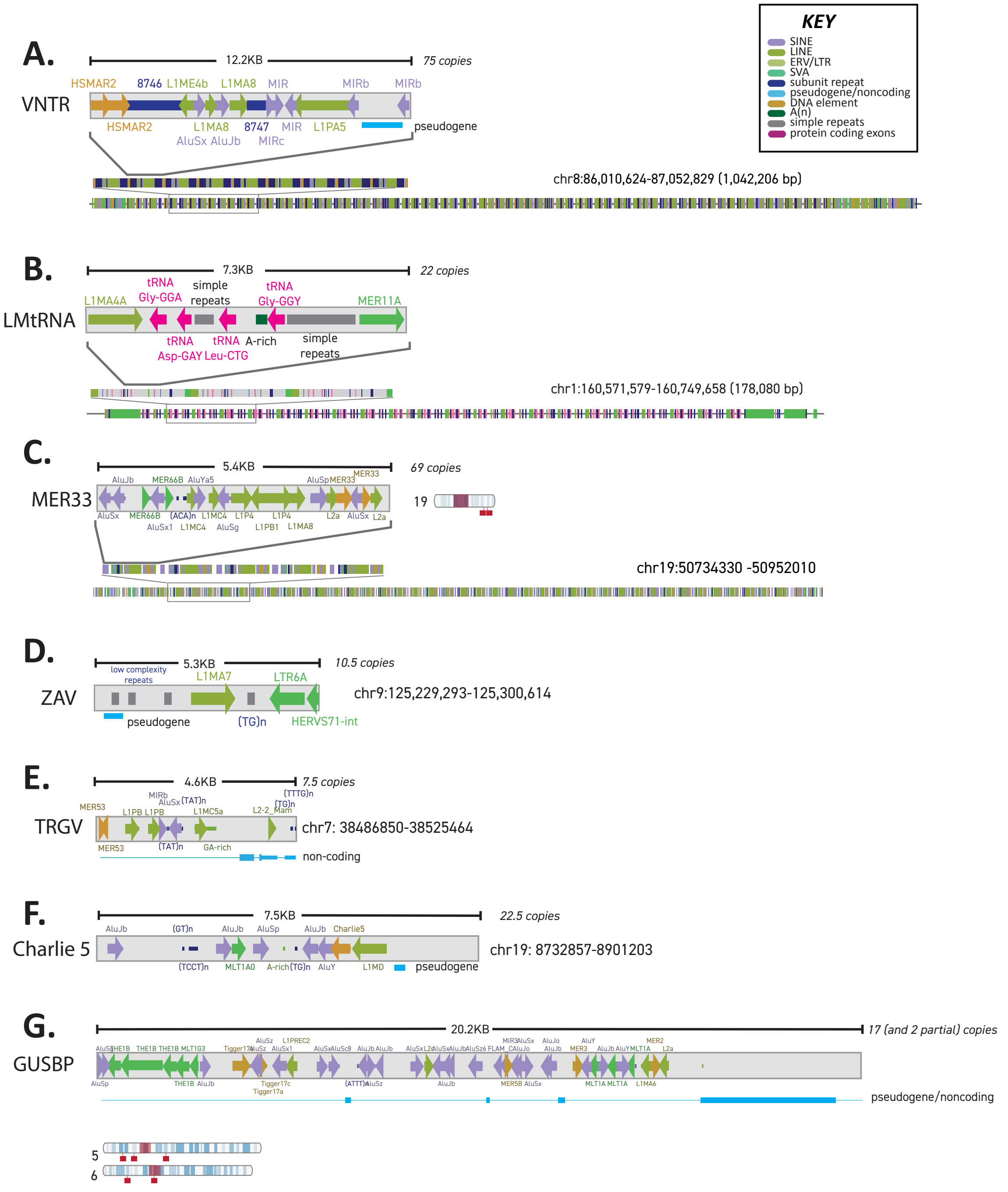


***Fig. S6. Annotations for composite elements in CHM13v1.0 lacking exonic material.*** *Repeat annotations for composite elements found without protein coding sequences:* ***(A)*** *VNTR,* ***(B)*** *LMtRNA,* ***(C)*** *MER33,* ***(D)*** *ZAV,* ***(E)*** *TRGV,* ***(F)*** *Charlie 5,* ***(G)*** *GUSP. Each subunit repeat type and orientation, including all TEs, simple and low complexity repeats, subunits and other features such as pseudogenes, color coded as per KEY (inset at top), is indicated. For each composite, the size of the composite core unit is indicated, as is the number of repeat units in the array and chromosomal location. The order of core units within the array are shown in two zoom images of Repeat Maskerv2 browser tracks for repeats found in large arrays* ***(A-C).*** *For composites found arrayed in more than one location* ***(C, G)****, chromosome ideogram indicates the locations of the composite arrays in CHM13. Centromere blocks (including centromere transition regions) are indicated in orange.*

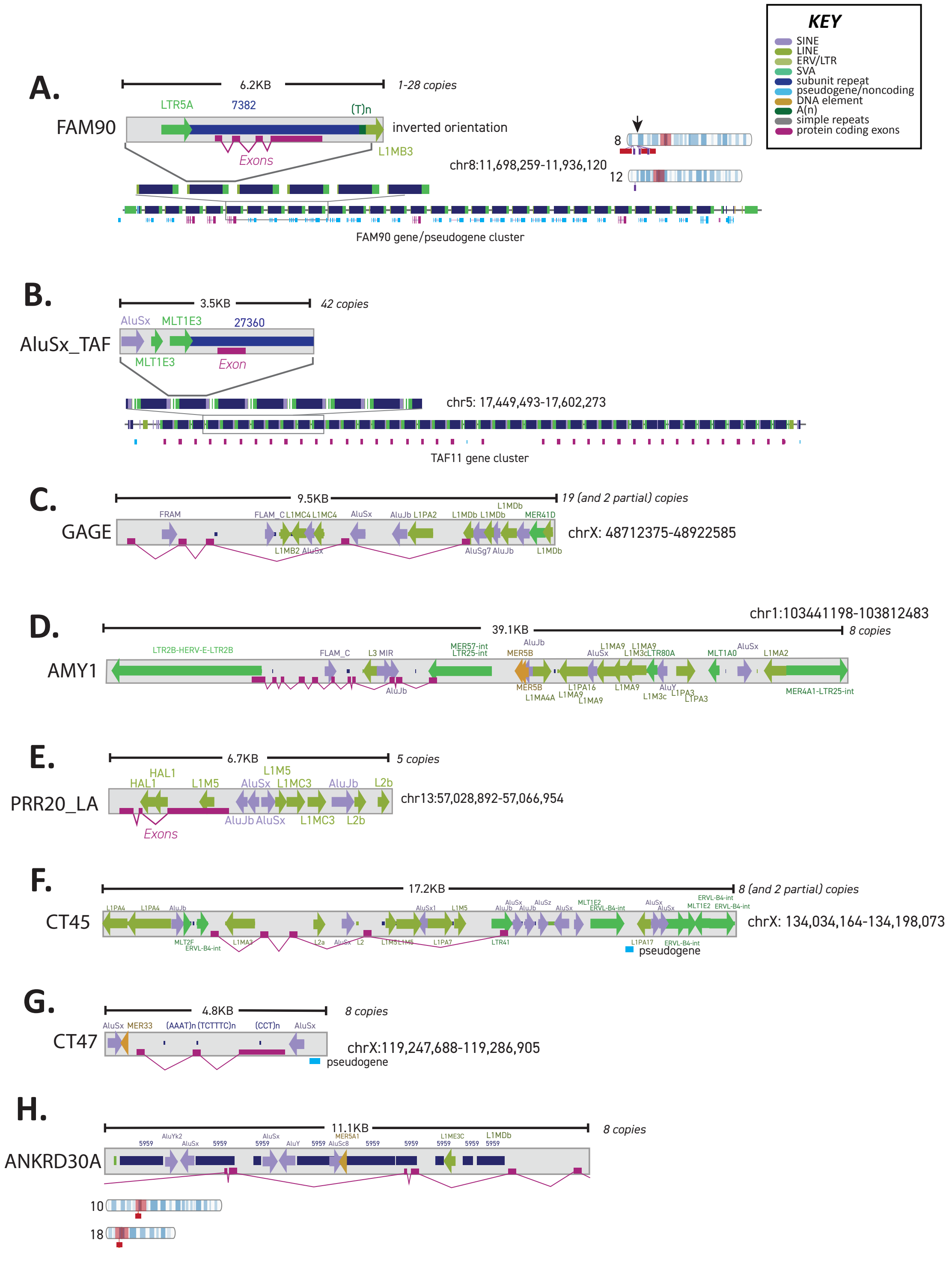


***Fig. S7. Updated annotations for composite elements containing gene predictions found in CHM13v1.0***

*Repeat annotations for composite elements found without protein coding sequences: **(A)** FAM90, **(B)** AluSx-TAF, **(C)** GAGE, **(D)** AMY, **(E)** PRR20_LA, **(F)** CT45, **(G)** CT47, **(H)** ANKRD30A. Each subunit repeat type and orientation, including all TEs, simple and low complexity repeats, subunits and other features such as exons, color coded as per KEY (inset at top), is indicated. For each composite, the size of the composite core unit is indicated, as is the number of repeat units in the array and chromosomal location. The order of core units within* *the array are shown in two zoom images of Repeat Maskerv2 browser tracks for repeats found in large arrays* ***(A-B).*** *For composites found arrayed in more than one location* ***(A, H)****, chromosome ideogram indicates the locations of the composite arrays in CHM13. Centromere blocks (including centromere transition regions) are indicated in orange.*

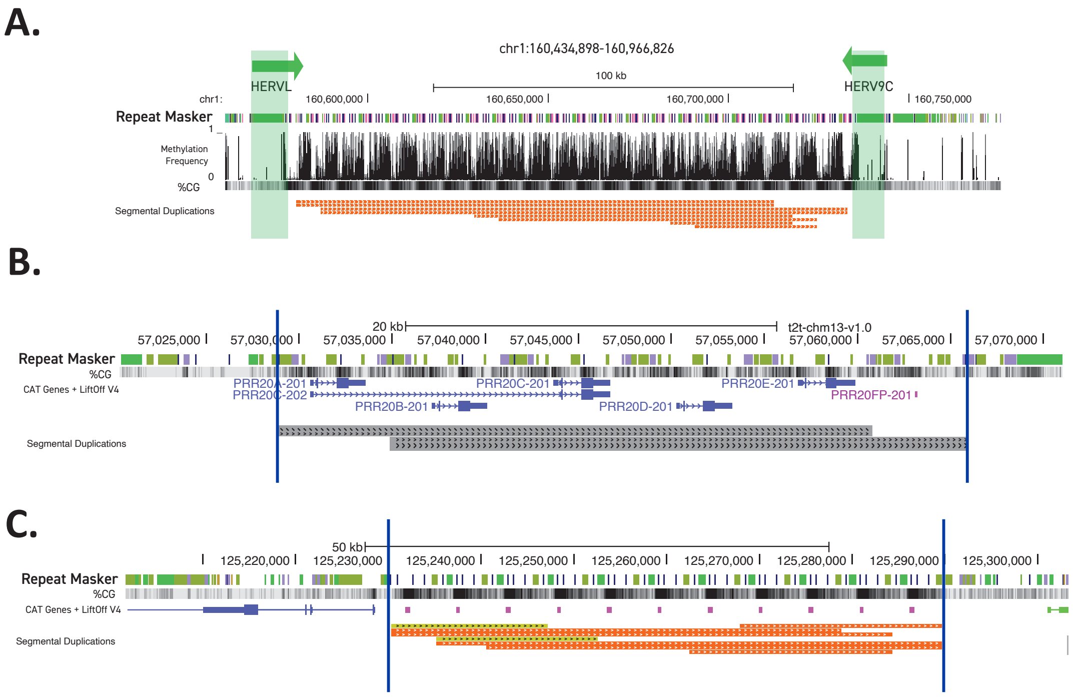


***Fig. S8. Discerning composite units from segmental duplications.***
***(A)*** *Browser track of the LMtRNA locus showing methylation frequency, %CpG, and segmental duplication annotations. HERVs flanking the array are shown in green, with transcript unit orientation indicated with an arrow.* ***(B)*** *and* ***(C)*** *Browser tracks of PRR-LA and ZAV composite arrays showing %CpG, gene annotations, and segmental duplication annotations. Array boundaries are indicated by a vertical line.*

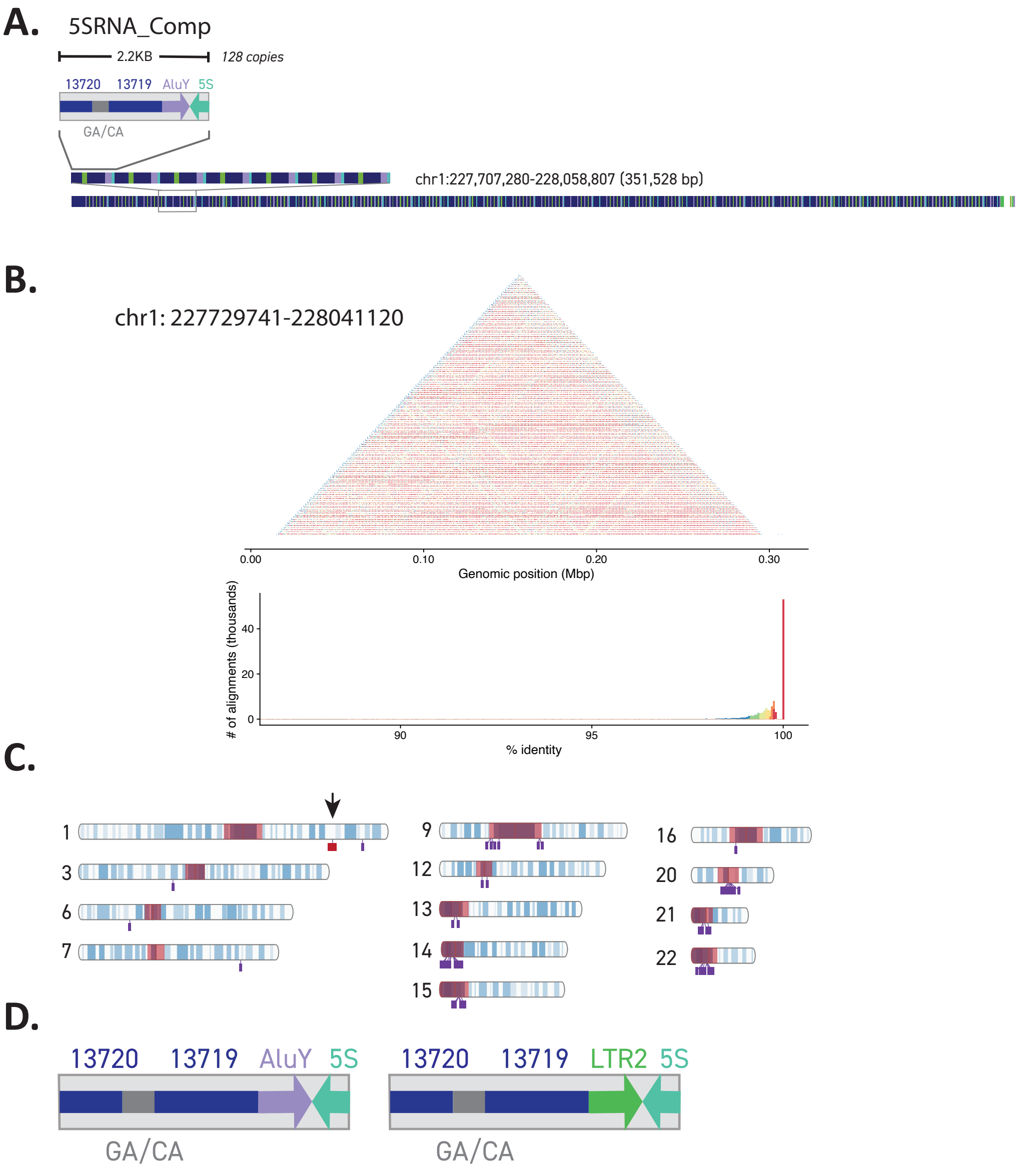


***Fig. S9. A composite containing part of the 5SRNA is found at multiple loci in CHM13 and arrayed at a single locus.***
***(A)*** *The 5SRNA composite contains an AluY insertion, three previously known repeats (GA-rich low complexity repeat, CA simple repeat, 5S) and two other newly annotated composite subunits (13719, 13720). The composite is found in an array on Chromosome 1. The order of core units within the array are shown in two zoom images of Repeat Maskerv2 browser tracks.* ***(B)*** *A self-alignment dot plot of 5SRNA composite subunits across the array. Histogram denotes the color scale and distribution of alignments for the plot showing high intra-array sequence similarity (with a peak ~100%). The array is visible in the dot plot as the brighter red triangle shape, in which connecting diagonals (the arms of the triangle) represent shared sequence identity A 5% size (bp) increase was added flanking the array, which is visible as the area with lower shared sequence identity (blue) on the left and right of the dot plot.* ***(C)*** *Ideogram of CHM13 indicates the 49 locations of the 5SRNA composites as singletons (purple) and the only array found in CHM13 (red). Arrow indicates the location of the array illustrated in* ***(A),*** *which is also the only location containing the AluY insertion. Centromere blocks (including centromere transition regions) are indicated in orange.* ***(D)*** *The 5SRNA composite is found as two different structures: with an AluY insertion (left) in array form and with a LTR2 insertion (right) at all*

*monomeric locations. Both contain three previously known repeats (GA-rich low complexity repeat, CA simple repeat, 5S) and two other newly annotated composite subunits (13719, 13720).*

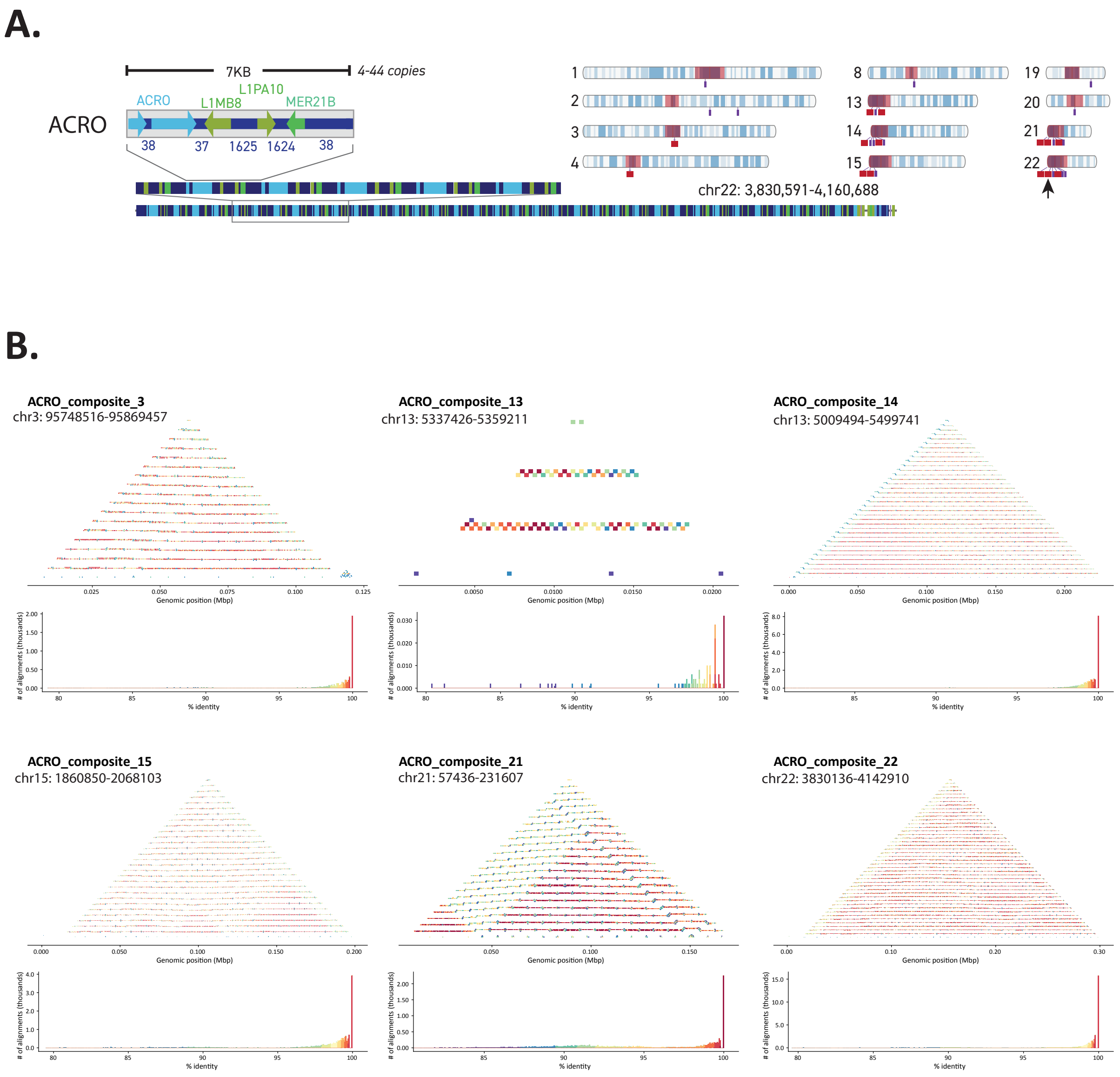


***Fig. S10. ACRO_Composites are found in arrays on multiple chromosomes in CHM13.*** ***(A)*** *Structure of the ~7Kbp ACRO composite includes four previously known repeats (ACRO, L1MB8, L1PA10, MER21B) and four newly annotated composite subunits (37, 38, 1624, 1625). Ideogram of CHM13 indicates the 30 locations of the ACRO composites as both singletons (purple) and arrays (red). Centromere blocks (including centromere transition regions) are indicated in orange. Arrow indicates the location of the array illustrated on the left.* ***(B)*** *Self-alignment dot plots for the largest ACRO composite arrays on each acrocentric chromosome (chr13, chr14, chr15, chr21, chr22) and one non-acrocentric chromosome (chr3) are shown displaying alignment similarity with histograms denoting the color scale and distribution of alignments for each independently colored plot. High sequence identity is seen within each array (>95%, excluding extra flanking region) and strong structural similarities between each array (with chr21 being a slight outlier, although the array still has the triangular structure typical*

*of a highly repetitive sequence). A 5% size (bp) increase was added flanking each array. Note that these plots differ in structure from Fig. S9 due to the size of the arrays.*

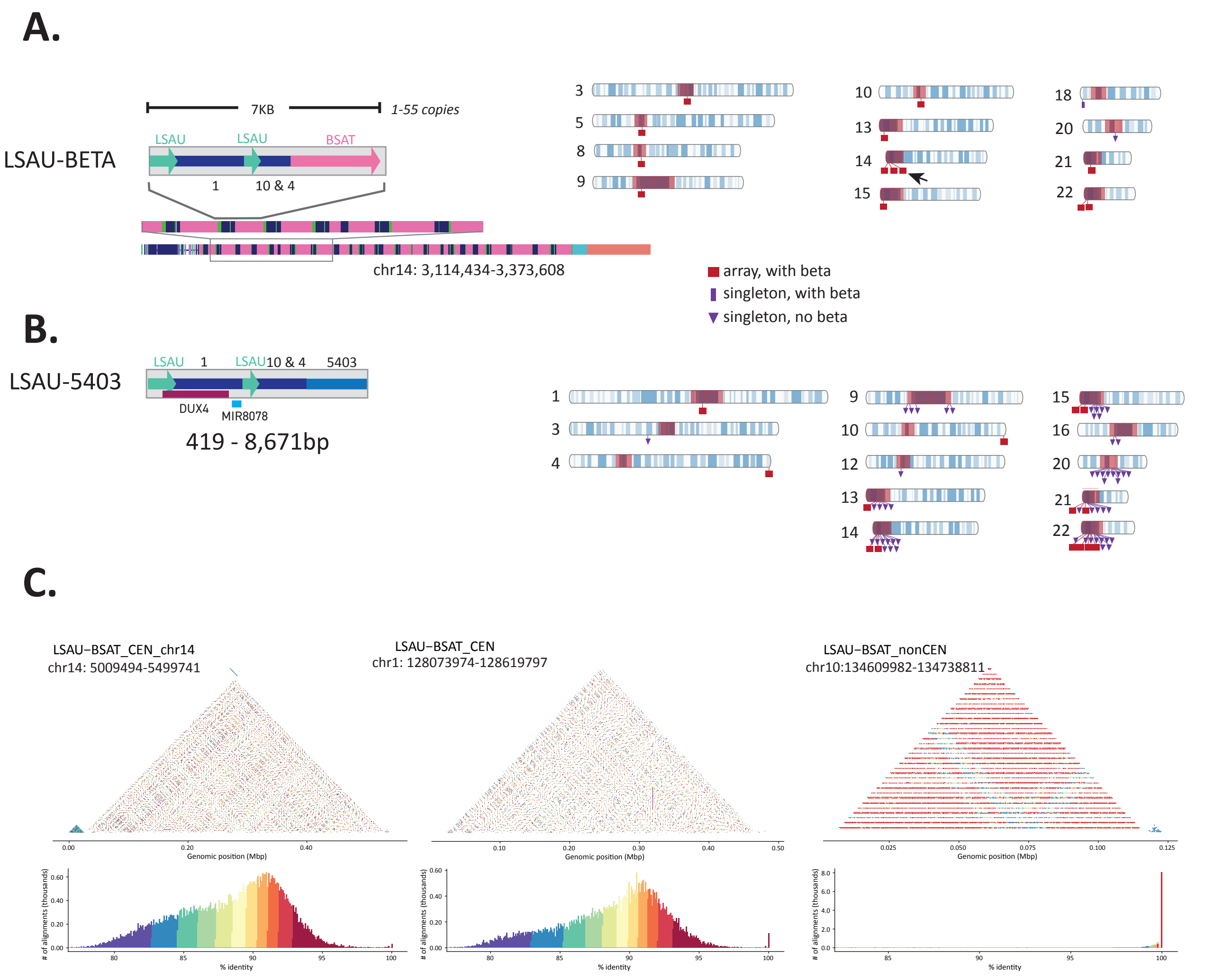


***Fig. S11. LSAU-BETA_Composites are found in two forms and arrayed on multiple chromosomes in CHM13.***
***(A)** The LSAU-BETA composite is found as two different structures: without the newly annotated 5403 composite subunit (Top; 15 loci) and with the repeat-5403 (Bottom; 57 loci). Both contain three other newly annotated composite subunits (1, 4, 10), as well as the LSAU satellite. Ideograms of CHM13 for each of the two different structures indicate the locations of the LSAU-BETA composites as singletons (purple) and arrays (red), as well as the presence (square) or absence (triangle) of the BETA satellite. BETA is not shown in the structure containing 5403 since it is not part of the composite itself, but rather found at one edge of each of the two arrays. These two arrays (chr4, chr10) are subtelomeric and associated with the DUX4 genes (4). Only one locus (chr8) lacks the LSAU satellite (star). Centromere blocks (including centromere transition regions) are indicated in orange. Arrows indicate the location of the arrays illustrated on the left. **(B)** Self-alignment dot plots for three example arrays (two centromeric and one non-centromeric) are shown displaying alignment similarity with histograms denoting the color scale and distribution of alignments for each independently colored plot. The spectral color scheme represents a scale of 0 (purple) to 100% (red) sequence similarity. The chr1 and chr10 loci both*

contain the 5403 subunit, while the chr14 locus does not. The two centromeric arrays have a lower intra-array sequence similarity (with a normal distribution suggesting a lack of non-random sequence similarity) compared to the non-centromeric array (with a single peak at ~100%). This suggests a much more complex structure in centromeric arrays, regardless of the presence of the additional 5403 subunit. The complexity of this array is also apparent in the Chromosome 10 non-centromeric array, given that the array pattern is broken approximately ⅔ of the way through (blue pattern). A 5% size (bp) increase was added flanking each array and each plot is colored independently.*

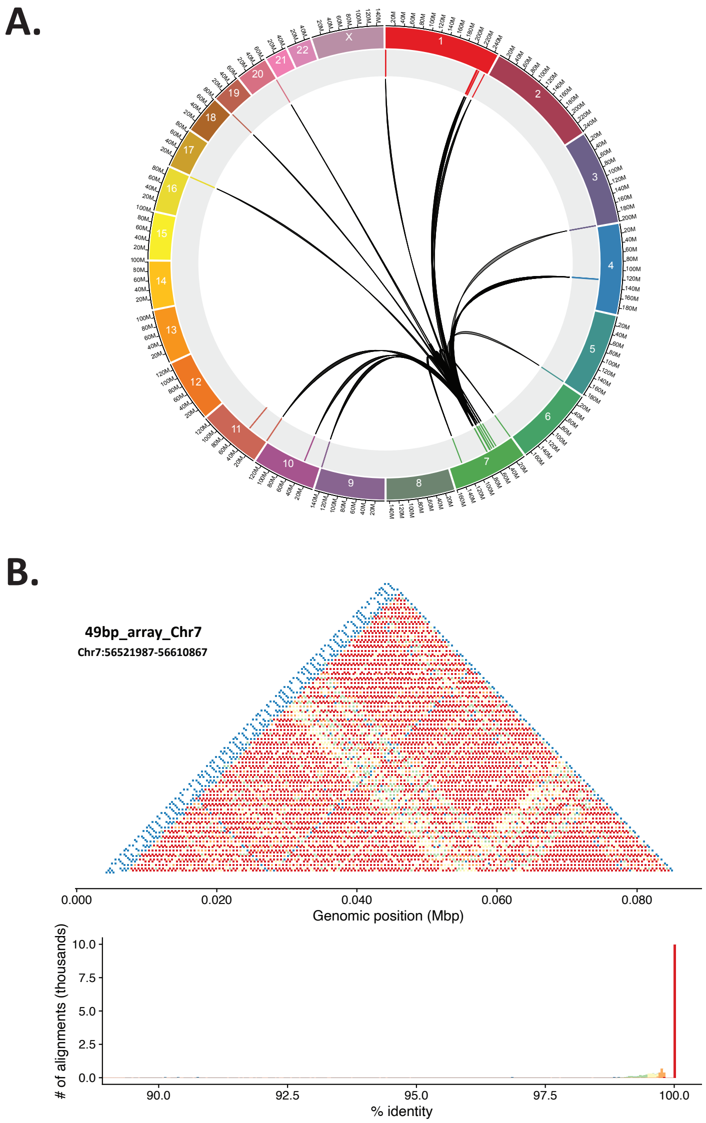


***Fig. S12. Segmental duplications spread TELO_Comp loci. (A)** Segmental duplication synteny map connecting TELO_Comp loci found in the centromere of Chromosome 7 with other TELO_Comp loci (Fig. 2B, D, Table S9) **(B)** A self-alignment dot plot of the 49bp array (ajax) on Chromosome 7 and histogram denoting the color scale and distribution of alignments for the plot showing high intra-array sequence similarity (with a peak ~100%). The array is visible in the dot plot as the brighter red triangle shape, in which connecting diagonals (the arms of the triangle) represent shared sequence identity. A 5% size (bp) increase was added flanking the*

*array, which is visible as the area with lower shared sequence identity (blue) on the left and right of the dot plot.*

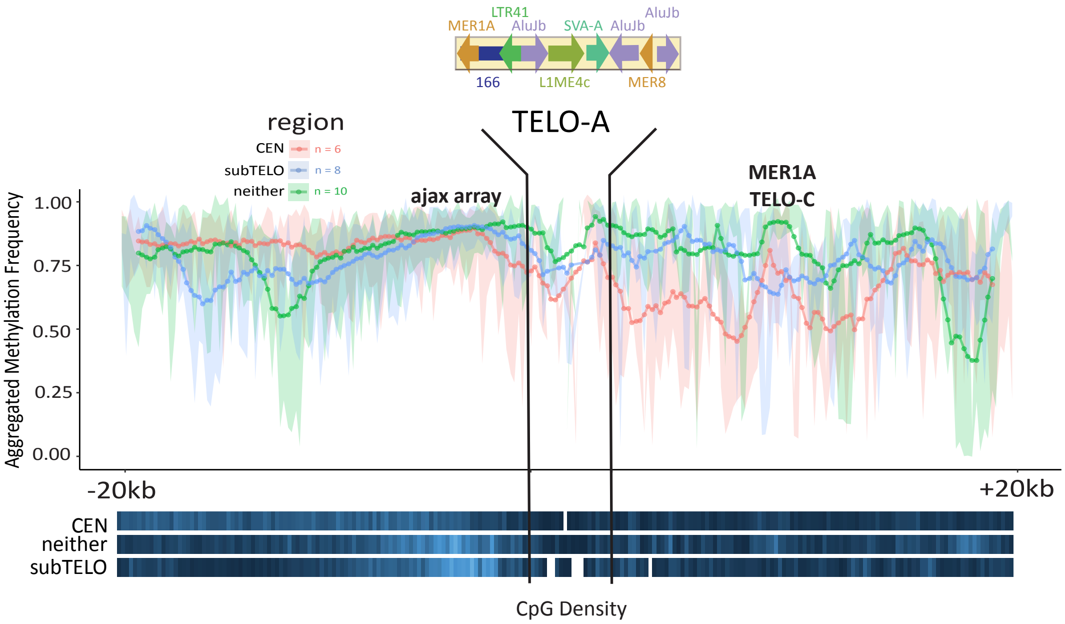


***Fig. S13. Methylation metaplot for HG002 reveals a unique epigenetic signature specific to TELO-Comp elements.***
*Metaplot of aggregated methylation frequency (average methylation of each bin across the region, 100 bins total) centered on the TELO-A subunit (top), ±20Kbp, grouped by chromosomal location (orange – centromeric, blue – subtelomeric, green – interstitial). CpG density for each group is indicated at the bottom (white - no CpG, dark blue - low CpG, bright blue - high CpG). The location of the ajax repeat array and the MER1A element within the TELO-C subunit are indicated (top).*

### 3. Phylogenetic Analyses

#### TELO_Comp Phylogenetic Analyses

CHM13 annotations for TELO_Comp TELO-A subunit were extracted from the genome as fasta sequences via bedtools (*8*) (Table S9). Sequences were aligned with MUSCLE (*26*). The evolutionary history was inferred by using the Maximum Likelihood method and General Time Reversible model (*27*). The tree with the highest log likelihood (-8364.23) is shown in Fig 2D. The percentage of trees in which the associated taxa clustered together is shown next to the branches. Initial tree(s) for the heuristic search were obtained automatically by applying the Maximum Parsimony method. A discrete Gamma distribution was used to model evolutionary rate differences among sites (5 categories (+G, parameter = 0.2836)). The tree is drawn to scale, with branch lengths measured in the number of substitutions per site. This analysis involved 24 nucleotide sequences with a total of 2845 positions in the final dataset. Evolutionary analyses were conducted in MEGA X (*28*, *29*).

### WaluSat and WaluSx Phylogenetic Analyses

#### *WaluSat+AluSx:*

*WaluSat:* The evolutionary history was inferred by using the Maximum Likelihood method and General Time Reversible model (*27*). The tree with the highest log likelihood (-2227.42) is shown in Fig. 3E. Initial tree(s) for the heuristic search were obtained automatically by applying the Maximum Parsimony method. A discrete Gamma distribution was used to model evolutionary rate differences among sites (5 categories (+G, parameter = 2.0168)). The tree is drawn to scale, with branch lengths measured in the number of substitutions per site. This analysis involved 1057 nucleotide sequences. There were a total of 73 positions in the final dataset. Evolutionary analyses were conducted in MEGA X (*28*).

*AluSx (with WaluSat):* The evolutionary history of AluSx-like elements was inferred by using the Maximum Likelihood method and T93 model. The tree with the highest log likelihood (-2155.25) is shown. The percentage of trees in which the associated taxa clustered together is shown next to the branches. Initial trees for the heuristic search were obtained automatically by applying Neighbor-Join and BioNJ algorithms to a matrix of pairwise distances estimated using the Maximum Composite Likelihood (MCL) approach, and then selecting the topology with superior log likelihood value. A discrete Gamma distribution was used to model evolutionary rate differences among sites (+G, parameter = 0.8559). This analysis involved 71 nucleotide sequences. (partial deletion option). There were a total of 289 positions in the final dataset. Evolutionary analyses were conducted in MEGA X (*28*)

#### *SST1 Phylogenetic Analyses*

CHM13 annotations were combined with known repeat locations and extracted from the genome as sequences via bedtools (*8*). The resulting elements were annotated with chromosome, coordinates, full length, intersection of centromere, telomere or interstitial chromosomal locations, and average methylation of the element. Sequences were aligned with MAFFT (*10*). The evolutionary history was inferred by using the RAxML method (*30*) and the GTR+G (general time reversible model with a gamma distribution of rate variation among sites) model (*27*) as matched by jModelTest (*31*). The consensus tree shown in Fig. 4A was generated from the resulting 100 bootstrap replicates.

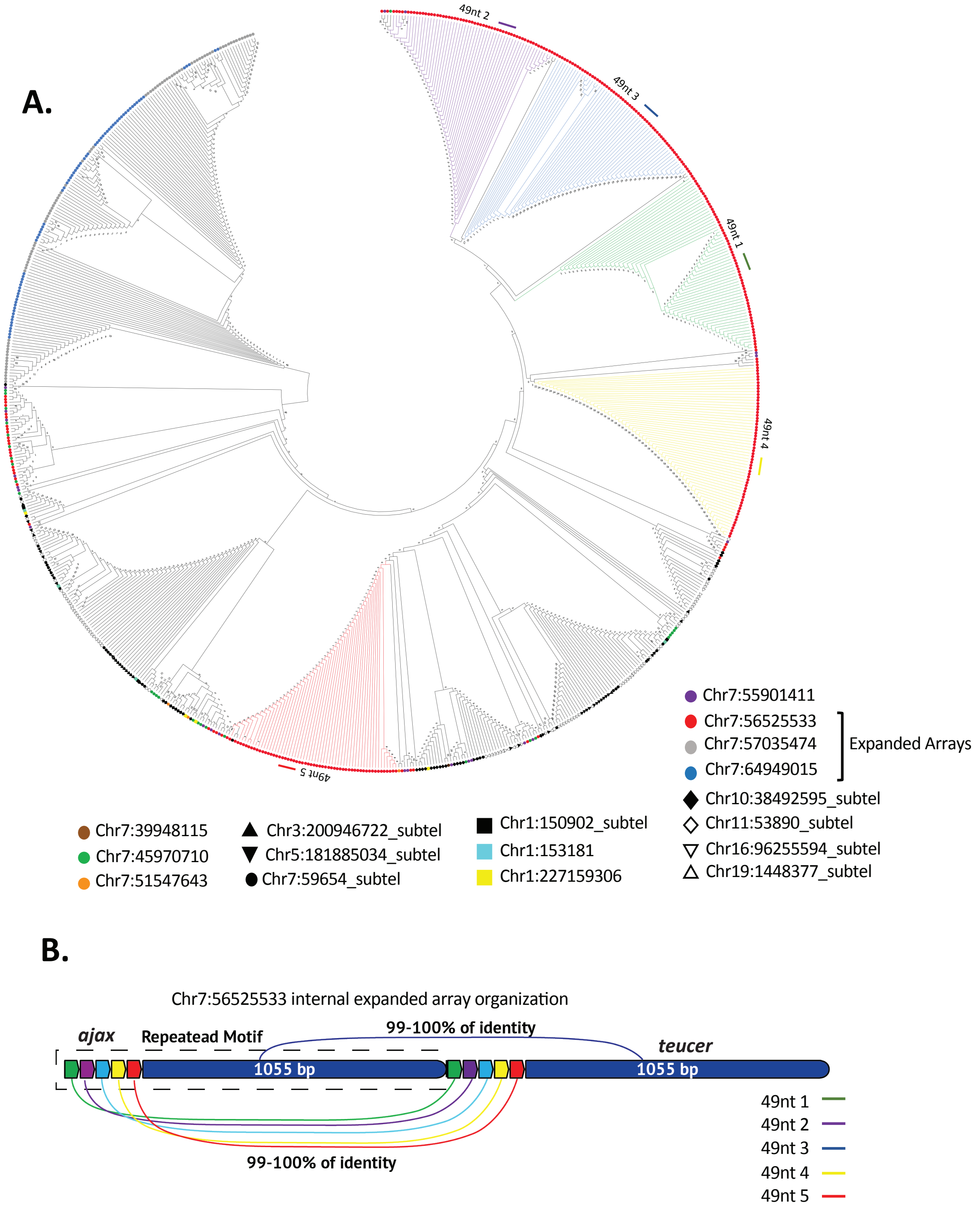


***Figure S14. Phylogenetic analyses of ajax repeats reveals different patterns of evolution influence subtelomeric vs pericentromeric arrays. (A)*** *A Neighbor Joining unrooted phylogenetic tree using 763 full-length repeats (Table S11) reveals ajax repeats in subtelomeric regions (black and open shapes) do not cluster together in array-specific or chromosome-specific subtrees. In contrast the ajax pericentromeric repeats from each of the three expanded*

*loci tend to cluster in array-specific clades, suggesting that these arrays evolve under concerted evolution. Chr7:56525533 ajax monomers are indicated by colored branches green, purple, light blue, yellow, and red for monomers 1 to 5 (see* ***(B)****), respectively, suggesting a higher order repeat, or super-repeat, structure across the array.* ***(B)*** *Schematic representation of Chr7:56525533 organization of ajax and the associated teucer repeat, which together comprise a composite element that evolves as a single unit (Fig. 2F). The 49nt monomeric ajax repeats of the Chr7:56525533 locus are indicated by colored arrowheads in green, purple, light blue, yellow, and red for monomers 1 to 5, respectively, whereas teucer is represented in dark blue.*

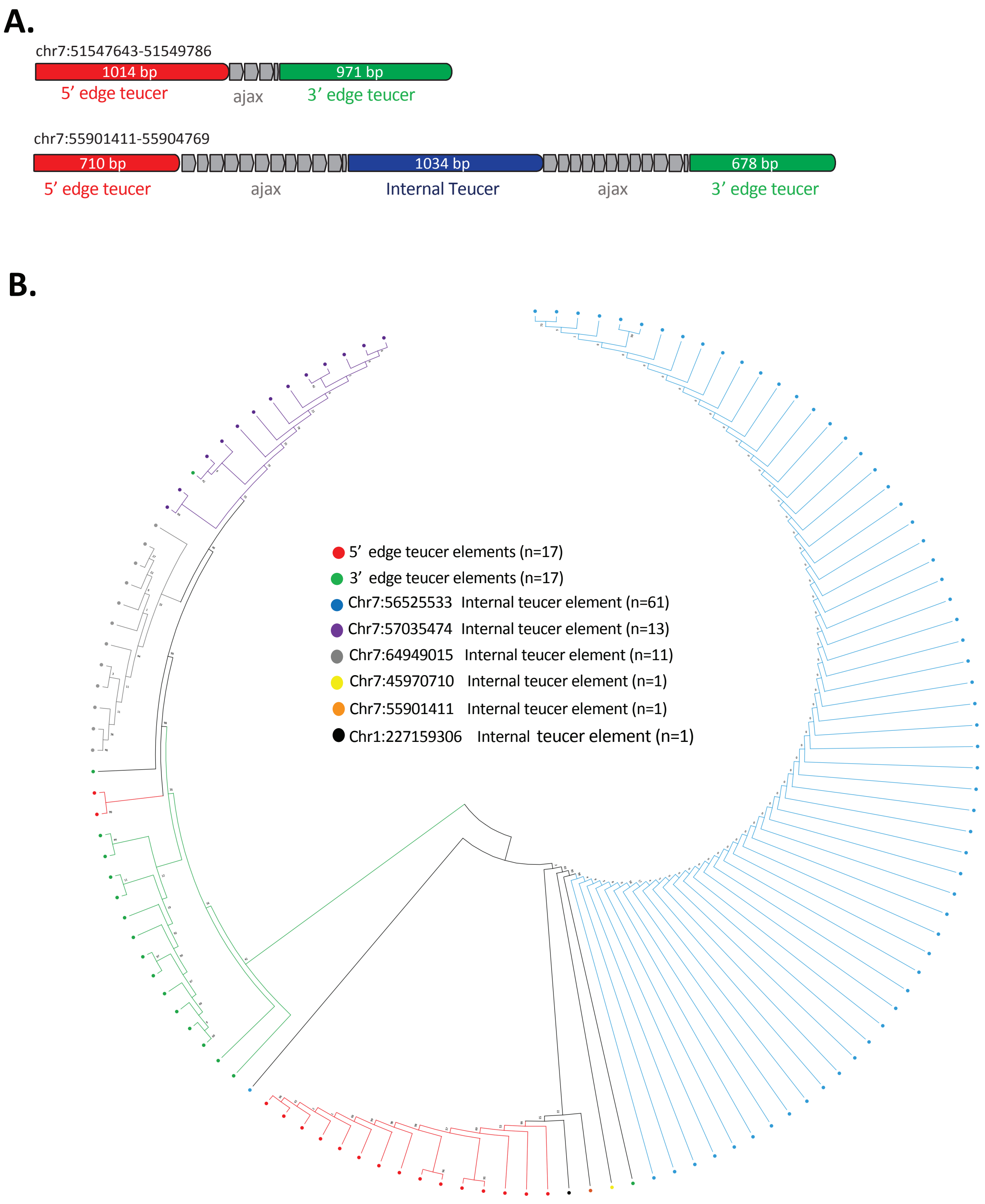


***Figure S15. Sequence relatedness between teucer arrays suggests location specific patterns of evolution and expansion.*** ***(A)*** *Schematic representation of ajax+teucer composite organization indicating 5’ edge teucer elements (red), ajax repeats (gray), internal teucer elements (dark blue), and 3’ edge teucer elements (green) described at two loci as indicated. Three separate consensi have been derived for the Teucer element based on the internal*

*sequence and the edge sequences. The different areas in which the element occurs appear to be under different evolutionary pressure and change at different rates.* ***(B)*** *A maximum likelihood phylogenetic tree using 122 teucer elements (Table S12) reveals that teucer derived from either the 5' edge (red) or 3' edge (green) form unique clusters, suggesting independent evolution by array position. Internal teucer elements from expanded arrays show array-specific clusters that correspond with the ajax phylogenetic analyses (Fig. S14). Chr7:56525533 internal teucer elements (light blue) show higher similarities with 5' end teucer elements, suggesting the 5' end contributed to the array expansion. The Chr7: 57035474 and Chr7: 64949015 arrays (purple and gray, respectively) cluster with 3' end teucer elements, suggesting the 3' end contributed to the array expansion. The relatedness of teucer elements from the three expanded arrays to the subtelomeric teucer elements suggest that independent events are responsible for the origin of Chr7:56525533 and Chr7: 57035474/Chr7: 64949015 arrays. All teucer elements were positioned in the same orientation and form the same junction with the ajax arrays.*

### 4. Transduction Analyses

#### Discovery of 3' and 5' transductions in CHM13

DNA transduction events for both chm13 and hg38 genome assemblies were analyzed using the modified TSDfinder tool (*32*), available at https://github.com/IOB-Muenster/TSDfinder. In order to identify 3' transductions mediated by Alu and L1 elements, we ran the TSDfinder with the default parameters (FIVE_PR_FLANK = 100, THREE_PR_FLANK = 3000) and the full-length SINE/Alu and LINE/L1 families present in the RepeatMaskerv2 output developed in this study. To identify SVA-driven 5' and 3' transductions, TSDfinder was run using full-length SVA elements with the following parameters: FIVE_PR_FLANK = 3000, THREE_PR_FLANK = 3000. Since TSDfinder originally was designed to find only 3' transductions, we used a one line command:

```
awk -F "\t" '{if ($1 ~ /\(/ ) print "-" "\t" length($6) "\t" $0 ; else print "+" "\t" length($6) "\t" $0 }'
PRE_ins_TEMP_"${chr_name}" | sed 's/(\|)//g' | sed \'s/ .. /\t/g' | awk -F "\t" '{if ($1 == "-" &&
($16-$2)-$4 >= 30) print $0 "\t" "5prTS"; else if  ($1 == "+" && $3-($15+$2) >= 30) print $0 "\t"
"5prTS"; else print $0}'
```

to identify 5' transductions based on the assumption there must be a 30-nucleotide distance between the 5' TSD and the start of the SVA. Those events labeled as “3prTS” (3prTS stands for 3' transduction) were subtracted and subjected to the validation process as described below.

#### Validating and categorizing the putative transductions

To validate and categorize each putative transduction discovered by TSDfinder, we defined a set of certain thresholds and five different confidence levels as follows (summarized in Fig. S16).

***Level 0:*** The lowest confidence level contains most of the transductions discovered by running TSDfinder. In this step, only those transductions whose both TSD and a poly(A) tail that consisted of homopolymer stretches of A's were removed as they likely are artifacts. This level consisted of 108,339 events.

***Level 1:*** In this step, the progenitor of each transduction event is identified. Initially, we extracted 3Kbp downstream from all family members of SINE/Alu, LINE/L1, Retroposon/SVA elements and made a “blastable" database using the following command from BLAST suite (v2.11.0+) (*33*):

*makeblastdb -in input.fa -dbtype nucl -parse_seqids*

In the case of SVA 5' transductions, 3Kbp upstream from the 5' end of all Retroposon/SVA elements were extracted to create a cognate database. Subsequently, the sequences of putative transductions were collected and masked with RepeatMasker (v4.1.0) (*34*) (*-q -species human -xsmall*), and aligned to their related databases using BLASTN (v2.11.0+) (*33*) (*-evalue 0.05 -max_target_seqs 5 -perc_identity 90*). The results of each BLAST search were analyzed to find the progenitor of each transduction event. A progenitor was considered to be the source of a transduced sequence only if all of the following criteria were met: 1) the identity between two sequences (i.e., a query and a subject) was equal or greater than 90%, 2) hit and subject had the same orientation, 3) at least 30% of the length of putative transduction was included in the alignment, 4) the start coordinates for a pair of corresponding query and subject were within 20 nucleotides of each other. Transductions whose progenitors shared the same 3Kbp flanking sequence were not included in the final list. The transductions that met all criteria were classified as level 1 confidence DNA transduction events. The first tier of filtering reduced the transduction events identified in CHM13 to 5,579 events.

***Level 2:*** The sequences of TSDs of transductions within the level 1 group were queried. In order to mitigate false positives and any imprecise calculation of transduction length, those transductions whose TSDs consisted of only homopolymer stretches or As or Ts were removed from the dataset. Therefore, members of this confidence level include transductions whose progenitors were found (Level 1) and their TSD sequence content is heterogeneous. The second tier of filtering reduced the transduction events identified in CHM13 to 5,284 events.

***Level 3:*** In this step, pairs of progenitor-offspring from Level 2 that were a part of segmental duplication events were screened and eliminated, based on the rationale that the relation of a progenitor and its corresponding offspring cannot be clearly attributed to a transduction event if the sequence identity extends well beyond the TSD (*32*). For 3' transduction events, 3Kbp downstream from each transduced segment in the progenitors was extracted and used to build a BLASTable database using the following command: *makeblastdb -in input.fa -dbtype nucl -parse_seqids*. Likewise, 3Kbp downstream of each transduced segment in the offspring was extracted and used as queries for BLAST searches. In the case of 5' transductions, 3Kbp upstream from the 5' ends were collected. Finally, we performed a BLAST analysis to find those progenitors-offspring pairs whose alignments extended past the TSD using the following command:

*blastn -query <input.fa> -db <database> -task megablast -evalue 0.01 -out <output_name> -outfmt '6 qseqid qlen sseqid slen length pident nident mismatch gaps qstart qend sstart send sstrand qcovs qcovhsp qcovus evalue score' -max_target_seqs 5 -soft_masking false -dust no -perc_identity 90*

We considered a pair of progenitor-offspring as part of a duplication event if all of the following thresholds were met: 1) at least 98% of the query length was aligned to the subject, 2) the number of gaps in the alignment was less than 20 nucleotides, and 3) the alignment start positions were within 20 nucleotides of each other. After marking and removing such duplication events, the remaining transductions were classified as Level 3. The third tier of filtering reduced the transduction events identified in CHM13 to 3,271 events.

***Level 4:*** Finally, we checked whether or not the remaining, Level 3 transduced segments carried another non-LTR (complete or partial) in their sequences. In many cases, the entire transduced DNA or its terminal fraction overlapped with another non-LTR element, rendering it difficult to confidently attribute a poly-A tail followed immediately by 3' TSD (one of the transduction signatures) to a real transduction event because they may occur coincidentally with the inserted sequence. Consequently, we defined two sub-classes for the highest confidence transduction calls, Level 4.a and 4.b. A transduction event was classified as Level 4.a if it only had another non-LTR in its middle section and was flanked by unique DNA sequence. We filtered out those transductions whose entire sequence or its terminal fraction overlapped with another non-LTR element. Similarly, if a transduced sequence was depleted of a non-LTR element, it was classified as Level 4.b. These filters were applied by intersecting the coordinates of transductions with non-LTR annotation obtained from RepeatMaskerv2 output. For this, we used bedtools intersect command (*8*) as follows:

For 4.a level:

```
bedtools intersect -wao -a <transduced_segments_LEVEL3.bed> -b <non-LTRs.chm13.bed> -f 1.0 -s | awk '$NF == 0 {print $0}' | cut -f1-6 > TEMP.bed
```

```
awk '{if ($6 == "-") print $1 "\t" $2 "\t" $2+50 "\t" $4 "\t" $5 "\t" $6; else print $1 "\t" $3-50 "\t" $3 "\t" $4 "\t" $5 "\t" $6}’ TEMP.bed > TEMP_50bp.bed
```

```
bedtools intersect -wao -a TEMP_50bp.bed -b ../<non-LTRs.chm13.bed> -s | awk '$NF == 0 {print $0}' | cut -f1-6> transductions.4a.bed
```

For 4.b level:

```
bedtools intersect -wao -a <transduced_segments_LEVEL3.bed> -b <non-LTRs.chm13.bed> -s | awk '$NF == 0 {print $0}' | cut -f1-6 > transductions.4b.bed
```

The fourth tier of filtering reduced the transduction events identified in CHM13 to 294 events in category 4.a. And 406 events in category 4.b.

In total, 356 Alus were progenitors for transduced DNA, among which *Alu*Sz with the size of 294 bp located on Chromosome 13 (coordinates: 10150959-10151253) seemed to be the most prolific with 4 offsprings, while the remaining progenitors each generated between 1 and 3 offsprings. In the case of L1, 175 elements were verified as the sources of L1 transductions among which L1PA2 with the size of 6015 bp (chr11:43501531-43507546) was the most productive LINE with 4 offsprings. As for SVA 3’ transductions, we found 47 elements as progenitors among which one element, SVA_F, had 3 offsprings and was the hotspot for producing SVA 3’ transductions (chr10:100720613-100723987). We find that 71 SVAs appeared to be sources of 5’ transductions. Five SVA loci, each with two offsprings, were hotspots for transducing genetic material across the genome as follows; chrX:41017260-41018573 (SVA_E), chr5:138057944-138060261 (SVA_D), chr5:127247501-127248869 (SVA_F), chr19:61243812-61244877 (SVA_E) and chr17:17858632 -17860263 (SVA_D). Interestingly, there were two SVA_F progenitors that had produced both 3’ and 5’ transductions; chr2:55647319-55648994 and chr6:107029321-107031165. The size of L1 progenitors ranged from 33 bp to 6163 bp with a median of 450 bp, suggesting that truncated L1 elements are also capable of transducing sequences and can produce as many offsprings as full-length elements, in agreement with (*35*). This pattern is also observed among Alu and SVA progenitors.

**Functional annotation of transduction events**

To investigate whether Level 4 transductions carry protein-coding sequences, we conducted a BLASTX (*33*) analysis. First, we downloaded the human proteome (https://ftp.ncbi.nlm.nih.gov/refseq/H_sapiens/mRNA_Prot/human.1.protein.faa.gz, on 15.05.2021) and created a BLAST database using the following command:

```
makeblastdb -in human_proteome.fa -dbtype prot -parse_seqids.
```

The sequences of Level 4 transductions were aligned to the human proteome database using blastx:

*blastx -query <input.fa> -db human_proteome.fa —task blastx-fast -evalue 0.00001 -outfmt '6 qseqid qlen sseqid slen length pident nident mismatch gaps qstart qend sstart send qseq sseq sstrand qcovs qcovhsp qcovus evalue score' -max_target_seqs 5 -soft_masking true*

Aligned sequences were further annotated in the Level 4 dataset of transduction events by location in CHM13.

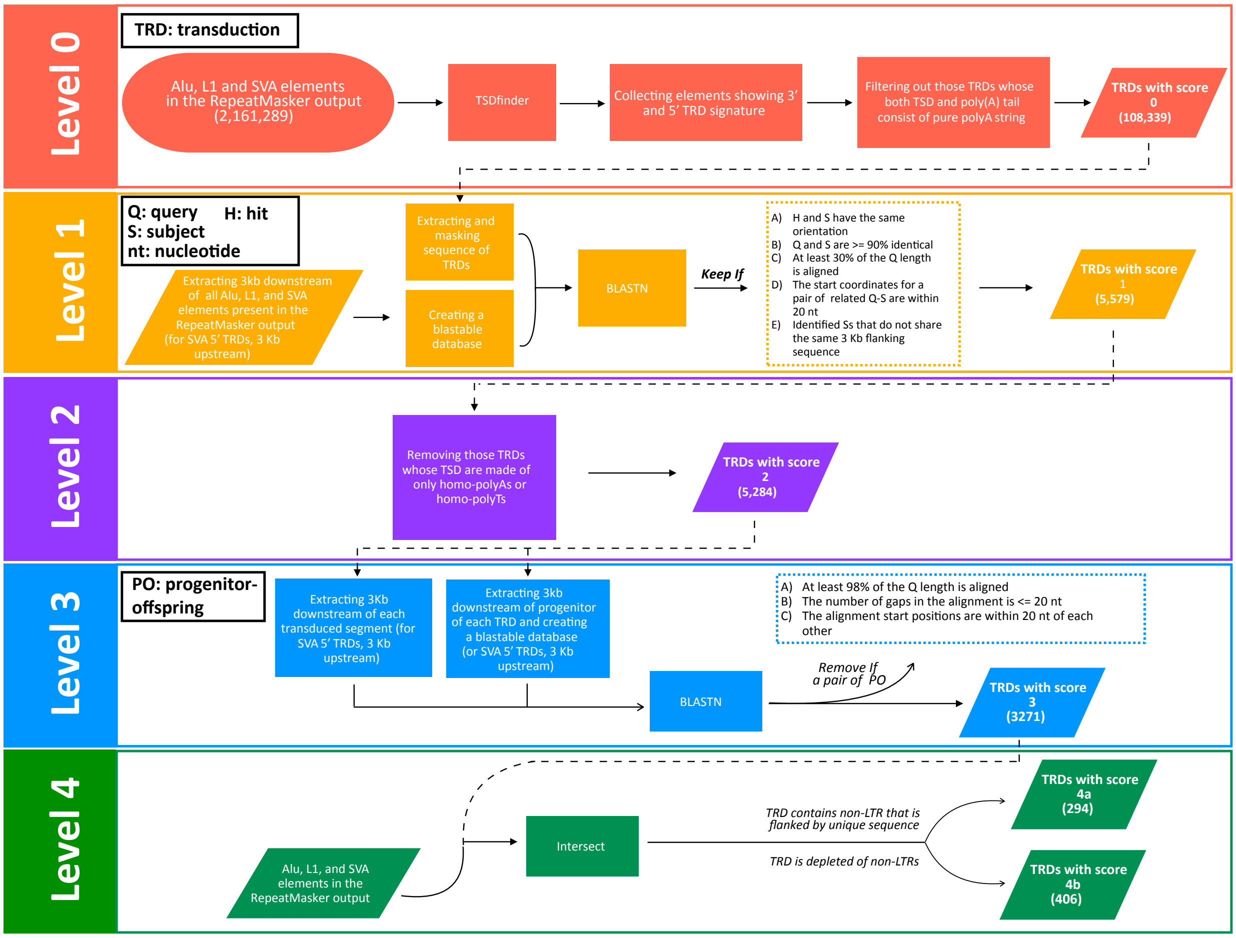


***Fig. S16.*** *Transduction analysis of CHM13 implemented four filtering steps to annotate high confidence events (Levels 1-4).*

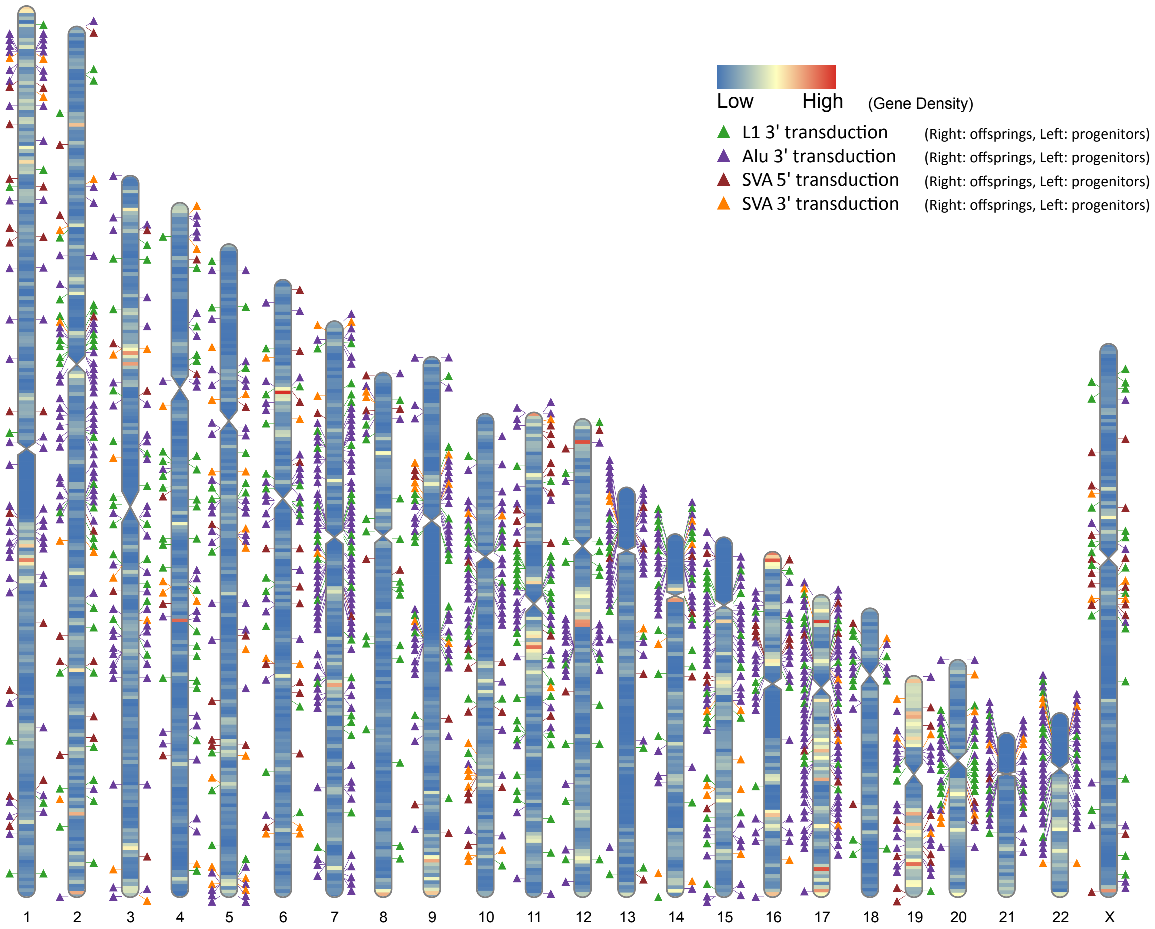


***Figure S17. Transduction events are found genome wide in CHM13.*** *Ideogram of CHM13 showing all transduction progenitors (left) and offspring (right) with respect to TE family (colored triangles) and gene density per 1Mbp bins.*

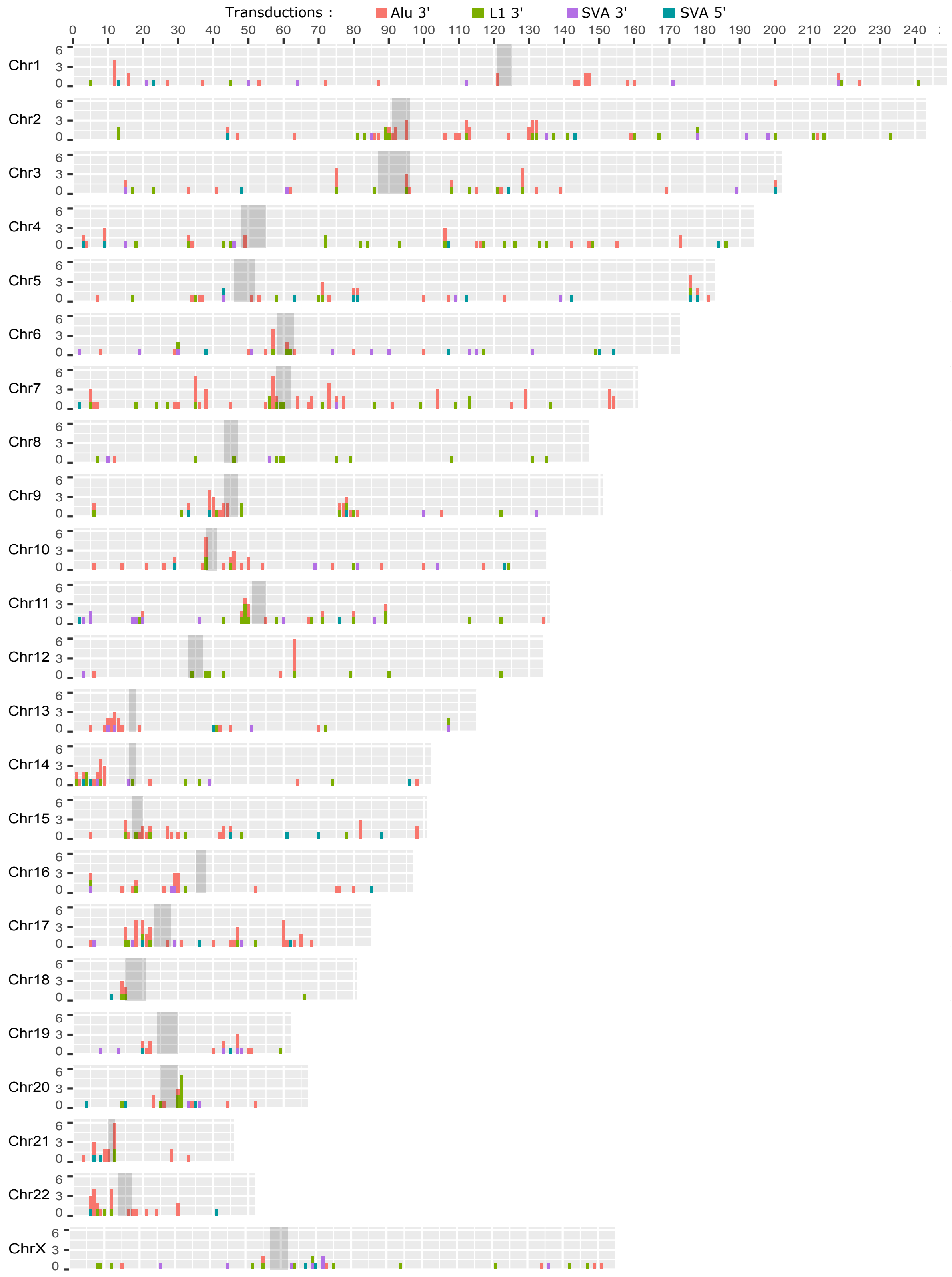


***Fig. S18.*** *Frequency of transduction events (offspring only) across each chromosome in CHM13v1.0. Element is indicated by color as per key at top.*

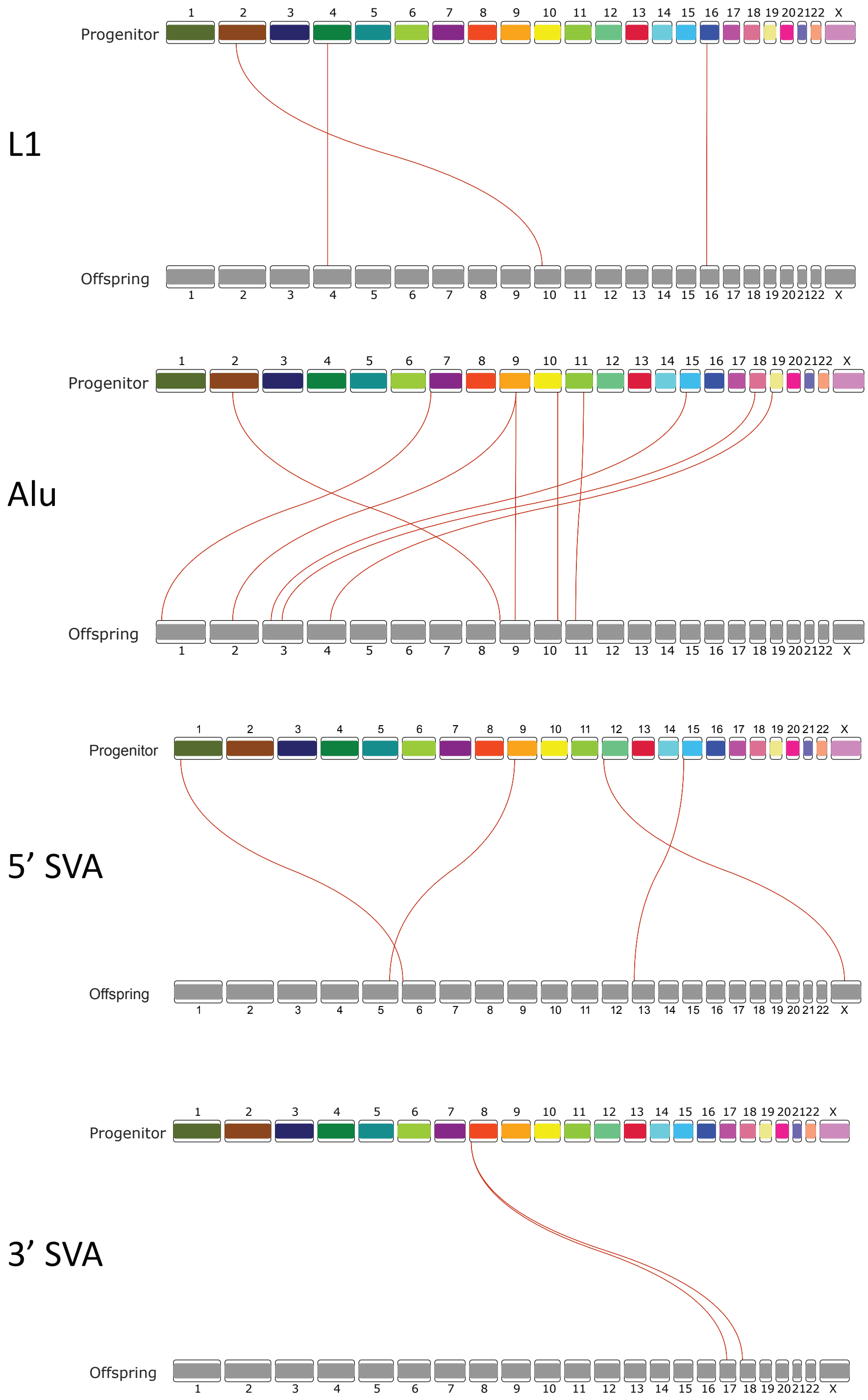


***Fig. S19.*** *Linkage graphs showing transduction progenitors and offspring across CHM13 for each repeat class (LINE, SINE, 5’ and 3’ SVA transduction events) as indicated. Red connections correspond to transduced sequences with >90% identity to protein-coding sequences (indicated with red arrowheads in Fig. 3A).*

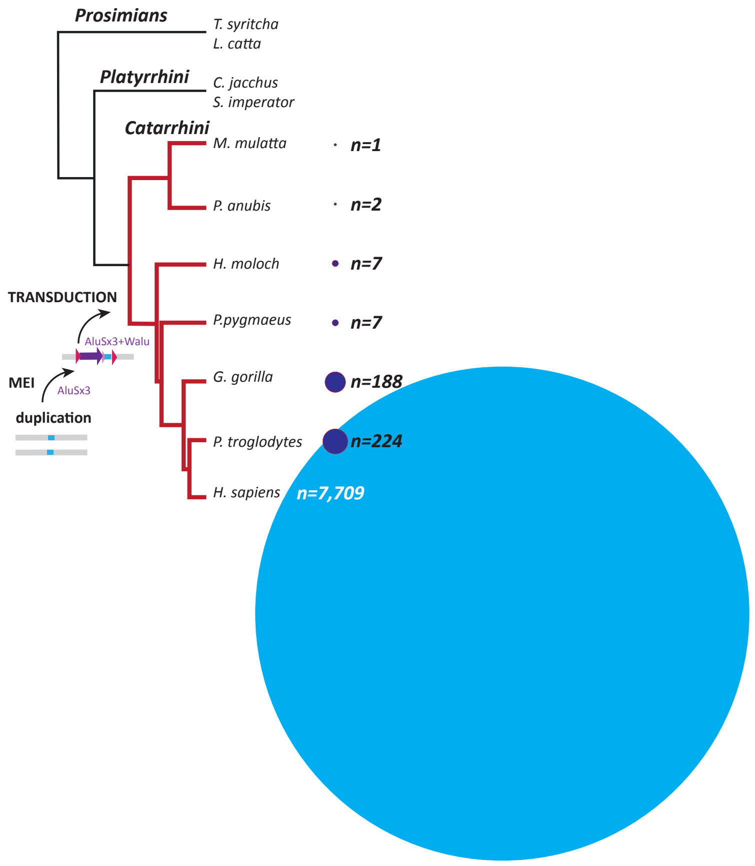


***Fig. S20. Phylogenetic analyses identify the transduction of WaluSat by an AluSx element in the last shared common ancestor with Hominoidea and Catarrhini.*** *Phylogenetic tree showing relationships among primates (Hominoidea, Catarrhini, Platyrrhini, and Prosimians). All primates shown carry the WaluSat novel satellite sequence as a solo locus. Catarrhini and Hominoidea (red branches) show evidence of transduction of WaluSat by an AluSx3 element. Copy numbers of the WaluSat are indicated by a proportional circle to the right of the branch (n = number of WaluSats in a tandem array). This provides evidence for a human specific amplification through the evolution of hominids.*

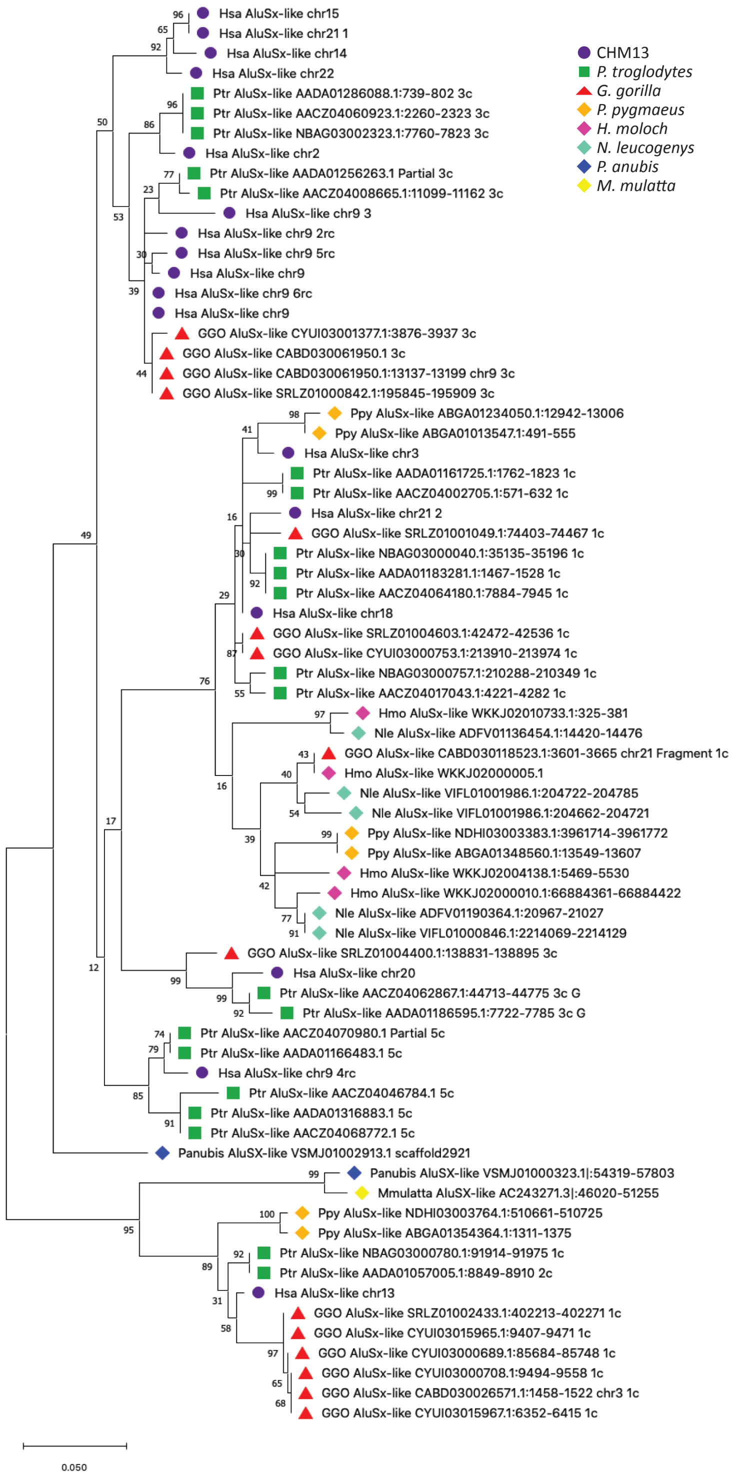


***Figure S21. Maximum-Likelihood tree derived from 71 AluSx-like elements associated with WaluSat monomeric or expanded arrays in Catarrhini.*** *Evolutionary origin and phylogenetic relationship of AluSx elements associated to WaluSat sequences are depicted as present in the Catarrhini common ancestor and having arisen at different times within Catarrhini lineage. The phylogenetic distribution of AluSx-like elements corroborates the hypothesis of*

*multiple transductions events and recent expansion in human acrocentric chromosomes. The evolutionary history of AluSx-like elements was inferred by using the Maximum Likelihood method and T93 model. The tree with the highest log likelihood (-2155.25) is shown. All branches are labeled with the bootstrap values with n = 1000 replicates.*

#### G-quadraplex (G4) analysis

G-quadruplex analysis was done with the GUI version of G4Hunter (*36*) at http://bioinformatics.ibp.cz/. *In silico* tandem array of the 64 nt WaluSat sequence was constructed in fasta. Coordinates for the Chromosome 14 WaluSat array are indicated in Fig. 3E.

### 5. Precision Run-on Sequencing (PRO-seq) and RNA Sequencing analyses

#### Cell permeabilization for CHM13 and RPE-1

For each replicate, adherent cells were washed 2x in cold 1x PBS before adding 5mL of buffer P (10mM Tris-Cl pH 8.0, 10mM KCl, 250mM sucrose, 5mM $MgCl_2$, 1mM EDTA, 0.05% Tween-20, 0.5mM DTT, 10% Glycerol). Cells were scraped, collected and 10uL was removed for cell counting and the remainder was centrifuged at 1000xg for 5 min. 1mL of buffer W (10mM Tris-Cl pH 8.0, 10mM KCl, 250mM sucrose, 5mM MgCl2, 1mM EDTA, 0.5mM DTT, 10% glycerol) was used to gently resuspend cell pellets, before adding an additional 9mL of buffer P, inverting and centrifugation at 1000xg for 5 min. 500uL of buffer F (50mM Tris-CL pH 8.0, 40% glycerol, 5mM $MgCl_2$, 0.1 mM EDTA, 0.5mM DTT) plus 0.5uL of RNase-inhibitor (SuperAse from Ambion) was used to resuspend the cell pellets, followed by another 500uL to wash the tubes; both were pooled together (1mL total), transferred to a 1.5mL tube, and centrifuged at 1000xg for 5 min. Finally, permeabilized cells were resuspended in 57µL of buffer F with 1µL of RNase-inhibitor added before snap-freezing in liquid nitrogen and storage at -80°C.

#### Illumina Library Preparation for CHM13 and RPE-1

PRO-seq libraries were prepared as previously described (*37*) with minor modifications. Approximately 2 x $10^6$ permeabilized cells were mixed with permeabilized *Drosophila* S2 nuclei in all 4-biotin-NTP run-ons (5 x $10^4$ *Dm* nuclei in each replicate), and run-on RNA was extracted with Norgen columns and eluted in 50uL H20. Base-hydrolysis included incubating in 25µL cold 1N NaOH for 10min on ice, followed by the addition of 125uL cold 1M TrisCl pH 6.8, a gentle vortex, and brief spin down before enrichment with streptavidin-beads. Following 3'-ligation and the second bead binding, both end-repair reactions and the 5'-ligation were all performed with nascent RNAs still bound to the beads (*38*). These on-bead reactions were done in a total volume of 20µL with constant rotation before elution from the beads and the subsequent reverse transcription and PCR steps. Test amplifications were performed on 5% of the library and samples were amplified to the ideal number of cycles for final preparation. Following final amplification, libraries were PAGE-purified to remove adapter-dimers and select molecules between 140-650bp in size. Libraries were then sequenced on an Illumina NextSeq 550, producing single-end 75bp reads.

#### Pre-processing and mapping of CHM13 and RPE-1 PRO-seq data

Raw fastq files were first quality trimmed (Phred score >=20) and adapter sequences removed using cutadapt (*39*). Reads below 20nt were removed and remaining reads were reverse complemented using the fastx-toolkit (*40*). *Drosophila* spike-in reads were removed by aligning to the Dm6 genome with bowtie2 (*41*) using "--very-sensitive" options. Remaining reads were then aligned to CHM13 (v1.0) with bowtie2 (*41*) using "-k100" and default options. Sorted bam files were converted to bed files with BEDtools (v2.29.0) (*8*), which were subjected to: 1) unique k-mer filtering and 2) conversion into bedgraphs with BEDtools (v2.29.0) for subsequent normalization with non-mitochondrial alignments to obtain counts in Reads-per-million Mapped (RPMM). Specifically, the raw counts N were normalized using the following equation: RPMM=[N/million_non-mito_alignments]]. Normalized bedgraphs were converted into BigWig files (GenomeBrowser/20180626) for data visualization.

#### SST1 PRO-seq data analyses

SST1 PRO-seq overlap repeat grouping cutoffs (repeats with > 200 and > 2,000 reads overlapping) were determined by plotting the distribution of read overlaps across all SST1 repeats (Table S17). An unpaired *t* test was performed to quantify the statistical significance of differences among SST1 repeats with high v. low read overlap by repeat length, percent divergence, percent insertions, and percent deletions as identified by RepeatMasker and average methylation as determined by (*24*) as described below (Note S9).

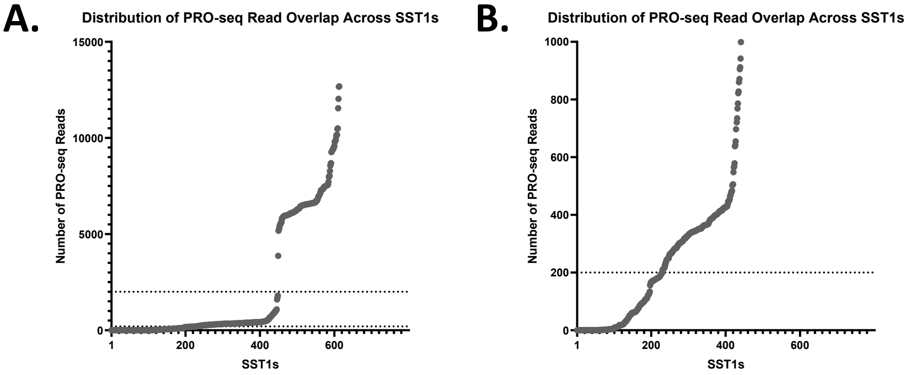


***Fig S22. (A)*** *Distribution of PRO-seq read overlap counts over all annotated SST1 repeats. SST1s with more than 2,000 overlapping PRO-seq reads were grouped (dotted line) and represent purple, yellow, and blue points in Fig. 4C and Fig. S23.* ***(B)*** *200 overlapping PRO-seq reads (dotted line) was used as a cutoff for statistical comparisons.*

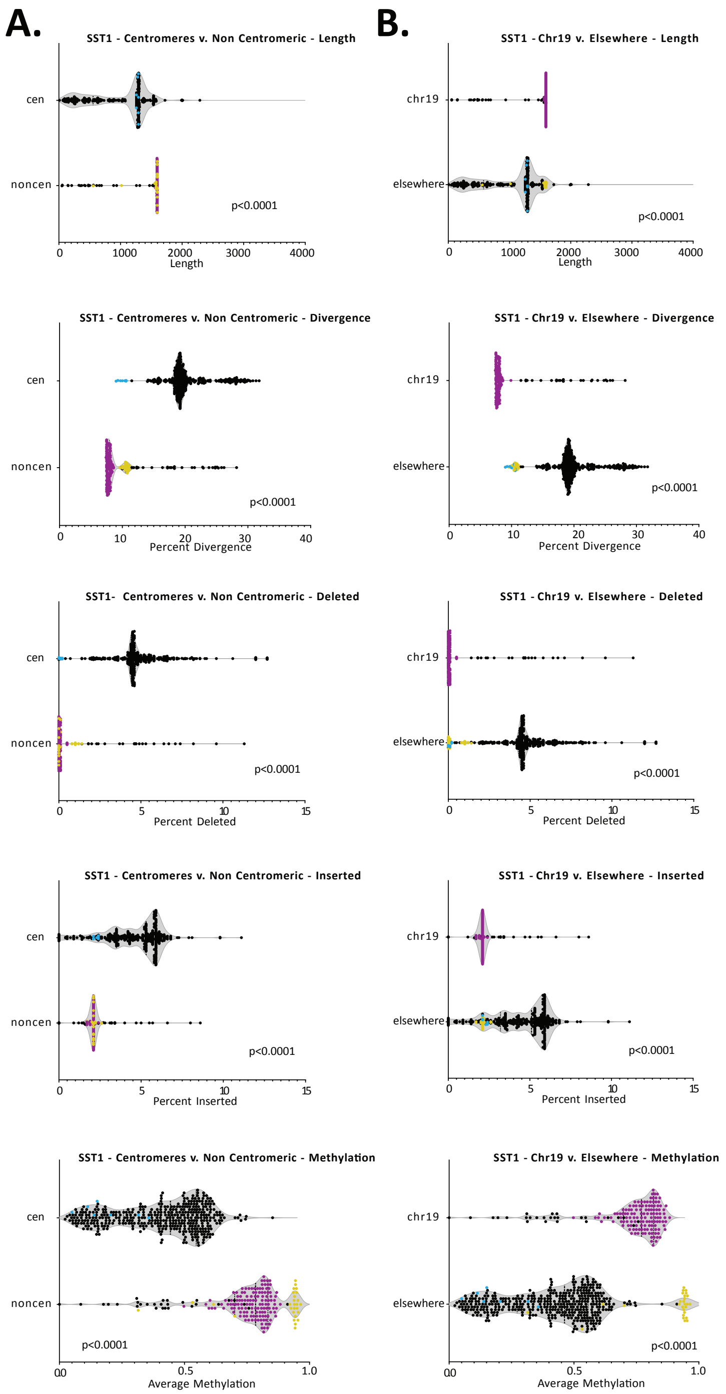

A.
B.
SST1 - Centromeres v. Non Centromeric - Length
cen
noncen
p<0.0001
Length
SST1 - Chr19 v. Elsewhere - Length
chr19
elsewhere
p<0.0001
Length
SST1 - Centromeres v. Non Centromeric - Divergence
Percent Divergence
SST1 - Chr19 v. Elsewhere - Divergence
Percent Divergence
SST1- Centromeres v. Non Centromeric - Deleted
Percent Deleted
SST1 - Chr19 v. Elsewhere - Deleted
Percent Deleted
SST1 - Centromeres v. Non Centromeric - Inserted
Percent Inserted
SST1 - Chr19 v. Elsewhere - Inserted
Percent Inserted
SST1 - Centromeres v. Non Centromeric - Methylation
Average Methylation
SST1 - Chr19 v. Elsewhere - Methylation
Average Methylation

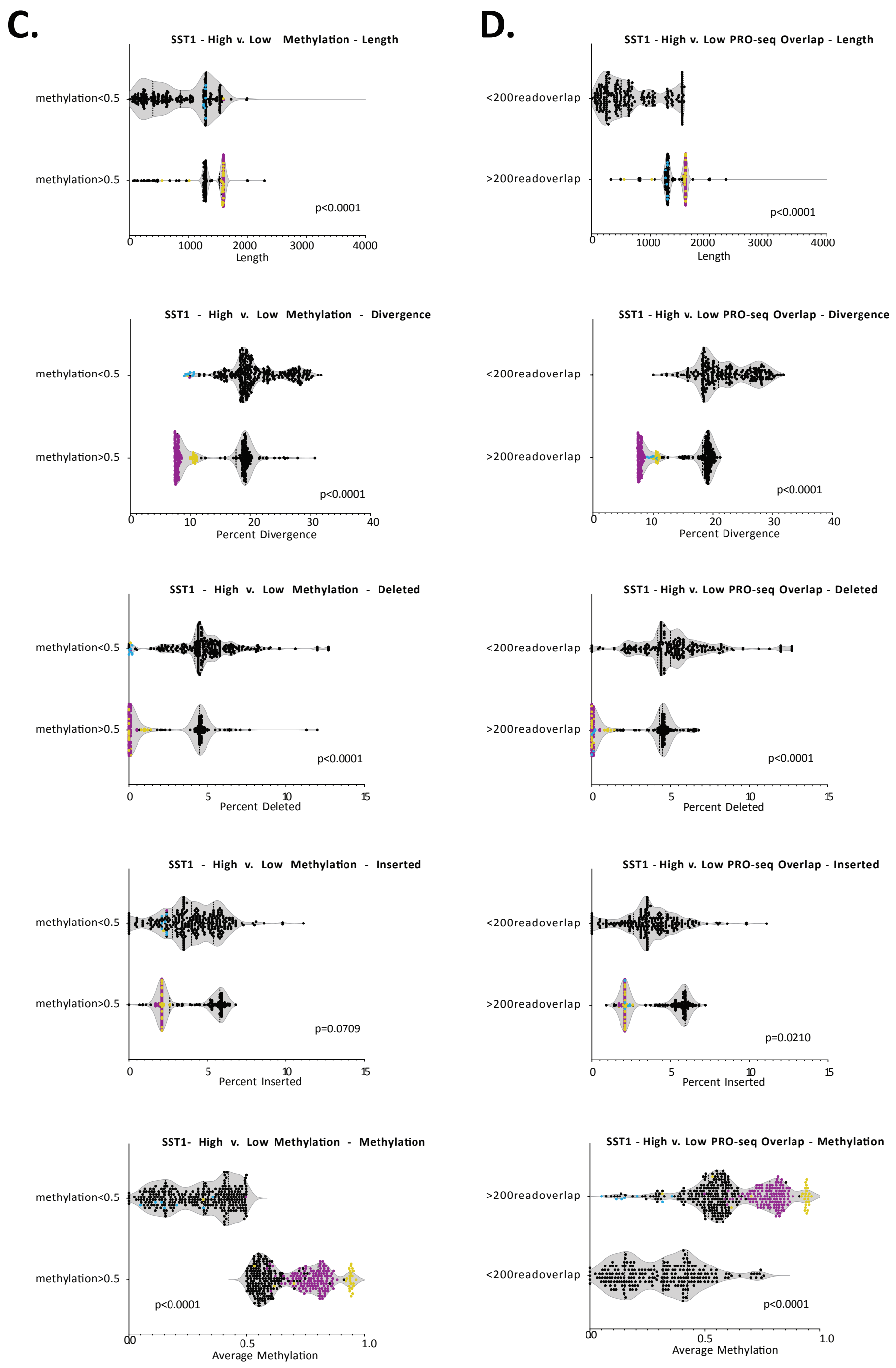


***Fig. S23.*** *Violin plots of SST1 elements show differences in length, divergence, deletions, insertions, and methylation of repeats found in the centromere* ***(A)****, on Chromosome 19* ***(B)****, with varying PRO-seq expression levels* ***(C)*** *and with varying methylation patterns* ***(D)****. Dot colors represent interstitial arrays on Chromosome 19 (purple), Chromosome 4 (yellow), and centromeric monomers on Chromosomes 9, 13, 14, and 21 (blue) with a read overlap higher*

*than 2,000. Non-centromeric SST1s, particularly those on Chromosome 19, are longer, less diverged, and possess higher average methylation than those situated in the centromeres. Similarly, SST1s with > 0.5 methylation and > 200 PRO-seq reads are longer and less diverged than those with lower transcription and lower methylation. All differences are statistically significant (p<0.0001-0.0210).*

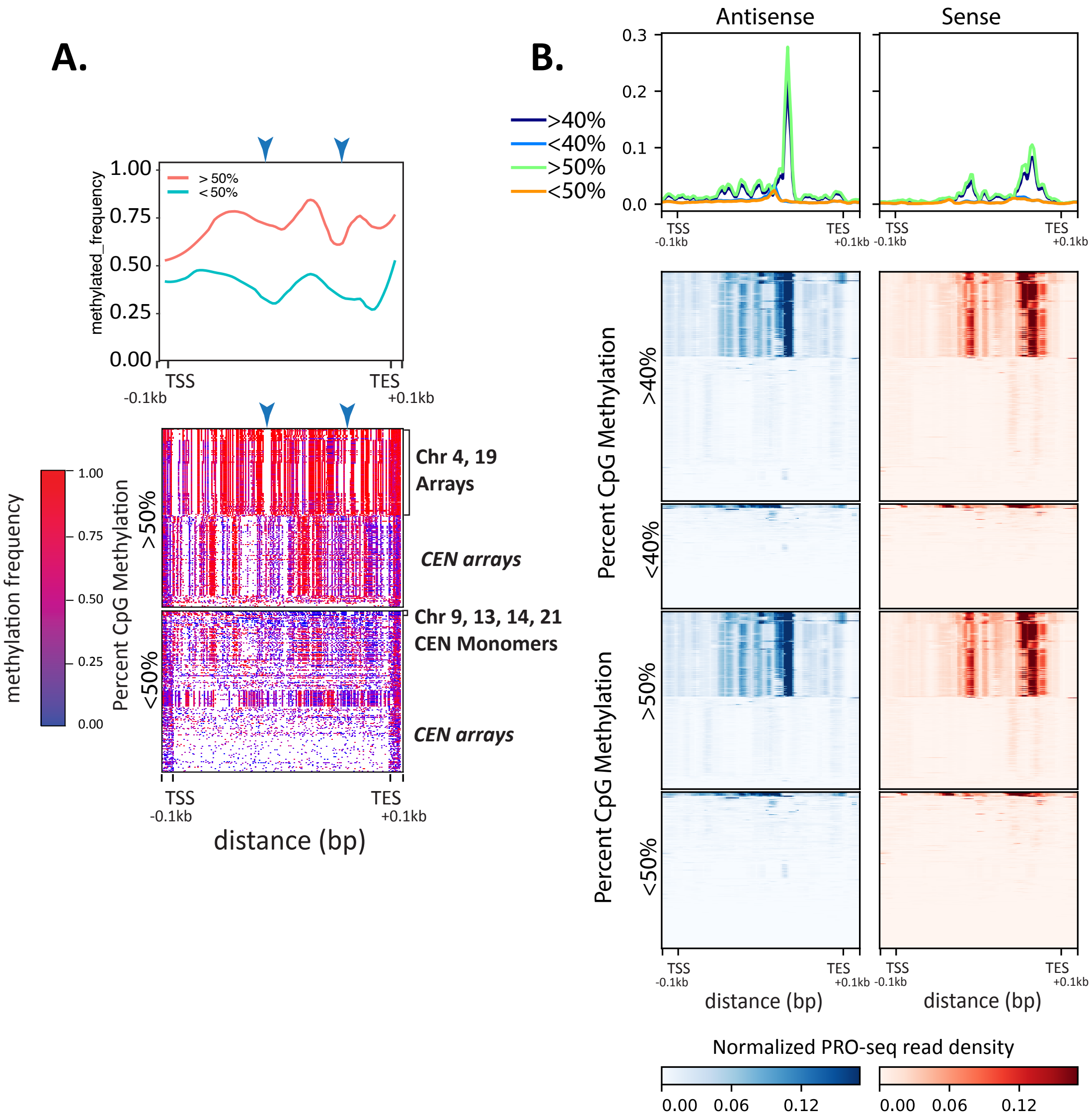


***Fig. S24.*** *CHM13 methylation and stranded PROseq profiles for SST1.* ***(A)*** *Methylation profiles for SST1 grouped by average methylation levels (>50% top, <50% bottom). Each element is scaled to a fixed size, TSS (transcription start site), TES (transcript end site), and ±0.1Kbp are shown (bottom). Clusters of specific SST1 loci are indicated to the right. Methylation frequency scale is on the left.* ***(B)*** *Heatmaps of PRO-seq density (blue scale, normalized reads per million in the antisense direction, red scale, normalized reads per million in the sense direction) grouped by average methylation levels (< and > 50%, < and > 40%).*

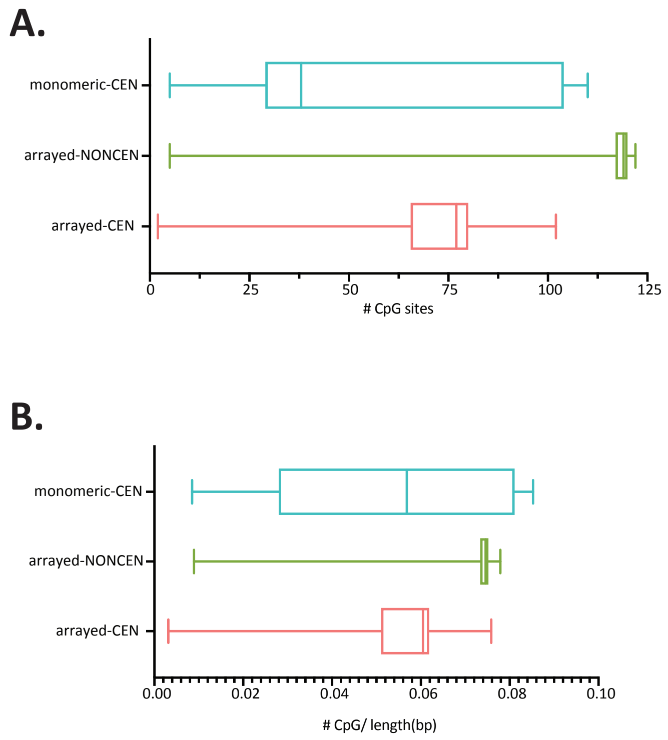


***Fig. S25.*** *Box plots of SST1 repeats showing CG density distribution in* ***(A)*** *CpG density for SST1 repeats 500bp-2Kbp in length delineated by location and repeat density (centromeric (CEN) vs non-centromeric (NONCEN), monomeric vs arrayed).* ***(B)*** *CpG density for SST1 repeats 500bp-2Kbp in length, normalized by element length, delineated by location and density (monomeric vs arrayed, centromeric (CEN) vs non-centromeric (NONCEN).*

### Mitotic Synchronization and Release for HeLa time course

Given the low rate of cell division and synchronization challenges presented by CHM13 cells, HeLa-S3 cells were used as a proxy, noting the caveat that this cell line carries high levels of karyotypic instability (*42*). HeLa-S3 cells at 25-30% confluency were treated with 2mM thymidine for 24 hours, released in fresh medium for 3 hours, then treated with 100ng/mL nocodazole for 12 hours (*43*). Mitotic cells were collected by shake-off, centrifuged and washed in 1x PBS, and then either grown on 15cm dishes in fresh medium for the corresponding time or immediately permeabilized (mitotic sample).

### Cell Cycle Analysis for HeLa time course

Prior to cellular permeabilization, 10% of each sample was removed and fixed in 75% cold (-20°C) ethanol. Cells were then stained with propidium iodide and DNA content was analyzed using a BD FACS Aria II. FCS files were read into R for downstream analyses using the flowCore package. Separately, mitotic HeLa cells were also stained with DAPI and manually analyzed by microscopy in order to differentiate cells in G2 from those properly arrested in prometaphase by level of DNA condensation.

#### Cell Permeabilization for HeLa time course

For each replicate time point, both floating cells in the growth medium and cells removed by scraping in 1x PBS were collected, pooled, and centrifuged at 1000xg for 5 min. Cells were resuspended in 1x PBS and 10% was removed for FACS analysis before completing the wash. 1mL of buffer P (10mM Tris-Cl pH 8.0, 10mM KCl, 250mM sucrose, 5mM $MgCl_2$, 1mM EDTA, 0.05% Tween-20, 0.5mM DTT, 10% Glycerol) was used to gently resuspend cell pellets, before adding an additional 9mL of buffer P and incubating on a shaker for 5min. Permeabilization was assessed with trypan blue and samples not initially permeabilized were again incubated on a shaker with buffer P containing 0.05% NP-40 for 5min. Permeabilized cells were then centrifuged at 1000xg for 5 min before resuspension in 1mL buffer F (50mM Tris-CL pH 8.0, 40% glycerol, 5mM $MgCl_2$, 0.1 mM EDTA, 0.5mM DTT), transferred to a 1.5mL tube, and centrifuged at 1000xg for 5 min. Finally, permeabilized cells were resuspended in 55µL of buffer F and 1µL of RNase-inhibitor was added before snap-freezing in liquid nitrogen and storage at -80°C.

#### Library Preparation for HeLa time course

PRO-seq libraries were prepared as previously described (*37*) with minor modifications. 0.9-4.5 x $10^6$ permeabilized cells were mixed with permeabilized *Drosophila* S2 nuclei in all 4-biotin-NTP run-ons (1 x $10^6$ *Dm* nuclei in each A replicate and 5 x $10^4$ in each B replicate). The rest of the library preparation is the same as in CHM13/RPE PROseq above and libraries were also sequenced on an Illumina NextSeq 550, producing single-end 75bp reads.

#### Pre-processing and mapping HeLa time course data

All data was pre-processed, mapped and post-processed the same way as CHM13 and RPE-1 (see above) with the following few exceptions: 1) alignments to the *D. melanogaster* genome included the "-k 1" option, 2) alignments to CHM13v1.0 included the "--very-sensitive" option and were only done with "-k 100", and 3) normalization was done using a combination of *D. melanogaster* spike-ins (to most accurately compare transcription levels across timepoints), and non-mitochondrial alignments (to uniformly rescale the counts across timepoints so as to obtain final values on a Read-Per-Million-Mapped scale comparable to that of the CHM13 data). Starting from the raw number of reads overlapping a given repeat N, the following equation was used to obtain the normalized counts: *[N/Dmel_norm factor]/(median_across_timepoints[*million_non-mito_ alignments*/Dmel_norm_factor])]*

#### H9 ChRO-seq data availability and pre-processing

External ChRO-seq data (GSE142316) for four different developmental stages (ES, DE, duodenum, ileum) of H9 cells was used for comparison to the CHM13 cell expression data. This data can be found at: https://www.ncbi.nlm.nih.gov/geo/query/acc.cgi?acc=GSE142316 and is initially reported in (*44*). H9 ChRO-seq data was pre-processed using the proseq2.0 pipeline as laid out in and using the code found here: https://github.com/Danko-Lab/proseq2.0 . The script proseq2.0.bsh was used with parameters -SE -G --UMI1=6 --UMI2=6 --Force_deduplicate=FALSE.This script generated adapter-trimmed and deduplicated fastq files which were used as input to Bowtie2 and CASK (Note S8) for repeat composition analysis.

#### Pre-processing, mapping and post-processing of RNA-seq data

Data from paired-end native RNA-seq using oligoDT (*15*) was processed with the same workflow as the CHM13 PRO-seq data, with the following modifications: reads below 100nt were removed, no reverse complement was required (as this is PRO-seq specific), Drosophila spike-ins were not included and therefore, did not need to be removed, and properly paired reads were filtered for with the SAM flag F1548. For the CASK analysis, only mate1 of each replicate was used.

### 6. Statistical analyses and data visualization

BEDtools (v2.29.0) (*8*) map was used to calculate average methylation (*-o mean*) and CpG density (*-o count*) across all repeats in RepeatMaskerV2 (RMv2) and incorporated into the 3D graphs and parallel plots, made using JMP®, Version 16. SAS Institute Inc., Cary, NC, 1989-2019. This method was also used to calculate average methylation for SST1.

BEDtools (v2.29.0) (*8*) was used to intersect SST1 and L1HS repeats with genomic locations (including centromere satellite annotations (*15*)), methylation (*24*), and transcriptional data (Note S6); these data were used to generate repeat groupings (e.g., overlapping a specified satellite annotation; <0.5 average methylation/ >0.5 average methylation, etc.). An unpaired *t* test was performed to quantify the statistical significance of differences among repeat groupings by repeat length, percent divergence, percent insertions, and percent deletions as identified by RepeatMasker and average methylation as determined by (*24*). Percent divergence from the consensus for repeats was taken from the RepeatMasker output. Violin plots were generated via GraphPad Prism software (v9.1.1) GraphPad Software, La Jolla California USA, www.graphpad.com.

Genomic data was visualized for presentation using RIdeogram (v0.2.2) (*45*)and Circos (v0.69-6) (*46*). Circa (v1.2.2) was used to generate segmental duplication ribbon plots (https://omgenomics.com/circa/). JMP®, Version 16. SAS Institute Inc., Cary, NC, 1989-2019, was used to make 3D graphs and parallel plots. Genome browser tracks and CenSat annotations for CHM13 are as described in (*4*, *5*, *15*, *24*).

#### Heatmaps and composite profiles

Count matrices for heatmaps and composite profiles were generated using deepTools2 (*47*). Repeat element groups with a large number of regions were randomly subset to a maximum of 50,000 regions. Data in bigwig format (normalized to either non-mitochondrial or spike-in alignments) was binned using 10bp windows and repeat elements were scaled to an equal size of 1Kbp with the flanking 100bp included in the matrices. Bins summarized the underlying bigwig data by taking the maximum value and composite profiles were created by averaging each bin across all regions in the group.

Methylation heatmaps were generated in R ggplot2 by normalizing repeat size by start and end position and using geom_tile() to plot CpG methylation frequency at each position. Code for methyl heatmaps matching the deepTools2 output can be found: https://github.com/timplab/T2T-Epigenetics/blob/main/TE/DeeptoolsHeatmaps.R

### 7. Identification of full length and immobile TEs; TE aging

The active families in the human genome for SINEs, LINEs, ERVs and retroposons are *Alu*Y, L1HS, HERVK, and SVA_E/F, respectively (*48*). Elements belonging to each family were extracted from the compiled CHM13 and GRCh38 RepeatMasker outputs. Full-length *Alu* belonging to the *Alu*Y family were defined as having a 3' start no shorter than 4 nucleotides (nt), and a 5' end position equal to or greater than 267 nt. Full-length L1Hs sequences were defined as having a length greater than or equal to 6000 nt. SVA_E and SVA_F elements were defined as full-length if the 3' start was no shorter than 50 nt with an end position greater than or equal to 1336 nt, to allow for length variability in the variable number tandem repeat (VNTR) region. ERV elements were considered full length if they met the following criteria: 1) Greater than or equal to 7500 nt (length includes the 5' and 3' flanking LTR structure and internal region); 2) Displayed a sequence structure with 5' and 3' flanking long terminal repeat (LTR) sequence with an internal coding sequence in the middle. Full length element counts and locations can be found in Table S18. These full length sequences were subsequently cross-referenced with PRO-seq data to determine transcriptional activity.

All classes of TEs (excluding DNA transposons) were grouped into relative age groups based on divergence and phylogenetic distribution (*1*, *49–55*), according to Table S19 . LINEs, SINEs and Retroposons were grouped by subfamily, while LTRs were grouped by family.

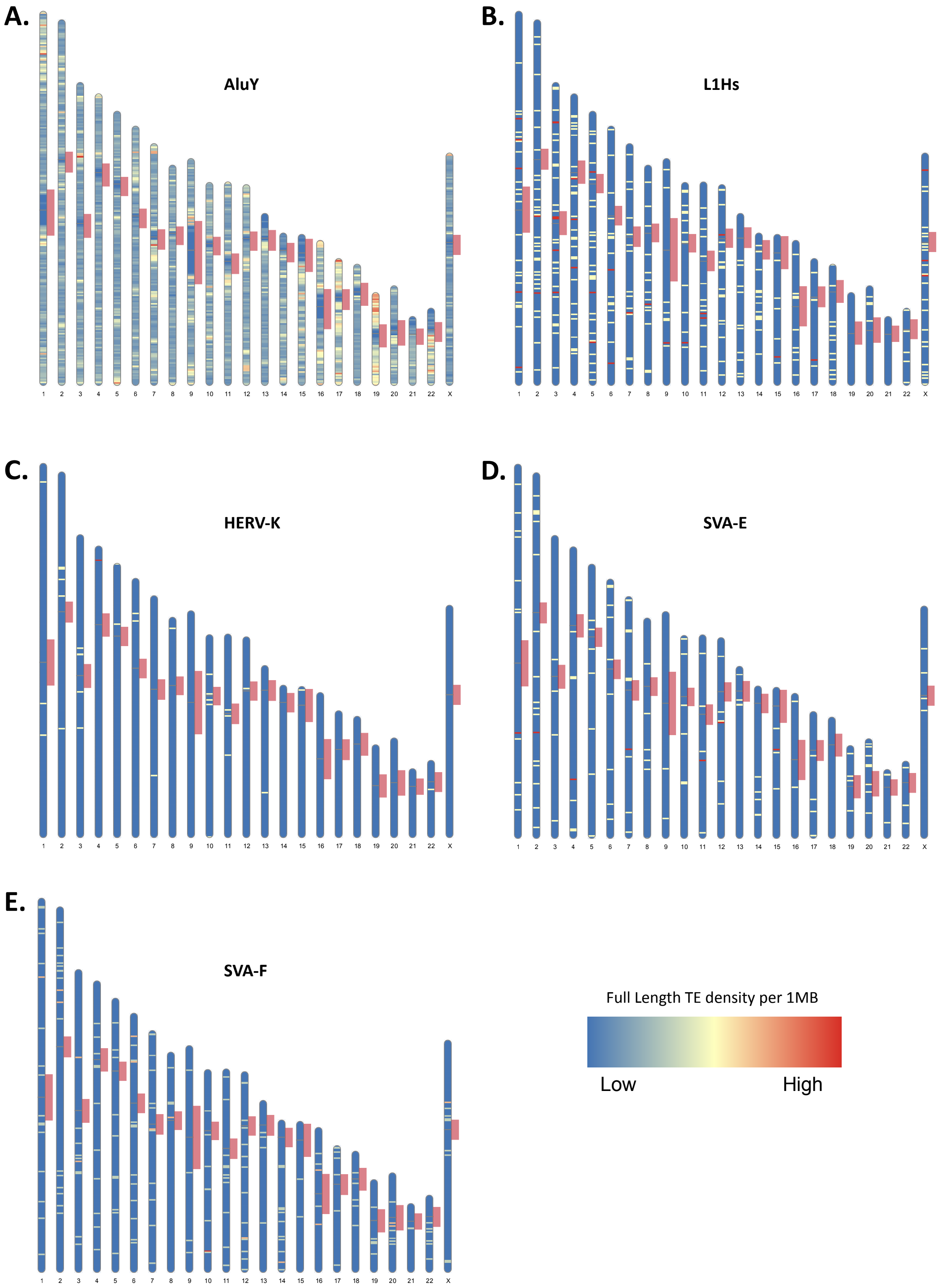


***Fig. S26. Ideogram of density per 1Mbp bins of full length retroelements in CHM13***. *(A) AluY, (B) L1Hs, (C) HERV-K, (D) SVA-E, (E) SVA-F. Centromere regions as per (15)) are shown in red. Density scale from low (blue, zero) to high (red, relative to total copy number).*

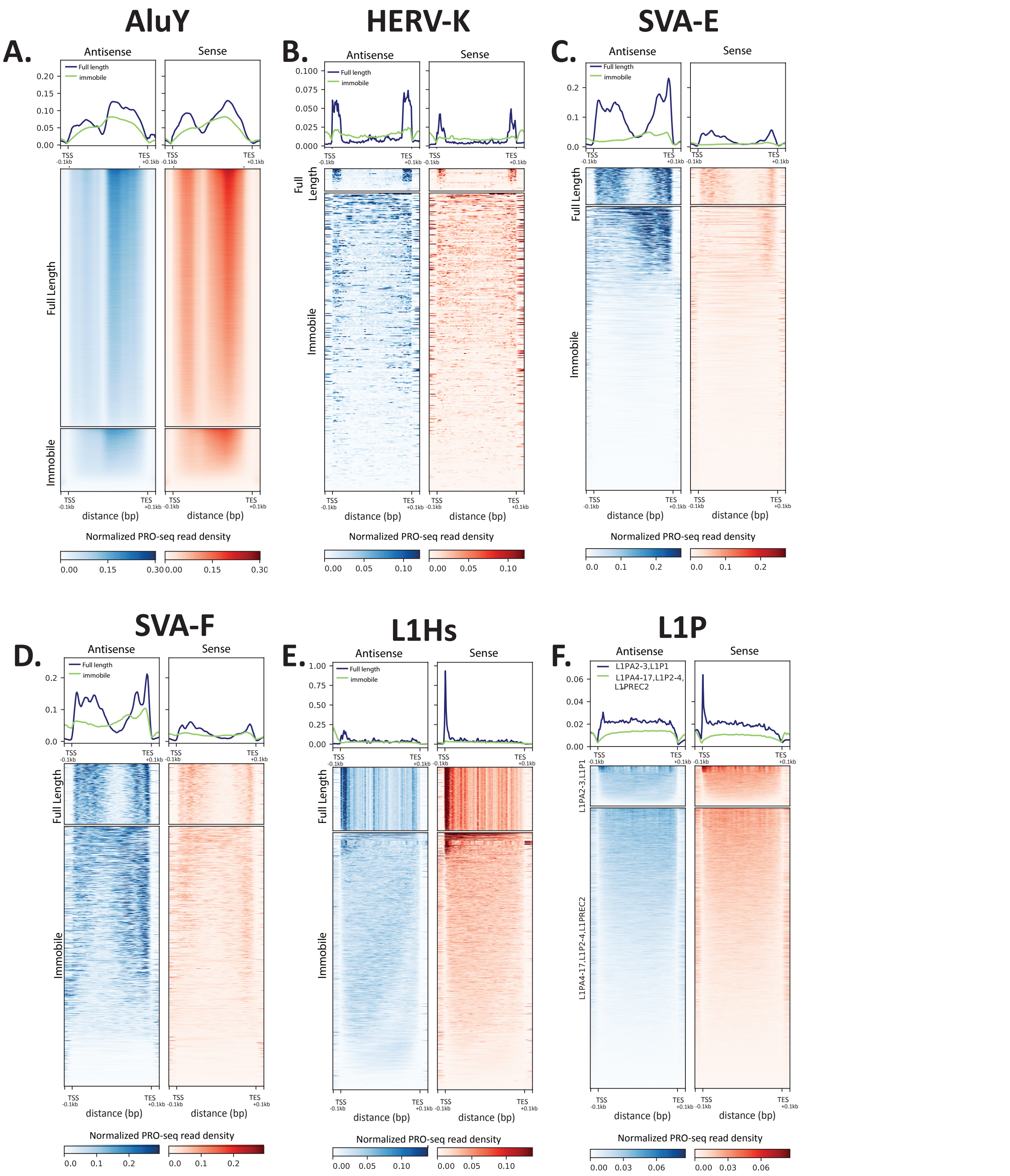


*Fig. S27.* ***Stranded PRO-seq profiles for (A) AluY, (B) HERV-K, (C) SVA-E, (D) SVA-F, (E) L1Hs*** *and* ***(F) L1P*** *subfamilies subdivided into full length and immobile elements. CHM13 PRO-seq density (normalized reads per million for antisense (blue) and sense (red). Average profiles (top line graphs, separated into sense and antisense read density) for TE subfamilies. Each repeat element is scaled to a fixed size; TSS (transcription start site). TES (transcription end site), and ±0.1Kbp are shown (bottom).*

| | | |
|---|---|---|
| **LINE** | L1HS | **youngest** |
| | L1PA2-3, L1P1 | **^** |
| | L1PA4-6, L1P2 | **^** |
| | L1PA7-9, L1P3 | **^** |
| | L1PA10-17, L1P4, L1PREC2 | **^** |
| | L1PB1-3, L1PBa-b, L1P5, L1PB, L1P | **^** |
| | L1PB4 | **^** |
| | L1M, MARE6, X9-LINE, HAL1 | **oldest** |
| **SINE** | AluY | **youngest** |
| | AluS | **^** |
| | AluJ | **oldest** |
| **LTR** | ERVK | **youngest** |
| | ERV1/H | **^** |
| | ERVL | **^** |
| | ERVL-MaLR | **^** |
| | Gypsy-LTR | **oldest** |
| **Retroposon (SVA)** | SVA_F, SVA_E | **youngest** |
| | SVA_D, SVA_C | **^** |
| | SVA_B, SVA_A | **oldest** |

**Table S19.** TE subfamilies (or families for LTR) grouped by approximate evolutionary age. DNA transposons not included in analysis.

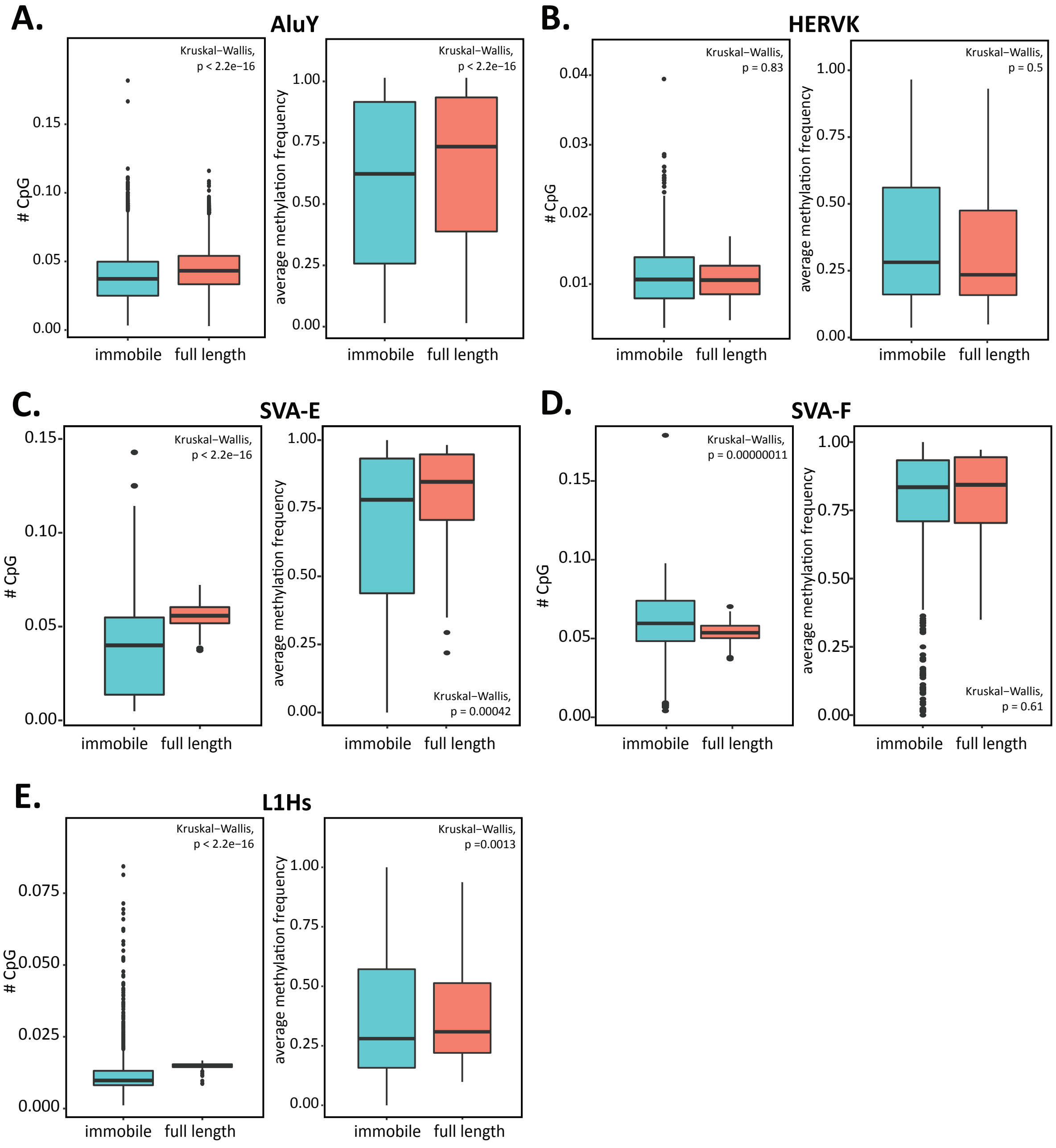


***Fig. S28.*** *Methylation and CpG density boxplot comparisons between full length and immobile* ***(A)*** *AluY,* ***(B)*** *HERV-K,* ***(C)*** *SVA-E,* ***(D)*** *SVA-F,* ***(E)*** *L1Hs elements in CHM13. Methylation frequency was calculated as average methylation per repeat element and CpG density was calculated as number of CpGs normalized to total repeat length. Statistically significant differences were calculated with Kruskal-wallis one-way analysis of variance.*

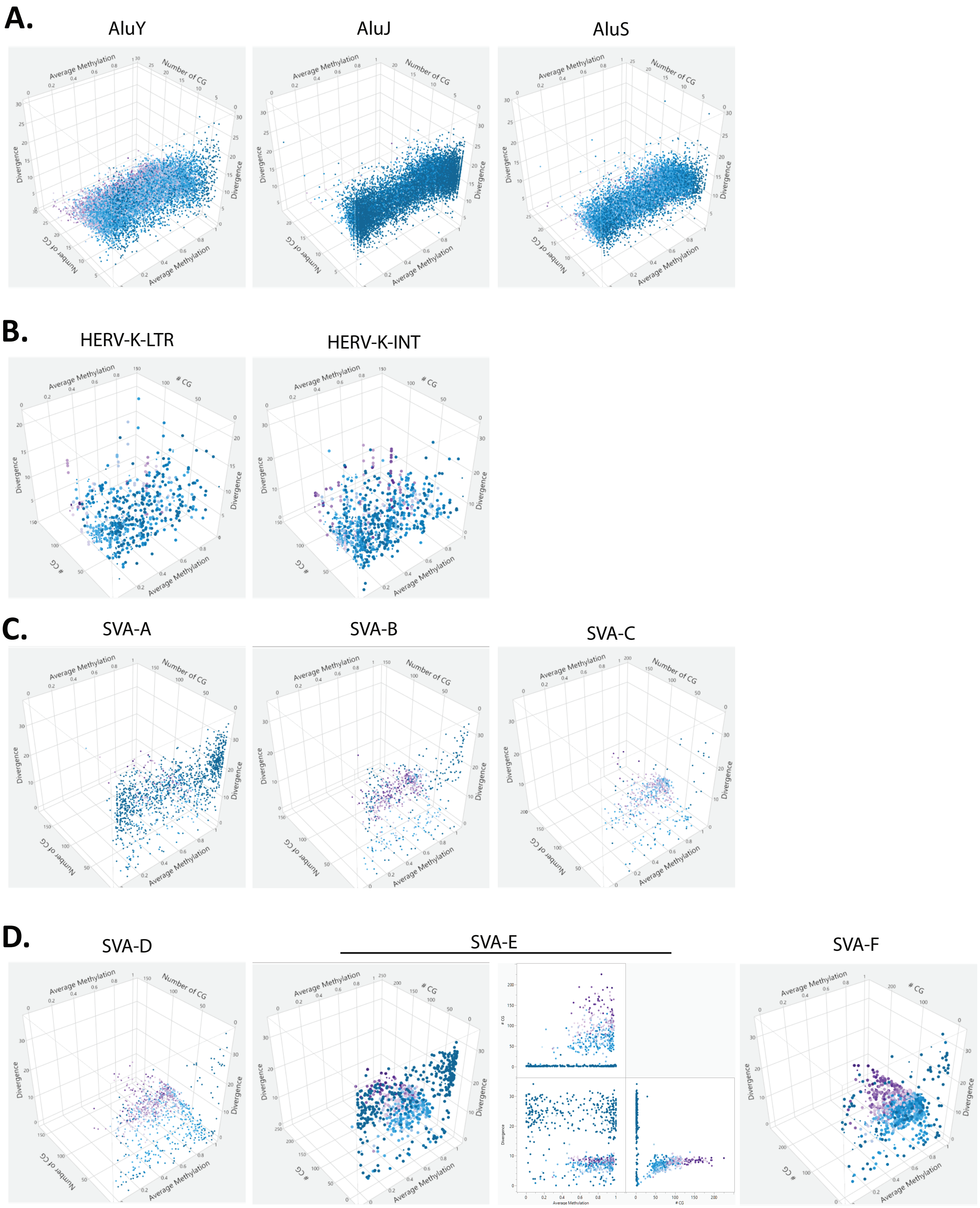
A.
AluY
AluJ
AluS
B.
HERV-K-LTR
HERV-K-INT
C.
SVA-A
SVA-B
SVA-C
D.
SVA-D
SVA-E
SVA-F
Average Methylation
Number of CG
# CG
Divergence

**E.**

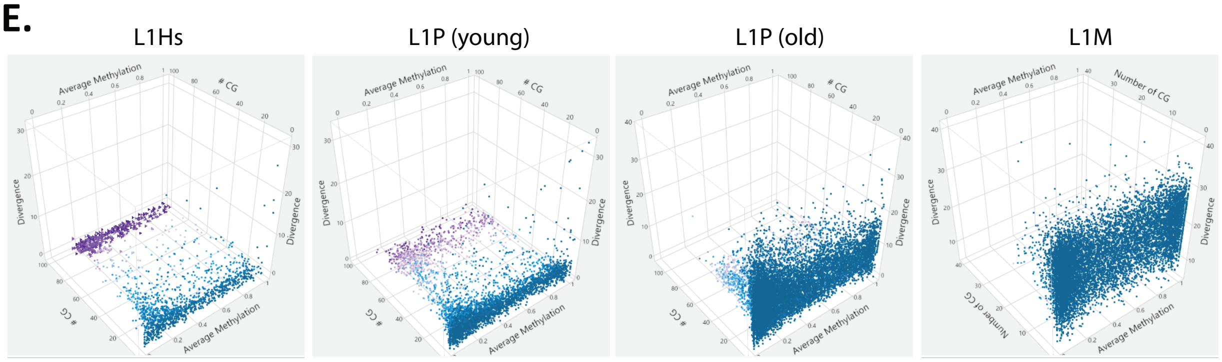


***Fig. S29***. *3D plots for **(A)** AluY, AluJ, AluS, **(B)** HERV-K (divided into LTR and internal regions), **(C)** SVA-A-C, **(D)** SVA-D-F, and **(E)** L1Hs, L1P (young and old), L1M. SVA-E is expanded to demonstrate the individual axes of the 3D plot. Axes represent scaled values for average methylation, # of CpG sites, and divergence from RepeatMasker consensus sequences for each instance of the element. Coloration by the number of overlapping Pro-seq reads where purple represents the highest read overlap and blue the lowest, on the scale matching each plot. These data are the same for Figure 5A-E.*

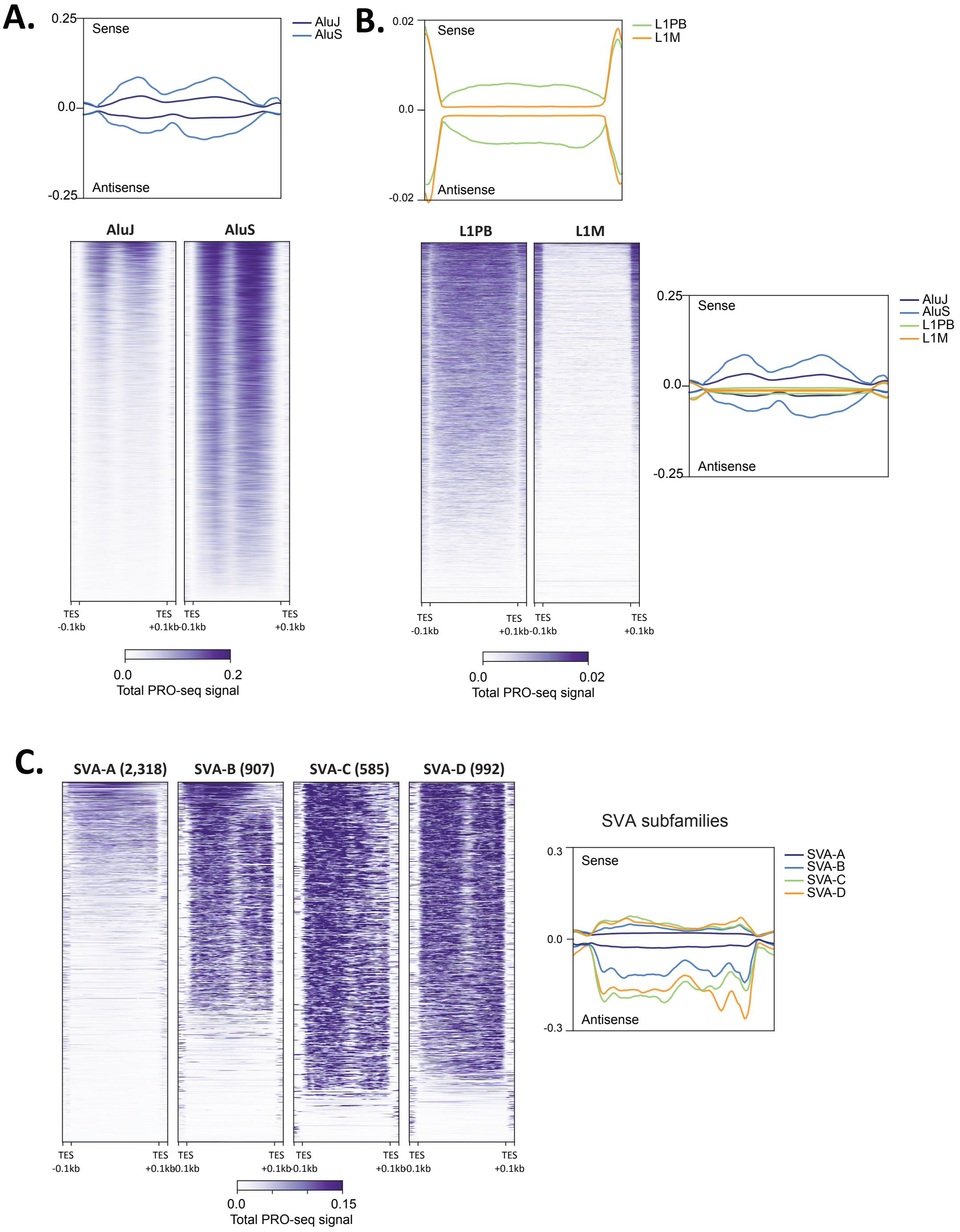


***Fig. S30. PRO-seq profiles for (A) Alu, (B) L1, and (C) SVA subfamilies****.CHM13 PRO-seq density (purple scale, normalized reads per million both sense and antisense aggregated) and average profiles (right line graphs, separated into sense and antisense read density) for TE subfamilies. Each repeat element is scaled to a fixed size; TSS (transcription start site), TES (transcription end site), and ±0.1Kbp are shown (bottom).* ***(C)*** *Number of SVA elements in each subfamily shown at the top of each panel.*

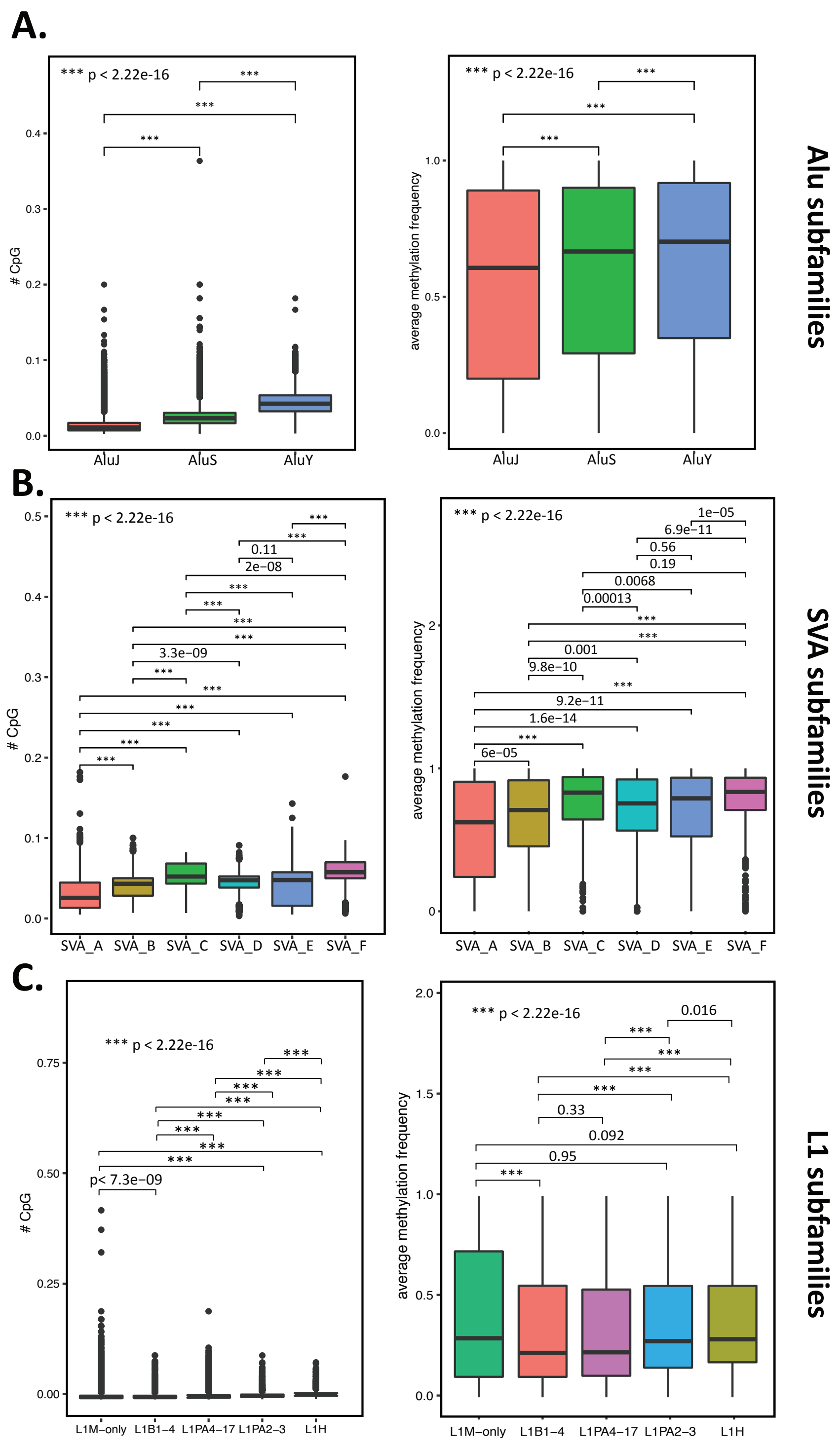


***Fig. S31.*** *Methylation and CpG density boxplots for* ***(A)*** *Alu subfamilies,* ***(B)*** *SVA subfamilies,* ***(C)*** *L1 subfamilies. Methylation frequency was calculated as average methylation per repeat element and CpG density was calculated as number of CpGs normalized to total repeat length. Statistically significant differences were calculated with Kruskal-wallis one-way analysis of variance.*

### 8. Centromere Transcription Analyses

As a complement to the comprehensive TE (herein) and centromere satellite repeat annotations (*15*), we implemented a three-pronged approach to defining the transcriptional landscape of CHM13 centromeres (Fig. S32). In a *mapping-dependent* approach, we mapped PRO-seq and RNA-seq data (Note S5 and below) and intersected reads with unique k-mers based on the CHM13v1.0 assembly and whole genome shotgun (WGS) PCR-free reads (*4*, *56*). As a complement, we implemented a new approach, *mapping-independent* sequence classification (CASK, Fig. S33), which utilized repeat annotations from CHM13 to form a database of k-mers capable of discerning specific repeat types or a refined group of repeats (i.e., ambivalence group). Unmapped PRO-seq and RNA-seq reads were annotated using CASK and the CHM13-dependent k-mer database. Finally, in a *genome-independent* approach, PRO-seq and RNA-seq reads were processed through RepeatMasker using the human Dfam 3.3 library (i.e., not specific to CHM13) (Fig. S32). Simultaneously, RepeatMaskerV2 (RMv2) was intersected (BEDtools v2.29.0) with cenSAT annotations for alpha-satellite only to identify and label those repeats overlapping alpha satellite designated HOR, dHOR, MON, and “none of the above" regions (requiring a minimum of 1bp overlap). This dataset was defined as the alpha-satellite specific RepeatMaskerV2 annotations (RMv2-alpha).

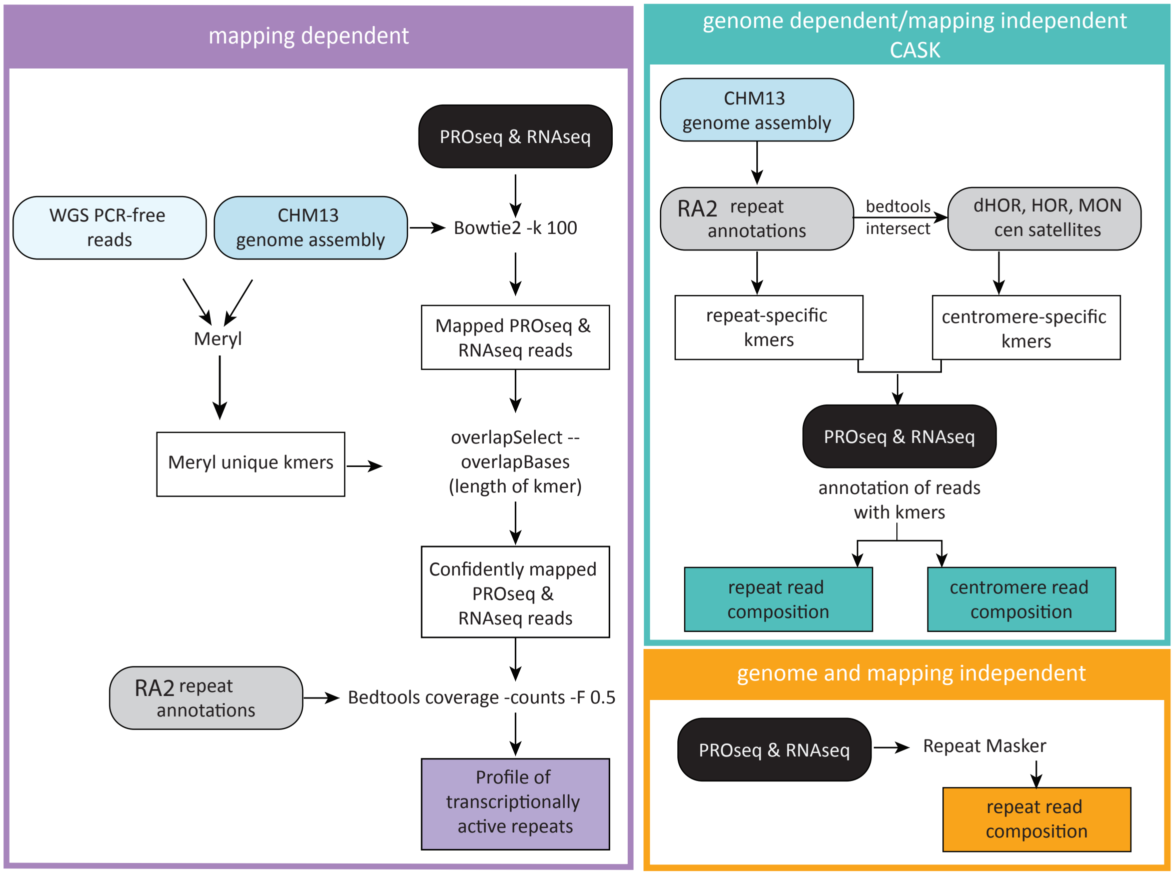


***Fig. S32. Overview of three-pronged repeat transcription pipeline.*** *Two new pipelines were developed to assess repeat transcription levels across the T2T-CHM13 genome, including 1) a mapping-dependent method relying on Bowtie2 and unique genome-wide k-mers (4) (purple) and 2) a mapping- independent method (CASK; see below for details and Fig. S33). Both methods are reliant upon having a genome assembly. Repeatmasker was simultaneously used to determine repeat content in the reads as a genome and mapping-independent method as per (57).*

**Mapping dependent PRO-seq analyses**

The benefit of having a complete, high-quality long-read assembly such as CHM13v1.0, allows for the generation of genome-wide unique k-mers spanning even the most repetitive regions of the genome. K-mer sizes of both 21 and 51bp were used in these analyses (*56*). These unique k-mers (generated through Meryl) were based on the CHM13v1.0 assembly itself and whole genome shotgun (WGS) PCR-free reads (*4*) for increased confidence in their uniqueness. Where indicated, bed files of the mapped reads (as described above, Note S5) were filtered through these unique k-mers using overlapSelect with the setting "-overlapBases=XXbp", where XX is equal to the length of the unique k-mer being overlapped (21bp or 51bp). This required that a minimum of the entire length of the unique k-mer must overlap a given read in order for that read to be retained. The 51-mer filtered data was primarily used over the unique 21-mers since the larger the k-mer, the more abundant they are throughout the genome, and hence, more reads are retained. Alternatively, we did not use a k-mer size larger than 51bp because an average PRO-seq read is ~55bp; thus, a longer k-mer size would result in the loss of most of the reads. RNA-seq reads were filtered through the same methods for consistency with the PRO-seq data (even though these reads are longer). Following filtering through unique k-mers, bed files were used for counting reads overlapping repeats or alpha-satellites and for bigwig generation for visualization, as described above in the "Pre-processing, mapping and post-processing of reads" section (Note S5).

**CASK: Classification of Ambivalent Sequences using k-mers**

We used CASK (Classification of Ambivalent Sequences using K-mers), a mapping-independent method to identify reads originating from repeat elements using their k-mer composition (Fig. S33A). Briefly, the genomic location of all repeats and their type annotation (e.g., L1H, L1P, *Alu*Y, etc.) were extracted from the final CHM13v1.0 RepeatMaskerv2 annotations (RMv2, Note S1). For each repeat type, the genomic sequences of all the instances of that repeat were extracted into a type-specific fasta file. These fasta files were input into KMC (*58*) to generate for each repeat type a type-specific k-mer database, consisting of all the k-mers (k=25) found across all instances of that repeat. Each type-specific k-mer database was then filtered to remove k-mers also present in parts of the genome that did not overlap with any repeat elements (these k-mers have limited usefulness for the purpose of identifying reads originating from repeats and could lead to false identification of repeats). Note that while each k-mer is represented only once in any given type-specific database, many k-mers are not unique genome-wide and may be found multiple times within the same or different instances of the repeat. Additionally, many k-mers are not exclusive to a given repeat type and may be shared across different repeat types and the corresponding type-specific kmer databases (e.g., a k-mer found in L1H may also be found in L1S, L1P, etc.). For each k-mer, we defined its ambivalence group as the set of all repeat types within which this k-mer was found. Using a custom pipeline, the type-specific k-mer databases were combined into a single "annotated k-mer database" listing all the k-mers found across repeats and their corresponding ambivalence group.

Starting from the trimmed and deduplicated fastqs (PRO-seq, RNA-seq, ChRO-seq, Note S5), sequencing reads containing one or more k-mers matching a k-mer in the annotated k-mer database were extracted using BBduk. For each read, we then computed the intersection of the ambivalence groups of all the matching k-mers within the read. This intersection can be interpreted as a consensus repeat assignment from all of the k-mers in the read. If the intersection contained a single repeat-type (e.g., L1P), the read was assigned to that repeat. If the intersection contained multiple repeat types, the read was annotated as ambivalent, and the possible set of repeat types for this read was recorded (e.g., an ambivalent read could receive an assignment {L1H or L1P}). Such ambivalent reads were used to compute upper bounds on the number of reads originating from a given repeat type (e.g., Fig. S33). CASK data shown without error bars ignore reads with ambivalent assignments and thus represent *lower bound* estimates of the repeat expression. Finally, although this scenario was rare (Fig. S33), if the intersection was empty (e.g., if one k-mer in the read was unique to L1P and another k-mer was unique to L1Hs), the read was annotated as containing “conflicting k-mers” and discarded from further analysis.

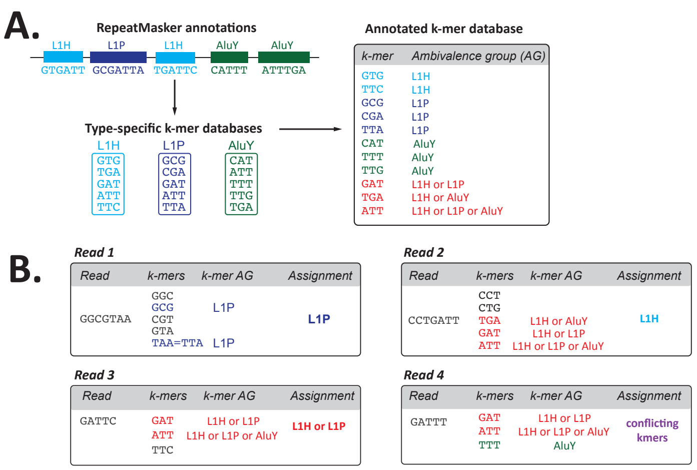


***Fig. S33. The CASK (Classification of Ambivalent Sequences using k-mers) algorithm. (A)*** *Main steps of the algorithm and* ***(B)*** *examples showing the repeat-type assignment for four reads with different k-mers composition.*

#### RepeatMasking of PROseq and RNAseq reads

RepeatMasker (v4.1.2-p1) was run on the trimmed reads of the individual replicates of the CHM13 PROseq and RNAseq datasets using a library consisting of the Dfam 3.3 database plus the new entries discovered as part of the TE analysis of the CHM13v1.0 genome assembly (RMv2, Fig. S1). The resulting RepeatMasker output files were then summarized using RM_summarizer.pl (perl v5.30.1; https://github.com/gabriellehartley/CENP-Gibbon-Analysis) to obtain the number of reads containing each repeat type. For the paired-end RNAseq datasets, mate1 and mate2 were run individually and the counts were summed. For both RNA-seq and PRO-seq, the relative abundance of each repeat was similar across replicates, and thus counts from both replicates were summed. A small percentage of reads that were retained had >1 repeat designation across the length of the read (0.21% of all PRO-seq reads, 1.32% of all RNA-seq reads).

#### TE Embeds within cenSAT annotations

As per the RMv2-alpha file generation above, the RepeatMasker AnnotationV2 was intersected (BEDtools v2.29.0) with all cenSAT annotations to identify and label those repeats overlapping any of the major satellite groups (e.g., alpha, beta, HSAT). A minimum of 1bp overlap was used to assess whether a TE was embedded and/or at the edge of one of these satellite regions.

#### Detecting Repeat Transcription - Method Comparisons

BEDtools (*8*) coverage was used to obtain counts of reads overlapping repeats defined in RMv2 across all mapping methods, requiring at least 50% of the read (using “-counts -F 0.5”) to overlap the repeat element. This method was also used to determine how many *repeats* had reads overlapping (Fig. S34-S35, Table S20). Alpha-satellite specific RepeatMaskerV2 annotations (RMv2-alpha) were used to obtain counts of reads that overlap repeats in the same manner as above (BEDtools coverage -counts -F 0.5). Since 50% of the length of a pre-processed PRO-seq read is ~25-30bp, and the CASK k-mer length is 25bp, these parameters are roughly equivalent for this comparative analysis. The relative abundance of each repeat was similar across replicates, and thus counts from both replicates were summed.

To compare all approaches, we first used different bowtie mapping parameters (default, k-100 and k-100 filtered for unique k-mers in CHM13) on PRO-seq and RNA-seq datasets (Fig. S34). For PRO-seq datasets, default parameters that report only the best mapped location ("BT2 default") and mapping parameters that support multi-mappers combined with intersecting those reads to only report those that overlap with a unique 51mer ("BT2 k100 51mer"), thus representing high confidence locations, show largely concordant repeat calls (Fig. S34). In contrast, RNA-seq datasets show largely concordant repeat calls for the "BT2 k100" and "BT2 k100 51mer" with a larger difference in calls compared to the BT2 default parameters (Fig. S34). Thus, different mapping approaches impact data interpretation for different types of transcription datasets. Overall, BT2 default parameters report fewer repeat calls, while Bt2 k100 with a 51mer unique k-mer overlap represent the highest confidence, and strictest, repeat calls.

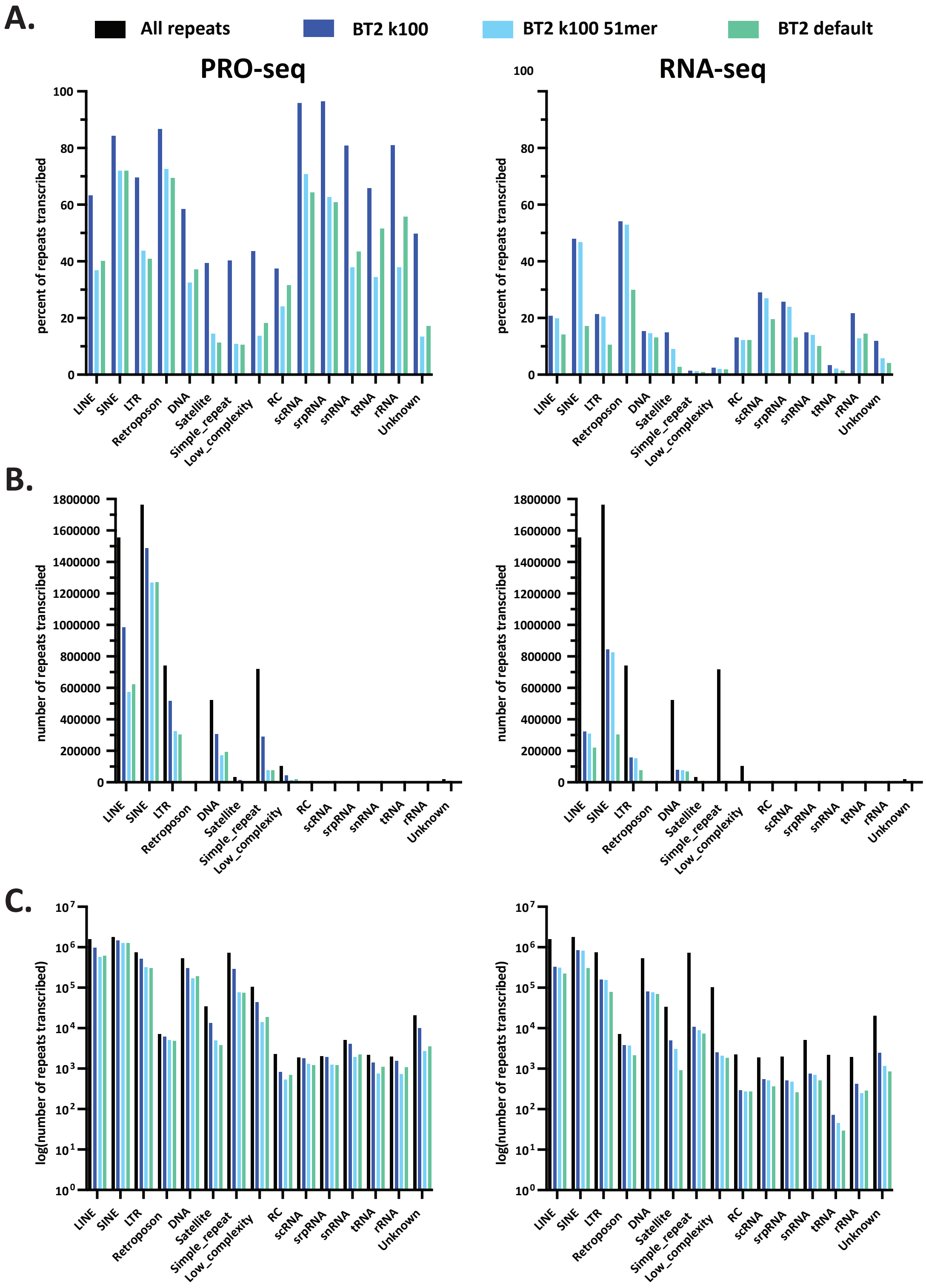


***Fig. S34.*** *Profile of transcriptionally active repeats across Bowtie2 mapping methods for PRO-seq (Left) and RNA-seq (right) data shown as* ***(A)*** *percent of repeats transcribed,* ***(B)*** *number of repeats transcribed, and* ***(C)*** *Log transformed number of repeats transcribed.*

When comparing across all repeat transcript annotation methods (Fig. S32), we find that CASK, RM and BT2 default annotations for PRO-seq data were largely concordant, while repeat annotations for RNA-seq data from CHM13 were variable across methods (Fig. S35). Given that PRO-seq captures nascent transcription while RNA-seq cannot distinguish new transcripts from stable and accumulating transcripts, combined with variable repeat calls across methods, indicates that PRO-seq provides a more robust representation of active transcription. Of note, while SINEs were the predominantly transcribed repeat across all datasets irrespective of the method employed (BT2, CASK, RM), allowing multi-mappers BT2 k100 or BT2 k100 51mer filtered, resulted in SINE read counts increasing, likely due to their high abundance in the human genome.

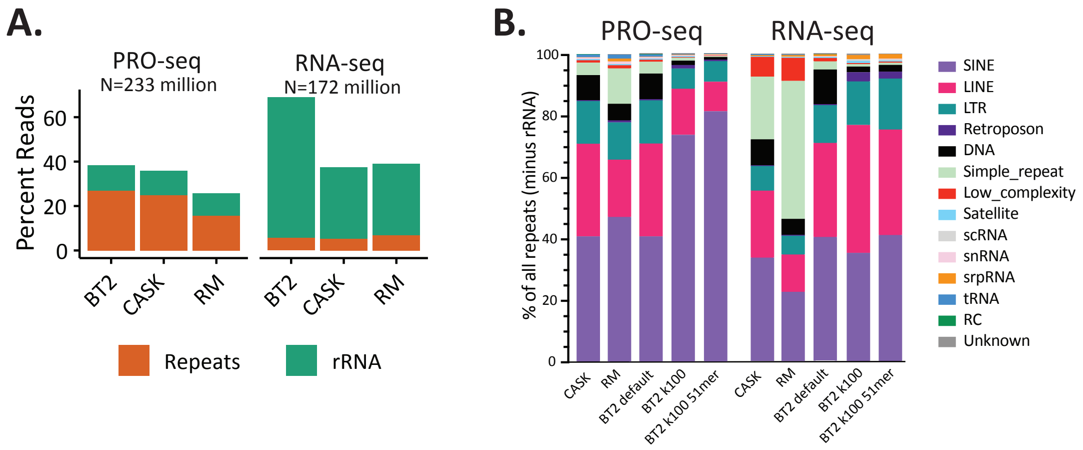


***Fig. S35. Transcription of repeats assayed by PRO-seq and RNA-seq. (A)*** *Percentage of reads assigned to repeat elements or rRNA by BT2, CASK, and RepeatMasker (RM) in the CHM13 PRO-seq and RNA-seq datasets (replicates combined).* ***(B)*** *Relative abundance of the 14 repeat classes defined in RMv2 (excluding rRNA), as quantified by BT2, CASK, and RM. Three settings were used for BT2 differing in the handling of multi-mappers: default (default BT2 options, resulting in single alignment randomly chosen for multi-mappers), k100 (-k 100 option, up to 100 alignments chosen for multi-mappers), k100 51mer (-k 100 option, but reads were post-filtered to overlap a genome-wide unique 51-mer).*

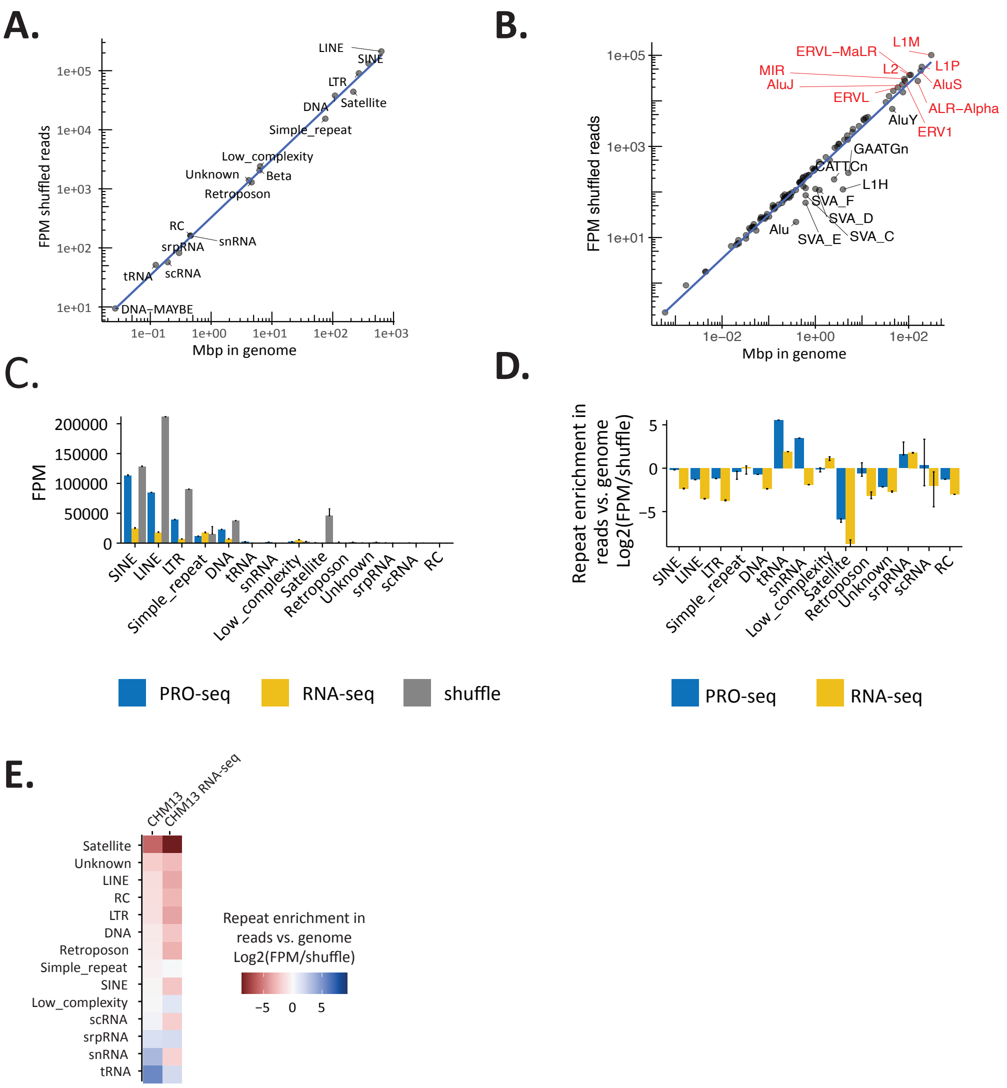


***Fig. S36. Repeat expression normalized to genome content. (A)*** *Linear scaling between the genomic abundance of each repeat family* ***(A)*** *and class* ***(B)*** *(Mbp), and the FPM values obtained by CASK for 100 million 65bp reads sampled from random positions in the genome (shuffled reads). Top 10 class by Mbp in CHM13 are indicated in red in* ***(B). (C)*** *Comparison between the observed repeat expression in CHM13 and the expression that would be expected from reads originating from random loci (shuffle), both quantified by CASK. Error bars represent upper bounds on the repeat expression, obtained by including reads with ambivalent CASK repeat assignment in the tally.* ***(D)*** *Repeat enrichment in the CHM13 transcriptome vs. CHM13 genome defined as the $log_2$ ratio of observed expression over shuffle. Error bars represent lower and upper bounds on the ratio, obtained by including shuffled or true reads with ambivalent CASK repeat assignment, respectively, in the tally.* ***(E)*** *Repeat enrichment across PRO-seq and RNA-seq data ranked from least (red) to most enriched (blue) in CHM13.*

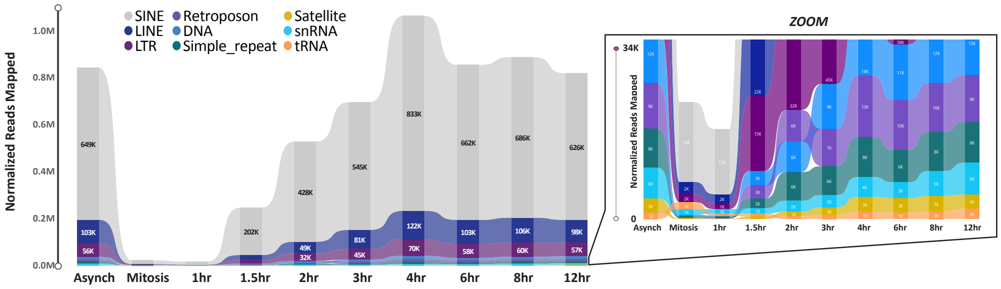


***Fig. S37.*** *Ribbon plots of repeat abundance in normalized PRO-seq data (shown as Reads per Million RPM) assessed by BT2 (-k100) in asynchronous and synchronized HeLa cells collected at time points across the cell cycle (key in inset). Zoom shows the reads for the lower range of expressed repeats, including all satellites classified in CHM13 (tan).*

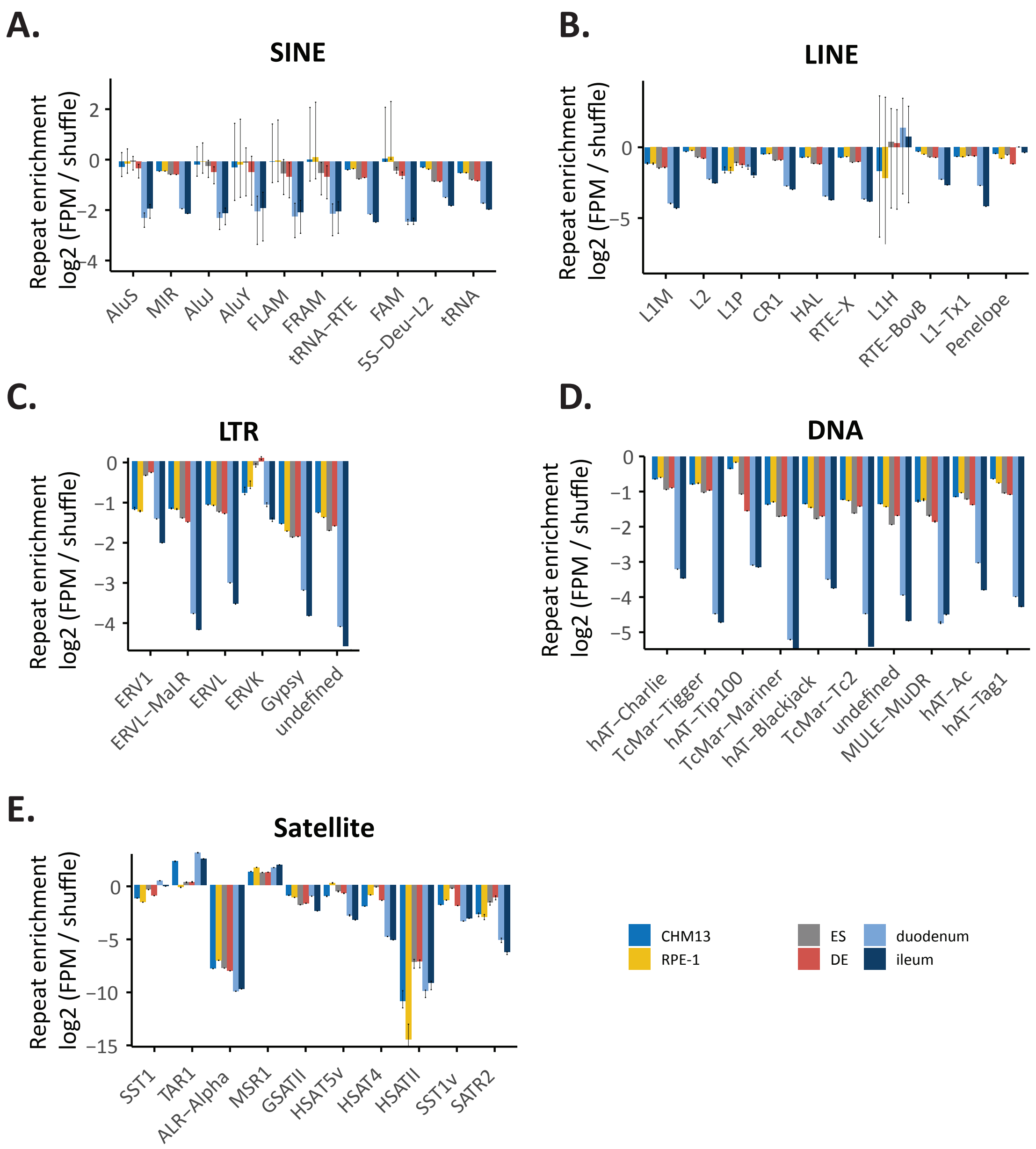


***Fig. S38**. **Repeat enrichment by repeat family in cell specific transcriptomes** (as per key, lower right) vs. CHM13 genome for **(A)** SINE, **(B)** LINE, **(C)** LTR, **(D)** DNA elements, and **(E)** Satellite repeat classes. Within each class, repeats are displayed by decreasing abundance (average expression across cell lines) from left to right, with the top 10 repeat families in each category shown. The repeat enrichment is defined as in Fig. S36D.*

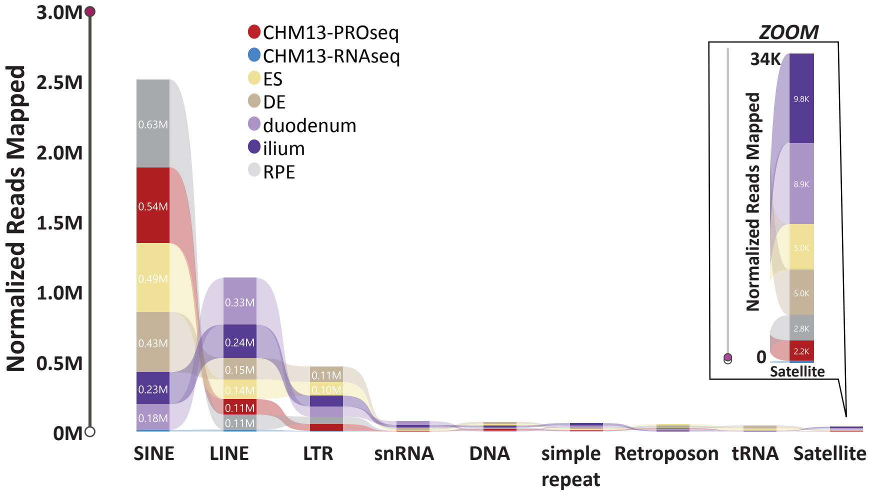


***Fig. S39.*** *Ribbon plots of repeat abundance in normalized PRO-seq data (shown as Reads per Million RPM) assessed by BT2 (-k100) across cell types (key in inset). Zoom shows the reads for the lower range of expressed repeats, including all satellites classified in CHM13.*

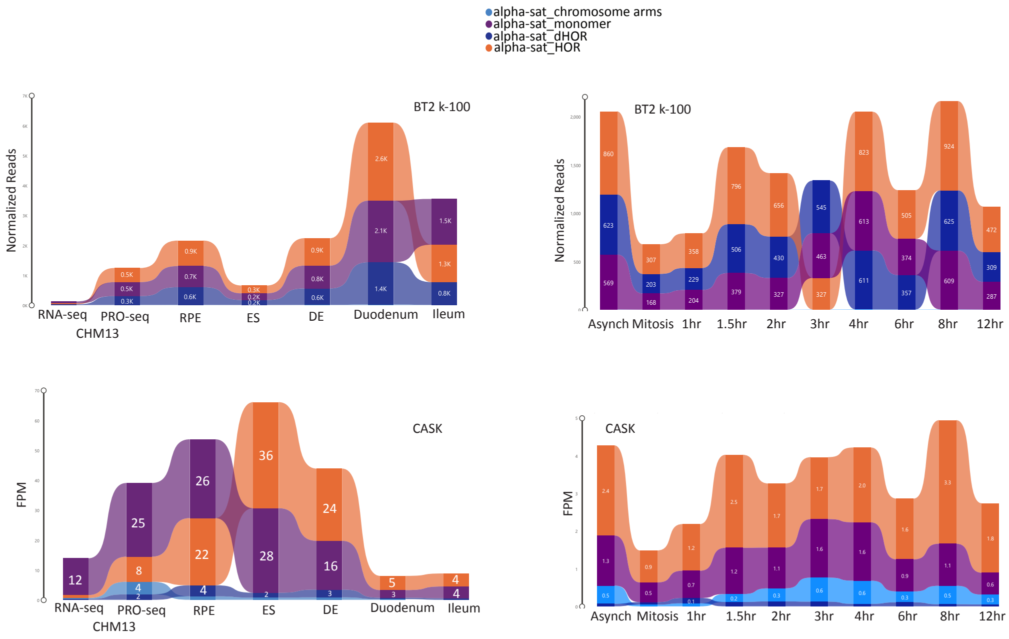


***Fig. S40.*** *Ribbon plots of alpha satellite repeat abundance in PRO-seq data (shown as Reads per Million RPM) assessed by BT2 (-k100)* ***(top)*** *and CASK* ***(bottom)*** *across cell types and developmental stages* ***(A)*** *and time points* ***(B)*** *(key at top).*

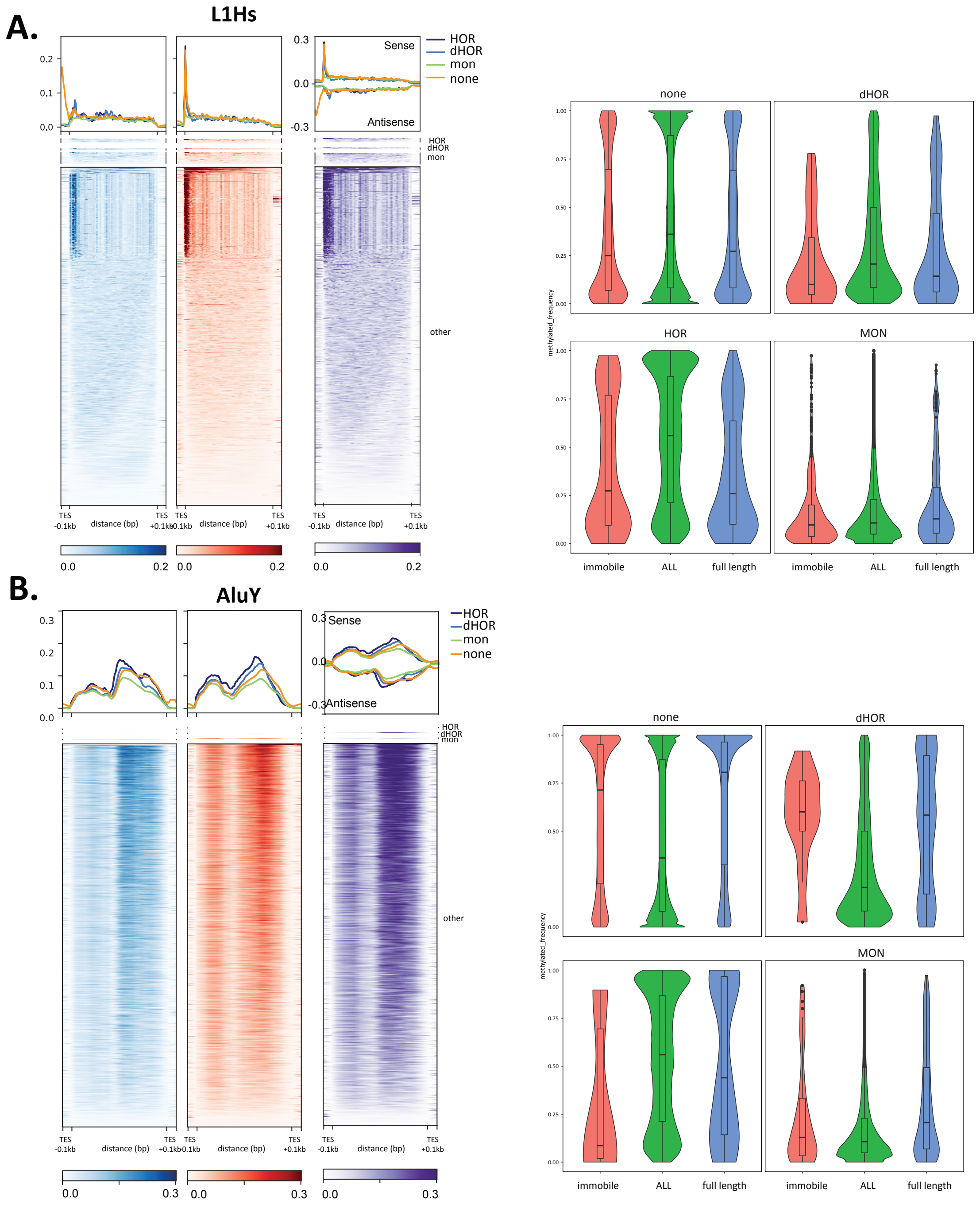


***Fig S41. Transcription and methylation profiles of embedded elements. (A)*** *L1Hs and* ***(B)*** *AluY.* ***Left Panel:*** *Heatmaps of CHM13 PRO-seq density (red scale, normalized reads per million) and average profiles (upper panels). HOR, dHOR and monomeric embeds are separated from all non-embedded elements (subsampled for AluY). Each repeat element is scaled to a fixed size; TSS (transcription start site), TES (transcription end site), and ±0.1Kbp are shown.* ***Right panel:*** *Violin plots of methylated CpGs for TEs grouped by their potential for mobility and compared across embedded category (dHOR, HOR, monomeric) vs not embedded.*

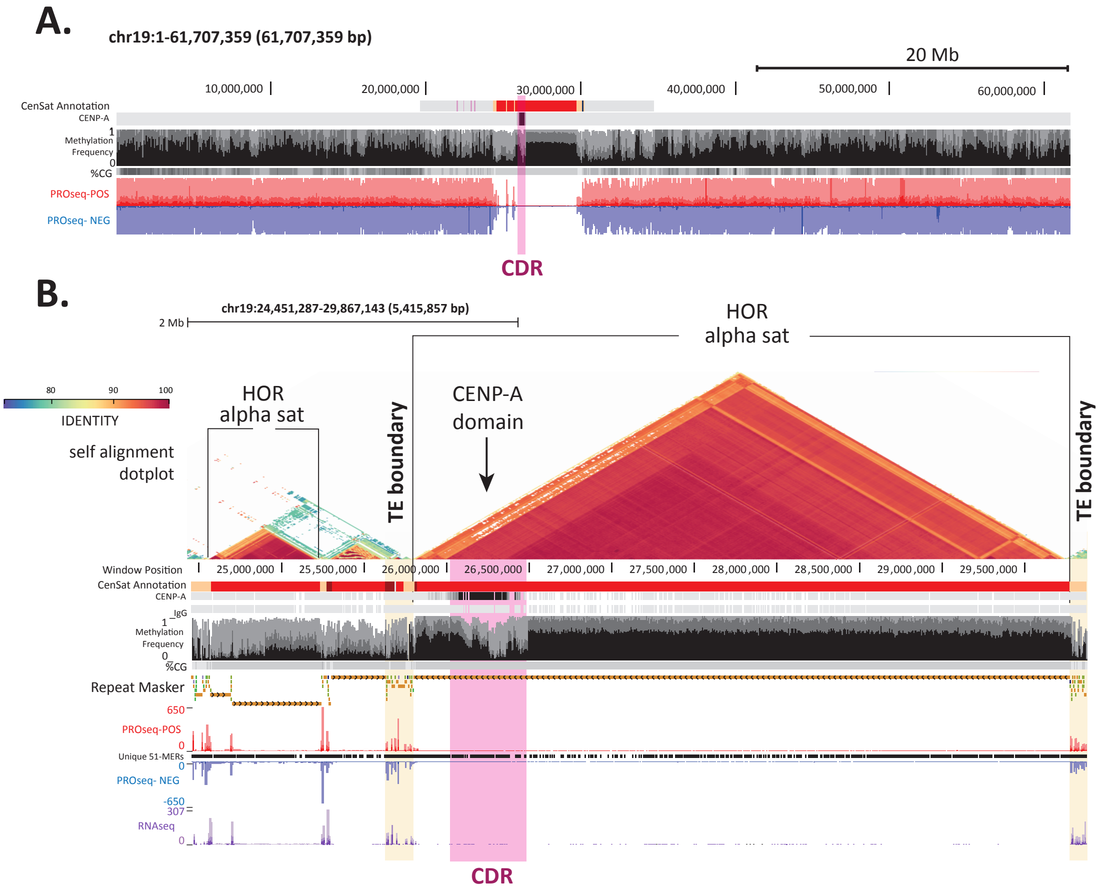


***Fig. S42. Sites of engaged RNA polymerase across Chromosome 19. (A)*** *Chromosome 19 from telomere to telomere. From top to bottom, cenSAT track (7), CENP-A CUT&RUN (7), methylation frequency (24), %CG, positive (red) and negative PRO-seq signal (blue). The CDR (24)(centromere dip region) that coincides with CENP-A (7) is indicated in pink.* ***(B)*** *Zoom of the centromere region of Chromosome 19. Tracks from top to bottom: self-alignment dot plot (scale of % identity is shown), cenSAT track, CENP-A CUT&RUN, IgG, methylation frequency, repeat masker v2 track, PRO-seq mapped reads (red = positive, blue = negative), unique 51mer k-mers, and RNA-seq mapped reads (purple). The CDR is indicated in pink, the TE boundaries are in tan.*

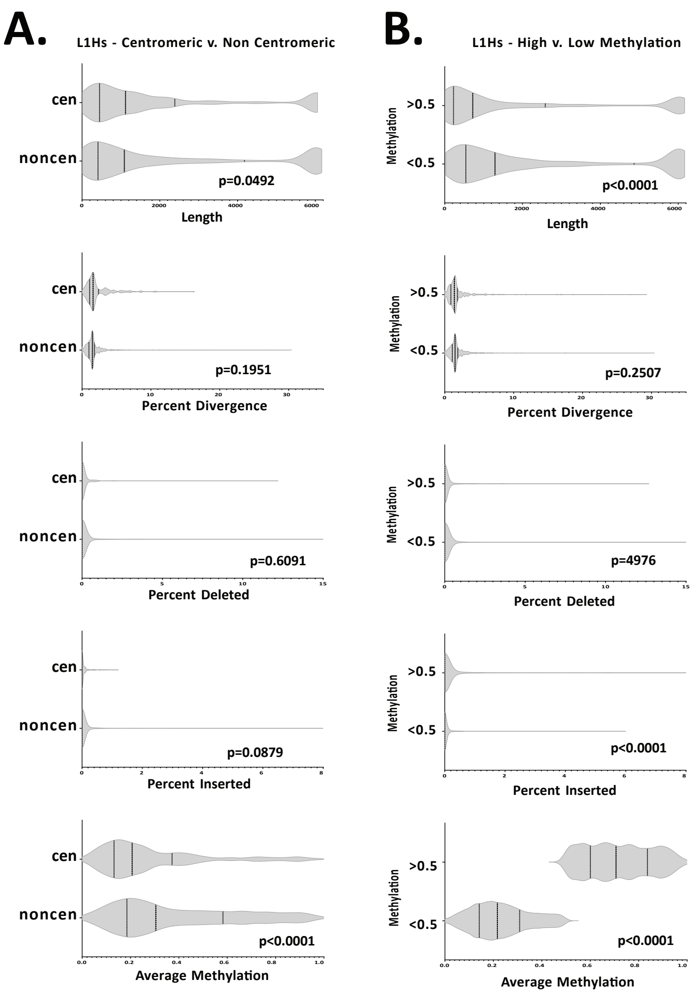
A.
L1Hs - Centromeric v. Non Centromeric
cen
noncen
p=0.0492
Length
p=0.1951
Percent Divergence
p=0.6091
Percent Deleted
p=0.0879
Percent Inserted
p<0.0001
Average Methylation
B.
L1Hs - High v. Low Methylation
Methylation
>0.5
<0.5
p<0.0001
Length
p=0.2507
Percent Divergence
p=4976
Percent Deleted
p<0.0001
Percent Inserted
p<0.0001
Average Methylation

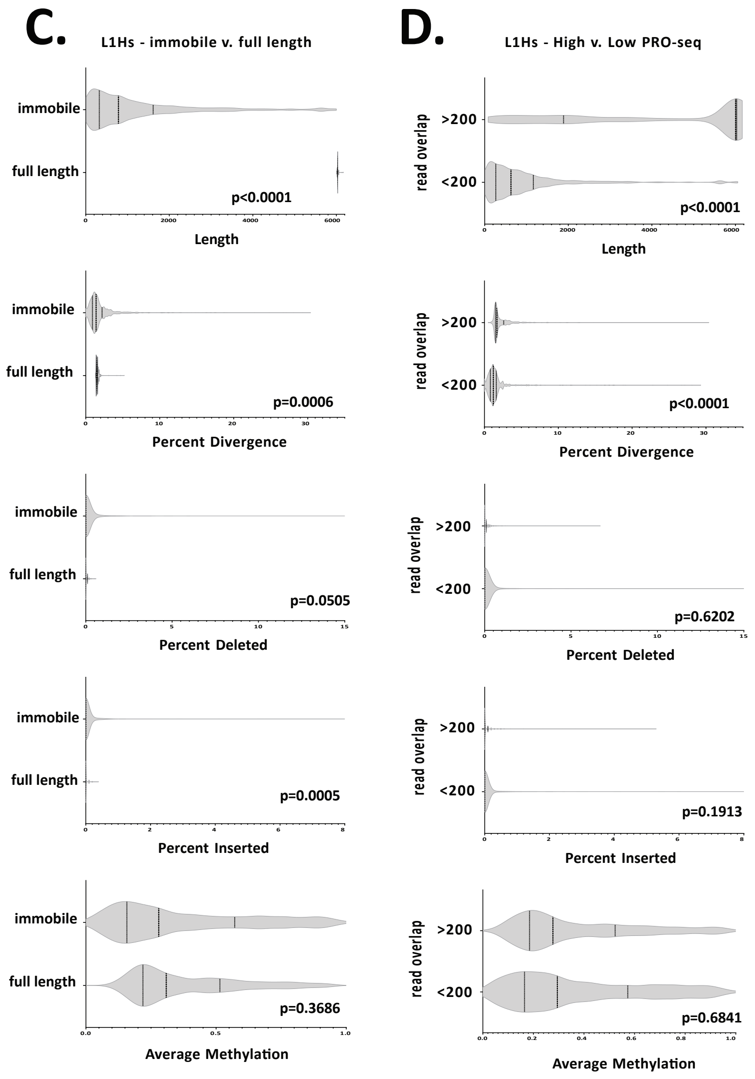


***Fig. S43.*** *Violin plots of L1HS elements show differences in length, divergence, deletions, insertions, and methylation of repeats found in the centromere (considered as centromere and centromere transitions as defined in (15)).* ***(A)****, with varying methylation patterns* ***(B)****, predicted to be "hot,"* ***(C)*** *and with varying PRO-seq expression levels* ***(D)****. Centromeric L1Hs are significantly shorter (p=0.0492) and have a lower average methylation (p<0.0001) than non-centromeric L1Hs. L1Hs with an average methylation < 0.5 are longer and have less insertions than L1Hs with an average methylation > 0.5 (p<0.0001). L1Hs predicted to be "hot" are longer, less diverged, and have less insertions than those predicted to be incapable of transposition (p=0.0005-<0.0001). Similarly, L1Hs with >200 overlapping PRO-seq reads are longer yet are more diverged from the consensus than L1Hs with <200 overlapping PRO-seq reads (p<0.0001).*

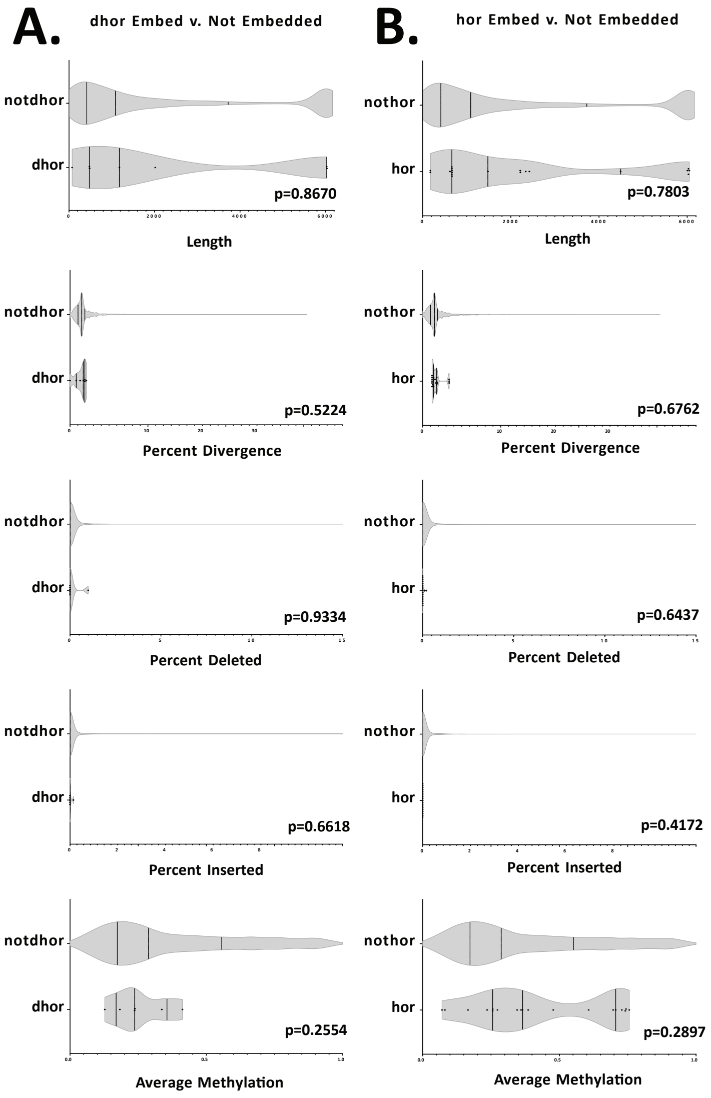
A.
dhor Embed v. Not Embedded
B.
hor Embed v. Not Embedded
notdhor
dhor
p=0.8670
Length
nothor
hor
p=0.7803
Length
notdhor
dhor
p=0.5224
Percent Divergence
nothor
hor
p=0.6762
Percent Divergence
notdhor
dhor
p=0.9334
Percent Deleted
nothor
hor
p=0.6437
Percent Deleted
notdhor
dhor
p=0.6618
Percent Inserted
nothor
hor
p=0.4172
Percent Inserted
notdhor
dhor
p=0.2554
Average Methylation
nothor
hor
p=0.2897
Average Methylation

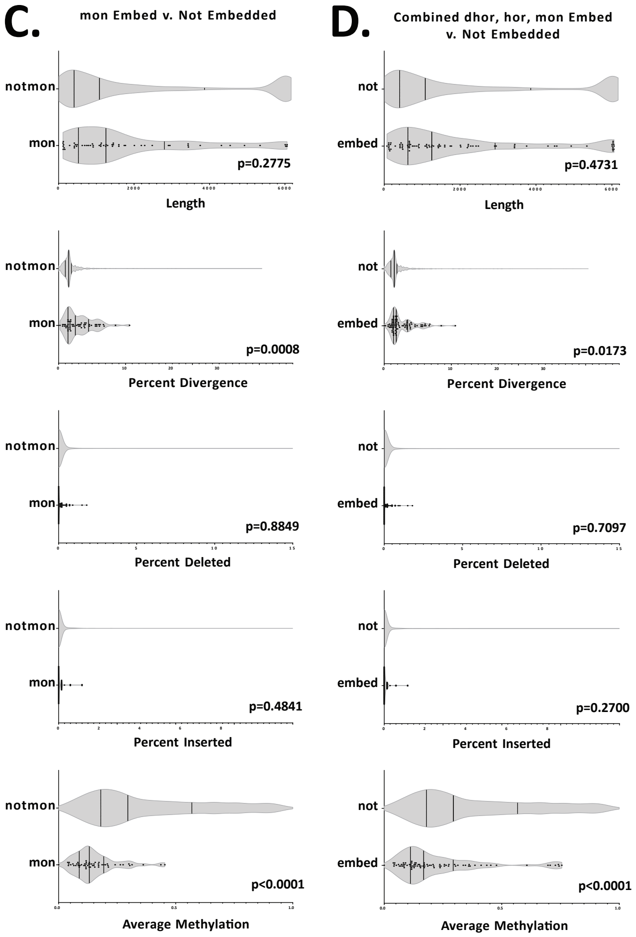
C.
mon Embed v. Not Embedded
notmon
mon
p=0.2775
Length
p=0.0008
Percent Divergence
p=0.8849
Percent Deleted
p=0.4841
Percent Inserted
p<0.0001
Average Methylation
D.
Combined dhor, hor, mon Embed v. Not Embedded
not
embed
p=0.4731
Length
p=0.0173
Percent Divergence
p=0.7097
Percent Deleted
p=0.2700
Percent Inserted
p<0.0001
Average Methylation

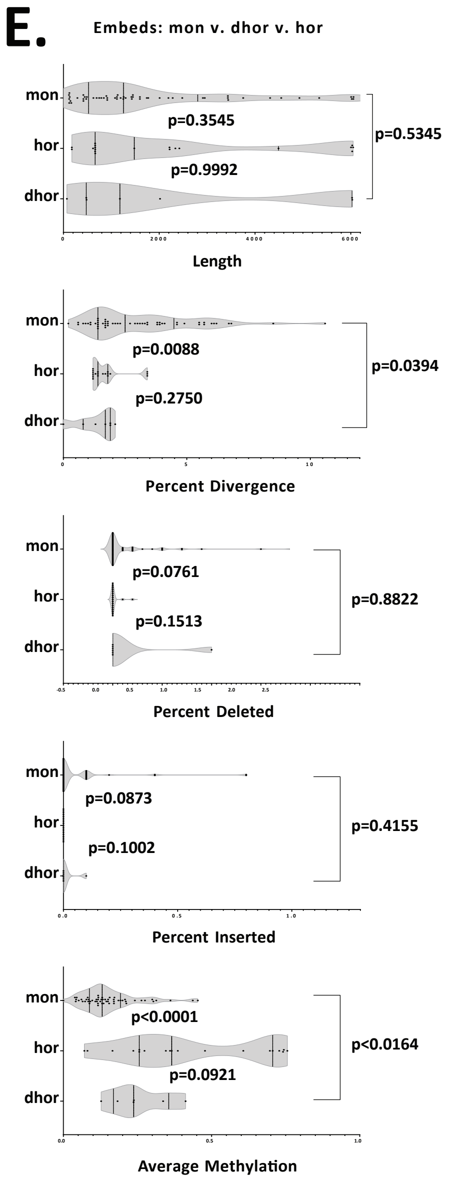


***Fig. S44.*** *Violin plots of L1Hs elements embedded in dHORs **(A)**, HORs **(B)**, and alpha satellite monomers **(C)** reveal few significant variations in length, divergence, deletions, insertions, and methylation of repeats. L1Hs embedded within alpha satellite monomers are both more diverged and less methylated than those not embedded in monomers (p=0.0008-<0.0001), accounting for the similar pattern in variation identified when comparing L1Hs embedded in these combined regions to unembedded repeats **(D)**. Comparing embeds in each classification **(E)** reveals that L1Hs embedded in monomers are more diverged and less methylated than those embedded in HORs (p=0.0088; p<0.0001, respectively) and dHORs (p=0.0394; p=0.0164, respectively), while those embedded in HORs and dHORs do not differ significantly.*

### 9. Repeat comparisons between CHM13v.10 and HG002

#### Methylation clustering

Methylation clustering was done by selecting all reads spanning a specific locus and using the mclust (v5.4.7) R package with the "VII" model to cluster methylation calls across the locus (cite package). Within mclust we specified G as being between 1 and 9 clusters. Positions with methylation calls that did not pass the threshold to be called methylated or unmethylated were assigned a value of 0.5. CpG density heatmaps were calculated by counting the total number of CpG sites per position relative to the repeat start and end and dividing by the total number of repeats in each group. Methylation single-read plots were generated in the ggplot2 R package using geom_rect() to plot individual reads with methylated CpGs as red and unmethylated CpGs as blue.

#### chrX liftOver analysis and repeat fasta comparison

Similar to the fasta sequence comparison of the unlifted CHM13 loci, the lifted chrX to HG002 coordinates were compared. A similarity score was assigned to each repeat based on crossmatch output as a percentage of the maximum score. Sequences with a score of greater than 90% and/or shorter than 50 bp were the threshold for concordant similarity or insufficient information for comparison, respectively. All other sequences were considered as potential polymorphic loci. These remaining 778 sequences of interest were filtered for length differences between the CHM13 and HG002 chrX liftOver coordinates. Simple repeats were not considered as part of this analysis. Differences that were further analyzed were loci 20 bp or greater if the CHM13 RM annotation was *Alu*, and 50 bp or greater for all other repeat types. 64 loci remained. For these 64 loci, the fasta sequence was extracted and subjected to RepeatMasker analysis.

| | TE subfamily | CHM13v1.0 | | | HG002 | | |
|---|---|---|---|---|---|---|---|
| | | All | FL | % FL | All | FL | % FL |
| Excluding the unassembled PAR1 region in HG002 | L1HS | 162 | 27 | 16.667% | 169 | 30 | 17.751% |
| | AluY | 5525 | 4508 | 81.593% | 5519 | 4516 | 81.826% |
| | SVA_E | 49 | 4 | 8.163% | 45 | 3 | 6.667% |
| | SVA_F | 43 | 10 | 23.256% | 43 | 12 | 27.907% |
| | HERVK | 23 | 0 | 0% | 55 | 1 | 1.818% |

| | TE subfamily | CHM13v1.0 | | | HG002 | | |
|---|---|---|---|---|---|---|---|
| | | All | FL | % FL | All | FL | % FL |
| Full assemblies | L1HS | 162 | 27 | 16.667% | 169 | 30 | 17.751% |
| | AluY | 5868 | 4757 | 81.067% | 5519 | 4516 | 81.826% |
| | SVA_E | 51 | 4 | 7.843% | 45 | 3 | 6.667% |
| | SVA_F | 45 | 10 | 22.222% | 43 | 12 | 27.907% |
| | HERVK | 23 | 0 | 0% | 55 | 1 | 1.818% |

***Table S24.*** *Full-length TE statistics for chrX in CHM13v1.0 and HG002. Since HG002 currently has a partially unassembled PAR1 region, the corresponding region in CHM13v1.0 (1-1929004) was excluded for the most accurate comparison to HG002 (top) but is also shown without this exclusion (bottom). Notably, the only TE family that is affected by this PAR1 region exclusion is AluY.*

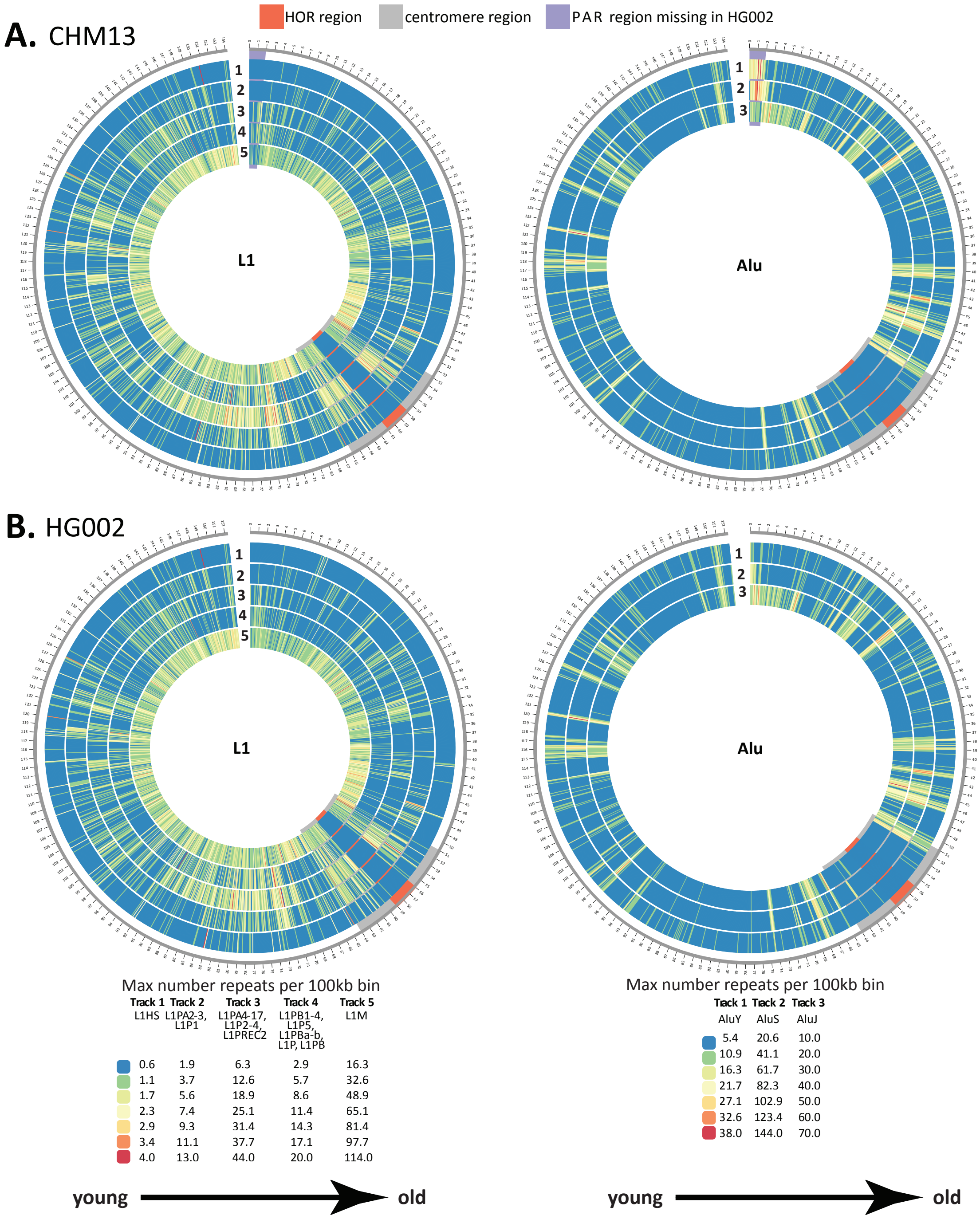


***Fig. S45. ChrX density of L1 and Alu subfamilies across CHM13 and HG002.*** *Subfamilies of L1s (LINE) and Alus (SINE) were grouped into general evolutionary age groups from youngest, including potentially mobile ones (L1HS, AluY), to oldest (L1M, AluJ). Counts of these grouped subfamilies were binned into Kbp windows across chrX in CHM13 and HG002 and are shown as Circos heatmaps. Centromere blocks (including centromere transition regions) are denoted by grey bars, HORs are denoted by orange bars, and the portion of the PAR1 that remains unassembled in HG002 is denoted in purple, all of which span all tracks. Tracks are numbered (1, 2, …) starting from the outer ring as indicated. In order to accurately compare density between the X's, each L1 or Alu subfamily track is shown on the same scale (i.e. CHM13 L1Hs*

*and HG002 L1Hs) with the scales for each subfamily group located below each set. At this resolution, Alus are depleted in centromeric region, possibly due to their enrichment in genic regions (59), but are found enriched in the PAR1 region with an increase AluY density closer to the telomere. In contrast, L1s have some of the highest density peaks at and around the centromere.*

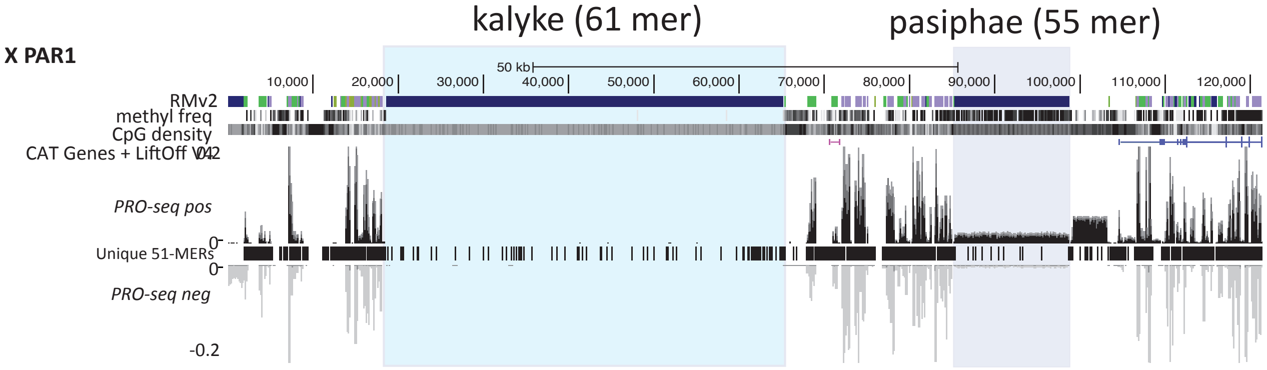


***Fig. S46. Pseudoautosomal region (PAR) of the CHM13 X chromosome contains new repeat annotations.*** *Included in these annotations are two tandem arrays, kalyke (61bp unit) and pasiphae (55bp unit). Kalyke has no detectable methylation while pasiphae has high levels of methylation coincident with some transcription (as detected by PRO-seq signal), indicating large arrays are differentially regulated on the X chromosome.*
